# Genome modeling and design across all domains of life with Evo 2

**DOI:** 10.1101/2025.02.18.638918

**Authors:** Garyk Brixi, Matthew G. Durrant, Jerome Ku, Michael Poli, Greg Brockman, Daniel Chang, Gabriel A. Gonzalez, Samuel H. King, David B. Li, Aditi T. Merchant, Mohsen Naghipourfar, Eric Nguyen, Chiara Ricci-Tam, David W. Romero, Gwanggyu Sun, Ali Taghibakshi, Anton Vorontsov, Brandon Yang, Myra Deng, Liv Gorton, Nam Nguyen, Nicholas K. Wang, Etowah Adams, Stephen A. Baccus, Steven Dillmann, Stefano Ermon, Daniel Guo, Rajesh Ilango, Ken Janik, Amy X. Lu, Reshma Mehta, Mohammad R.K. Mofrad, Madelena Y. Ng, Jaspreet Pannu, Christopher Ré, Jonathan C. Schmok, John St. John, Jeremy Sullivan, Kevin Zhu, Greg Zynda, Daniel Balsam, Patrick Collison, Anthony B. Costa, Tina Hernandez-Boussard, Eric Ho, Ming-Yu Liu, Thomas McGrath, Kimberly Powell, Dave P. Burke, Hani Goodarzi, Patrick D. Hsu, Brian L. Hie

## Abstract

All of life encodes information with DNA. While tools for sequencing, synthesis, and editing of genomic code have transformed biological research, intelligently composing new biological systems would also require a deep understanding of the immense complexity encoded by genomes. We introduce Evo 2, a biological foundation model trained on 9.3 trillion DNA base pairs from a highly curated genomic atlas spanning all domains of life. We train Evo 2 with 7B and 40B parameters to have an unprecedented 1 million token context window with single-nucleotide resolution. Evo 2 learns from DNA sequence alone to accurately predict the functional impacts of genetic variation—from noncoding pathogenic mutations to clinically significant *BRCA1* variants—without task-specific finetuning. Applying mechanistic interpretability analyses, we reveal that Evo 2 autonomously learns a breadth of biological features, including exon–intron boundaries, transcription factor binding sites, protein structural elements, and prophage genomic regions. Beyond its predictive capabilities, Evo 2 generates mitochondrial, prokaryotic, and eukaryotic sequences at genome scale with greater naturalness and coherence than previous methods. Guiding Evo 2 via inference-time search enables controllable generation of epigenomic structure, for which we demonstrate the first inference-time scaling results in biology. We make Evo 2 fully open, including model parameters, training code, inference code, and the OpenGenome2 dataset, to accelerate the exploration and design of biological complexity.

## 1. Introduction

Biological research scales from molecules to systems to organisms, seeking to understand and design functional components across all domains of life (Darwin, 1859; Mendel, 1866; Dobzhansky, 1951). Creating a machine to design functions across the diversity of life would require it to learn a deep, generalist representation of biological complexity. While this complexity surpasses straightforward human intuition, advances in artificial intelligence offer a universal framework that leverages data and compute at scale to uncover higher-order patterns (Vaswani et al., 2017; Kaplan et al., 2020). We reasoned that training a model with these capabilities would require data spanning the full spectrum of biological diversity to discover emergent properties similar to those found in other fields (Radford et al., 2019).

All domains of life express complex functions from DNA sequences (Watson and Crick, 1953; Nirenberg and Matthaei, 1961), yet genomic content and length vary dramatically across organisms. Prokaryotic genomes maintain relatively simple organization (Jacob and Monod, 1961; Overbeek et al., 1999), while eukaryotic evolution has produced intricate genomic architectures characterized by extensive noncoding regions, alternative splicing patterns, and multiple layers of epigenomic control (Chow et al., 1977; Brownell et al., 1996). These features underpin the emergence of multicellularity, sophisticated traits, and intelligent behaviors that are unique to eukaryotic life (Szathmáry and Smith, 1995).

We previously demonstrated that machine learning models trained on prokaryotic genomic sequences can model the function of DNA, RNA, and proteins, as well as their interactions that create complex molecular machines (Nguyen et al., 2024a; Merchant et al., 2024). However, extending this sequence modeling paradigm to eukaryotic genomes would require advances in data curation, model architecture, training and inference infrastructure, and inference-time compute to address the scale and complexity of eukaryotic genomes.

Here we present Evo 2, a biological foundation model that is trained on a representative snapshot of genomes spanning all observed evolution. Emphasizing generalist capabilities over task-specific optimization, Evo 2 achieves robust prediction and generation performance from molecular to genome scale and across all domains of life. We trained two versions of Evo 2 at 7B and 40B parameters, leveraging over 9.3T tokens at single-nucleotide resolution. These models were trained with a context window up to 1M tokens and demonstrate effective retrieval across the full context. To enable the research community, we release, to our knowledge, the largest-scale fully open language model to date, including open-source training code, inference code, model parameters, and the OpenGenome2 training data.

Evo 2 exhibits strong performance across biological sequence tasks. Building upon our previous work (Nguyen et al., 2024a), Evo 2 learns how mutations affect protein, RNA, and organismal fitness, while now generalizing beyond prokaryotes to include humans, plants, yeast, and other eukaryotes. Remarkably, without any variant-specific training, architectural optimization, or multiple sequence alignments, Evo 2 is the first language model capable of scoring the impact of all variant types on pathogenicity and splicing, achieving accurate and state-of-the-art performance in predicting the pathogenic effects of noncoding variation. Furthermore, a supervised model built on Evo 2 embeddings attains state-of-the-art performance on classifying *BRCA1* variants of unknown significance in breast cancer.

To elucidate the model’s learned concepts, we applied mechanistic interpretability techniques that decompose large language model representations into understandable components (Cunningham et al., 2023; Bricken et al., 2023). Using sparse autoencoders (SAEs), we identified a diverse set of features corresponding to key biological signatures, including intron and exon boundaries, transcription factor motifs, and protein structure characteristics. These feature-based annotations can also be leveraged for discovery tasks, such as identifying prophage regions and mobile genetic elements.

Evo 2 can also leverage its unique representation of biological complexity to generate new genomic sequences. We first demonstrate unconstrained generation of genome- and chromosome-scale sequences with improved naturalness compared to previous genomic language models. This includes the ability to generate complete mitochondrial genomes, minimal bacterial genomes, and entire yeast chromosomes. We also demonstrate how inference-time search can guide generation with Evo 2 to successfully achieve complex design tasks. In particular, we demonstrate controllable generation by using models of epigenomic state to design novel DNA sequences for which we can specify the location and length of chromatin-accessible regions, allowing us to write simple Morse code messages into our epigenomic designs. In doing so, we demonstrate the first inference-time scaling results for biological language modeling, extending our previous work that showed

Evo 2 and future iterations of the DNA foundation modeling paradigm represent the first steps toward generative biology for genomic and epigenomic design. This computational ability, combined with our recent experimental advances in large-scale programmable DNA manipulation (Durrant et al., 2024), may enable the direct programming of diverse synthetic life. Furthermore, combined with application-specific scoring functions to provide inference-time guidance, Evo 2 enables the design of complex biological architecture beyond DNA alone.

## 2. Results

### 2.1. Evo 2 model architecture, training procedure, and data

Evo 2 represents a major advance in genomic language models, scaling to 40 billion parameters and handling sequences of up to 1 million base pairs in length. Evo 2 was trained on genetic sequences from all domains of life and is useful for predictive and generative tasks across multiple scales of complexity (**Figure 1A**). We release OpenGenome2, a new dataset compiled from curated, non-redundant nucleotide sequence data, totaling over 8.8 trillion nucleotides from bacteria, archaea, eukarya, and bacteriophage (**Figures 1B** and **S1A**).

**Figure 1.**
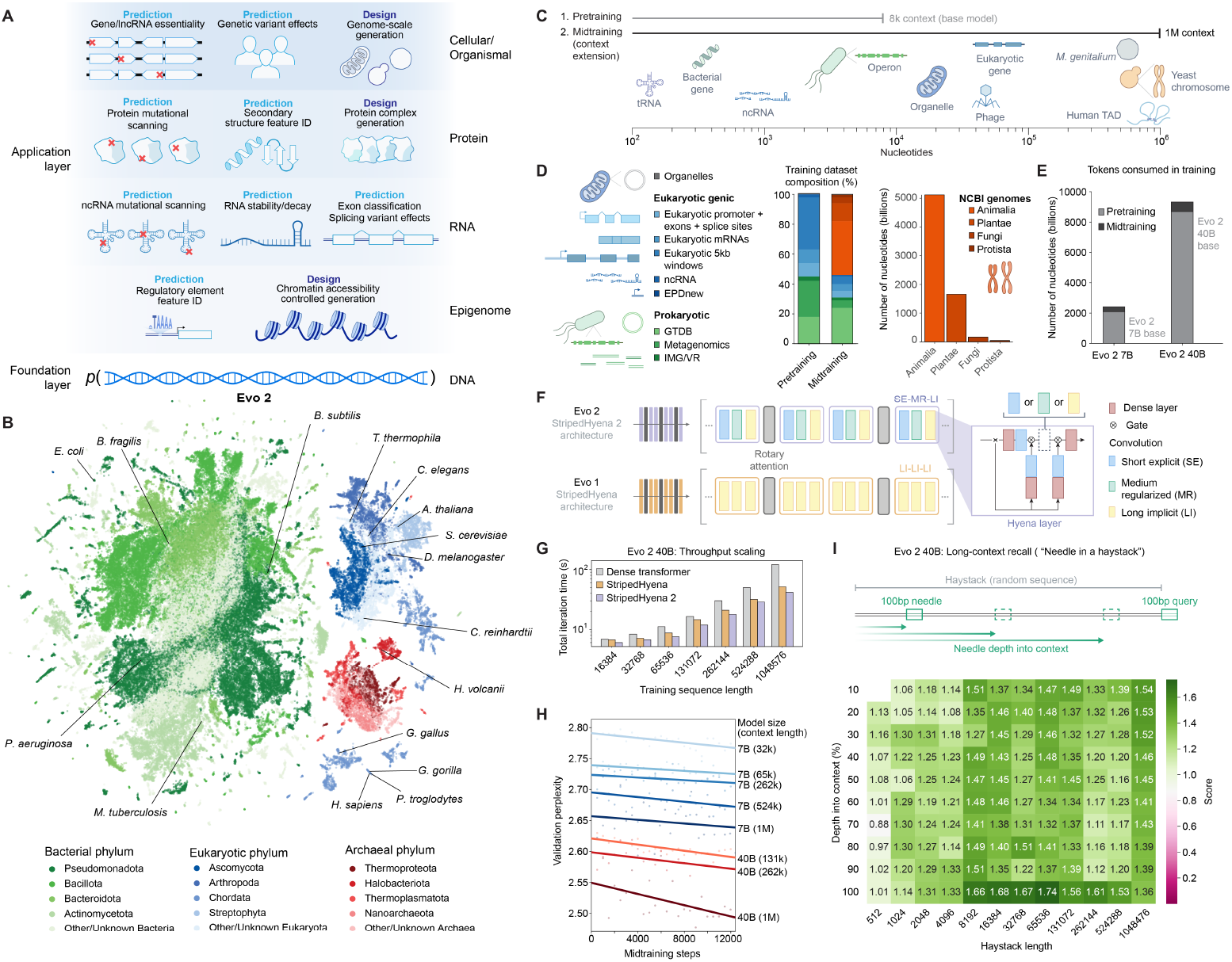
Overview of model architecture, training procedure, datasets, and evaluations for Evo 2. (**A**) Evo 2 models DNA sequence and enables applications across the central dogma, spanning molecular and cellular scales. (**B**) Evo 2 is trained on data encompassing trillions of nucleotide sequences from all domains of life. Each UMAP point indicates a single genome. (**C**) A two-phase training strategy optimizes model performance while expanding up to 1 million base pairs to capture wide ranging biological patterns. (**D**) Novel data augmentation and weighting approaches prioritize functional genetic elements during pretraining and long-sequence composition during midtraining. (**E**) The number of tokens used to train Evo 2 40B and 7B, split into the short phase pretraining and the long context midtraining. (**F**) Schematic of the new multi-hybrid StripedHyena 2 architecture, showing the efficient block layout of short explicit (SE), medium regularized (MR), and long implicit (LI) hyena operators. (**G**) Comparison of iteration time at 1024 GPU, 40B scale between StripedHyena 2, StripedHyena 1 and Transformers, showing improved throughput. (**H**) Validation perplexity of Evo 2 midtraining comparing the model size and context length, showing benefits with scale and increasing context length. (**I**) A modified needle-in-a-haystack task evaluates Evo 2’s long context recall ability up to 1 million sequence length, showing the model performs effective recall at 1 million token context. the first scaling laws for DNA sequence pretraining.

We trained two versions of Evo 2: a smaller version at 7B parameters trained on 2.4 trillion tokens and a full version at 40B parameters trained on 9.3 trillion tokens. Both Evo 2 7B and 40B are trained in two phases to capture biological length scales from molecular to organismal (**Figure 1C**). Our first stage of pretraining uses a context length of 8,192 tokens, with data weighting focused on genic windows to learn functional genetic elements (**Appendix B.2**), followed by a multi-stage midtraining phase over which we extend Evo 2’s context length to 1 million tokens to learn the relationships between elements across long genomic distances (**Figures 1C,D**). This matches best practice among large language models for natural language, where initial pretraining at shorter context lengths improves both efficiency and overall model quality (Gao et al., 2024b; Dubey et al., 2024; Liu et al., 2024). As in Evo 1, we excluded genomic sequences from viruses that infect eukaryotic hosts from the training data. We verified that these data exclusions led to high perplexity on genomic sequences from eukaryotic viruses (**Figure S2A**), indicating poor language modeling performance in this domain.

Evo 2 uses StripedHyena 2, the first convolutional multi-hybrid architecture (Ku et al., 2025). Multi-hybrids are a new class of model architectures designed explicitly to leverage the synergy between multiple different types of operators, arranged in a striped pattern. In particular, StripedHyena 2 relies on a combination of three different variants of input-dependent convolution operators (Poli et al., 2023) and attention (**Figure 1F**), improving efficiency of training at scale on both short and long sequences. StripedHyena 2 provides substantially higher throughput (at 40 billion parameters, up to 1.3 × speedup at 16 thousand context length and 3× speedup at 1 million context length) than highly optimized Transformer (Vaswani et al., 2017) baselines and previous generation hybrid models based on recurrences or long convolutions, such as StripedHyena 1 (Poli et al., 2024) (**Figure 1G**). StripedHyena 2 also improves loss scaling on DNA against both Transformers and StripedHyena 1 (**Figure S1D**).

We compare different context extension methods for StripedHyena 2 and find methods using rotary embeddings can be applied to effectively extend the context length, similarly to StripedHyena 1 in Evo 1 (Ku et al., 2025). This enables us to train up to 1 million base pairs in context length through a multi-stage extension phase, which shows improvements in loss with both model scale and longer context (**Figure 1H)**. With a synthetic long-context evaluation, we demonstrate that the model can implement recall to retrieve a 100 base pair needle from different positions in a 1 million base pair long “haystack” of random DNA sequence (**Figure 1I** and **S1C**).

To promote open science and facilitate community development, we have released Evo 2’s model parameters, training code, inference code, and training data freely available under an open-source license (**Discussion**). This makes Evo 2 one of the largest-scale fully open AI models, not just in biology but also compared to natural language models based on the Transformer architecture (**Table 1**).

**Table 1.** Comparison of training FLOPS across flagship language and biology models, showing Evo 2 as the largest fully open model. FLOPS are estimated without accounting for mixed-precision or pretraining context length.

| Model | Model Size | Tokens | FLOPS est. |
| --- | --- | --- | --- |
| **Fully open: data, infrastructure, weights** |  |  |  |
| Evo 2 40B | 40.3B | 9.3T | $2.25 \times 10^{24}$ |
| Pythia Large | 12B | 300B | $2.16 \times 10^{22}$ |
| OLMo 2 Large | 13B | 5.6T | $4.37 \times 10^{23}$ |
| **Open data, weights** |  |  |  |
| Evo 1 | 7B | 300B | $1.26 \times 10^{22}$ |
| Falcon 180B | 180B | 3.5T | $3.78 \times 10^{24}$ |
| StarCoder 2 15B | 15B | 4.3T | $3.87 \times 10^{23}$ |
| **Open weights only** |  |  |  |
| DeepSeek V3 | 37B (671B total) | 14.8T | $3.28 \times 10^{24}$ |
| Llama 3.1 Large | 405B | 15T | $3.64 \times 10^{25}$ |
| **Closed** |  |  |  |
| ESM 3 Large | 98B | 771B | $1.07 \times 10^{24}$ |
| xTrimo Large | 100B | 1T | $6.00 \times 10^{23}$ |

### 2.2. Evo 2 predicts mutational effects on protein, RNA, and organismal fitness across all domains of life

By learning the likelihood of sequences across vast evolutionary training datasets, biological sequence models can learn how mutational effects correlate with biological functions without any task-specific finetuning or supervision. This is referred to as zero-shot prediction. Previously, however, effective zero-shot mutational effect prediction has been shown for language models trained only on protein sequences (Meier et al., 2021; Notin et al., 2023) or genome language models trained only on prokaryotic sequences (Nguyen et al., 2024a). Given that Evo 2 learns a sequence likelihood landscape across all three modalities of the central dogma (DNA, RNA, protein) and all three domains of life (prokaryota, archaea, eukaryota), we sought to assess whether Evo 2 enables mutational effect prediction across all of these modalities and organisms (**Figure 2A**).

**Figure 2.**
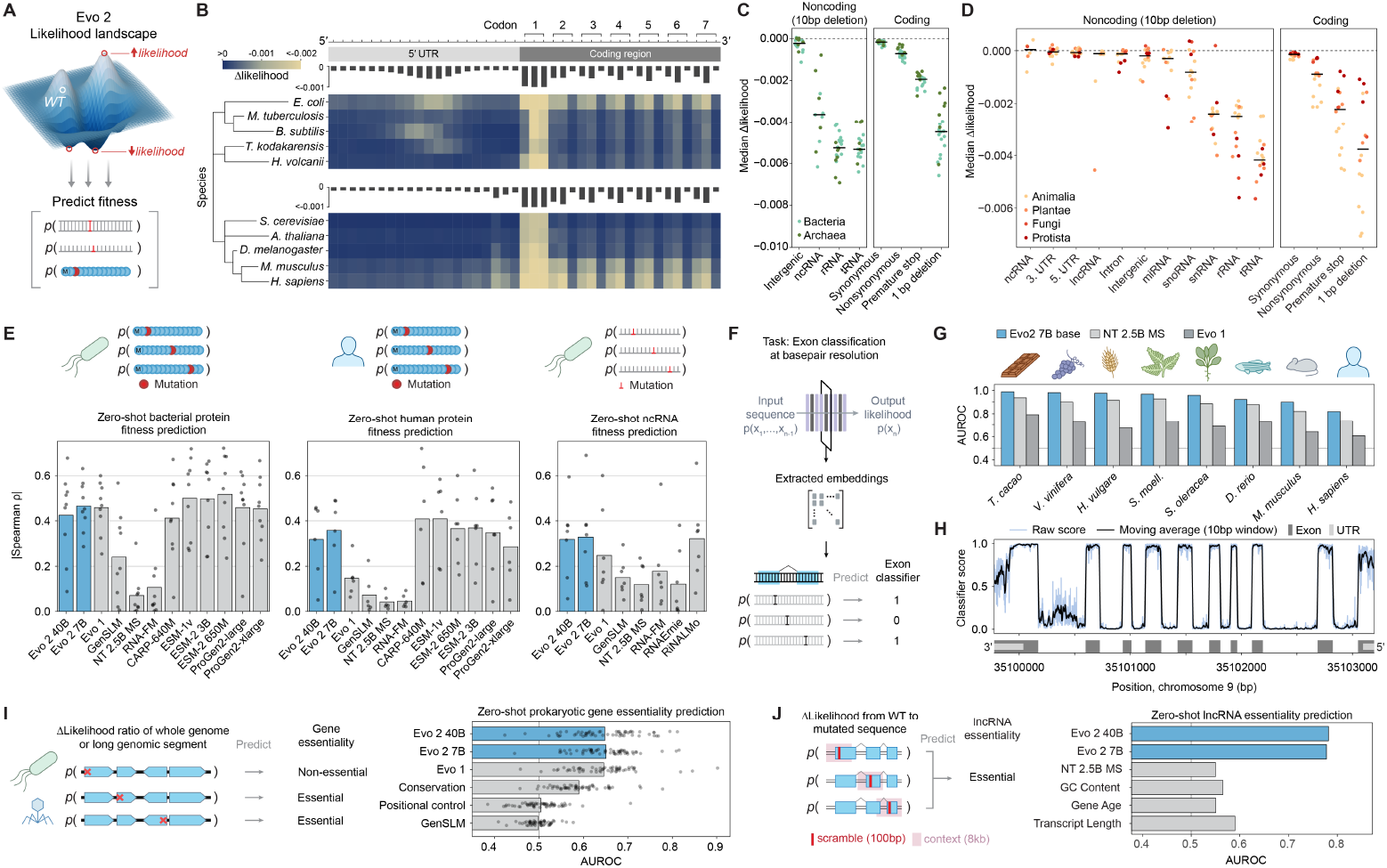
Evo 2 predicts mutational effects on protein, RNA, and organismal fitness across all domains of life. (**A**) We used Evo 2’s zero-shot likelihoods to predict the effects of DNA, RNA, or protein mutations on molecular function or organismal fitness. (**B**) We used the change in Evo 2’s sequence likelihood caused by different mutations along gene start sites for various model species across the tree of life. Evo 2 predicts mutations to be unlikely in the start codons of protein-coding genes, the first two bases of each codon of the coding region, and the ribosome-binding sites of the 5′ UTR. See **Figure S3A** for an additional analyses. (**C, D**) For different prokaryotic (**C**) and eukaryotic (**D**) sequences, we scored the likelihood of different types of mutations in different genomic elements using Evo 2 7B. Results align with known biology such that regions under stronger evolutionary constraint are also more sensitive to mutational likelihoods under Evo 2. Scatter represents median change in likelihood from WT to mutant sequence per species, colored by domain (**C**) or kingdom (**D**); horizontal line indicates median of the scatter distribution. (**E**) We used a wide range of diverse deep mutation scanning (DMS) assays to assess the Spearman correlation of zero-shot likelihoods from models with experimental assays. (**F**) Schematic of our single-nucleotide resolution exon classifier based on embeddings from Evo 2. (**G**) We compared single-nucleotide resolution exon classifiers trained on embeddings from Evo 2, Nucleotide Transformer (NT), or Evo 1 across eight held-out species, with performance measured by area under the receiver operating characteristic curve (AUROC) in classifying exonic base pairs. (**H**) Genome browser track showing predictions from the Evo 2 embedding-based exon classifier scanned across the human *STOML2* locus, where the vertical axis is the predicted classifier score and the horizontal axis is genome position. (**I**) We used the mutational likelihood of premature stop codon insertions (as a genetic perturbation) to use Evo 2 to predict genes as essential or nonessential, as determined by experimental gene essentiality assays across bacterial and phage species (shown as overlaid scatter). (**J**) We used the mutational likelihood of scrambled sequence (as a genetic perturbation) to use Evo 2 to predict human lncRNAs as essential (*N* = 46) or nonessential (*N* = 5,417) in all tested cell lines, as experimentally determined using lncRNA cellular essentiality screens.

Coding sequences across all domains of life follow a fundamental structure: they begin with a start codon (Marcker and Sanger, 1964), end with a termination codon (Brenner et al., 1965), and are translated using a triplet codon code that defines the reading frame (Nirenberg and Matthaei, 1961). To assess whether Evo 2 captures these core biological principles, we first evaluated how single nucleotide variants (SNVs) impact Evo 2 likelihoods in the genomic sequences around the start codons of protein-coding genes. We introduced these mutations into the wildtype sequence at each position and calculated how the Evo 2 predicted likelihoods changed across thousands of such loci (**Figures 2B** and **S3A**). We observed strong changes in the likelihood for mutations within the start codons in both prokaryotic and eukaryotic species. This was followed by a three-base periodicity pattern reflecting the triplet codons, with changes at the wobble positions showing lower impact on likelihood. For both prokaryotic and eukaryotic genomes, we observed a pattern upstream of the CDS that was consistent with conserved ribosome binding sites (Shine and Dalgarno, 1974; Kozak, 1989). We also observed similar patterns for SNVs around stop codons (**Figure S3B**). These results confirmed that the model had learned these fundamental genetic features across the domains of life, despite not having seen any annotations of these sequences in its training data.

Beyond coding sequences, our current understanding of genomes presumes that specific genetic alterations should result in differential phenotypic consequences. For example, non-synonymous mutations should be more disruptive than synonymous ones, frameshifts and premature stop codons should be maximally disruptive, and in essential noncoding elements, deletions should have greater consequences than those in intergenic regions. By measuring the impact of a variety of mutations across both noncoding and coding sequences, we sought to evaluate whether Evo 2 likelihoods capture these known priors (**Figures 2C,D**). Across 20 prokaryotic species and 16 eukaryotic species, we observed changes in model likelihoods consistent with known biological constraints. Within coding sequences, non-synonymous variations, premature stop codons, and frameshift mutations caused much larger changes in likelihood than synonymous mutations. In noncoding regions, deletions in tRNAs and rRNAs had significantly greater effects than deletions in intergenic and other noncoding loci, reflecting the known essential roles of these RNAs. Interestingly, the 40B model exhibited higher sensitivity to deletions in miRNA and snoRNA sequences compared to the 7B model (**Figure S3C**), suggesting that larger models likely capture more nuanced regulatory features. Additionally, Evo 2 performance exceeds that of other DNA language models on three recently published zero-shot evaluation tasks of human noncoding regulatory sequences, demonstrating progress in modeling these notoriously “fuzzy” DNA elements (**Figure S3D**) (Patel et al., 2024).

Recognizing that our training data contained genomes with distinct genetic codes, we also tested how different premature stop codons impacted species that differed in their stop codon usage (**Figure S3E**). We found that the model clearly learned the difference between the standard code (stop codons TAA, TAG, and TGA), the mycoplasma code (Code 4, stop codons TAA and TAG), and the ciliate code (Code 6, stop codon TGA). Interestingly, 4 to 8 kb context windows around the premature stop codons were necessary to correctly identify the ciliate stop codon code, demonstrating the value of longer context windows for this task (**Figure S3F**).

While Evo 2 predictions reflect the expected importance of different genetic alterations, a key question is whether these likelihoods correspond to empirically measured functional effects. Deep mutational scanning (DMS) provides a systematic framework for measuring the fitness impact of mutations across a wide range of proteins and noncoding RNAs (ncRNAs). By comparing Evo 2’s zero-shot likelihoods to experimental measurements from DMS, we found that Evo 2’s sequence likelihoods correlate with diverse definitions of fitness for both prokaryotic and eukaryotic protein and ncRNA molecules (**Figure 2E**). Notably, Evo 2 is competitive with state-of-the-art autoregressive protein language models in predicting the fitness of both bacterial and human proteins, and achieves state-of-the-art performance on noncoding RNA fitness prediction.

We also tested the ability for Evo 2 to predict mutational effects in protein sequences from viruses that infect human hosts. We found no correlation between Evo 2 likelihood and viral protein fitness (**Figure S2B**). This is consistent with our data exclusions having the intended effect of weakening both language modeling performance (**Figure S2A**) and downstream prediction tasks.

Additionally, since Evo 2 is pre-trained on both DNA and RNA sequences, it has learned sequence features that contribute to molecular fitness across both domains. Therefore, we expect that the likelihood scores assigned to RNA sequences by the model would be associated with their overall stability. To test this hypothesis, we conducted a zero-shot evaluation by comparing the model-derived likelihoods for human mRNAs against their average decay rates measured in an mRNA stability dataset. Consistent with an expected negative association between model scores and mRNA decay, we found that among the evaluated models, only Evo 2 likelihoods were found to negatively correlate with the mRNA decay rates, with the 40B model showing a stronger negative correlation than the 7B (**Figure S3G**).

Since Evo 2 learns from eukaryotic genomes, we assessed its ability to capture the canonical exonic and intronic patterns that represent some of the fundamental building blocks of the genetic code. To do this, we leveraged Evo 2 embeddings to develop a single-nucleotide resolution classifier of exon labels from reference annotations (**Methods**; **Figure 2F**). After optimizing small-scale classifiers on embeddings from each model, we evaluated their performance across eight diverse species held out from classifier training. The Evo 2 embedding-based classifier achieved superior performance to models trained on embeddings from Nucleotide Transformer and Evo 1. Our classifier also had high accuracy on this task, with areas under the receiver operating characteristic curve (AUROCs) ranging from 0.82-0.99 across species (**Figure 2G**). As an example demonstration, we scanned the classifier across the human protein-coding gene locus of the *STOML2* gene. The classifier’s probabilities showed strong correspondence with annotated exon locations, with distinct transitions at exon-intron boundaries (**Figure 2H**). Using Evo 2 sequence embeddings, potentially combined with bioinformatic approaches, may aid functional annotation of genetic components in poorly characterized genomes.

Beyond molecular or gene-level prediction tasks, we previously showed that Evo 1 can predict mutational effects on whole organism fitness in prokaryotes. Using zero-shot likelihoods to score the effects of premature stop codon insertions into bacterial and phage genomes, we found that the Evo 2 models matched the performance of Evo 1 in predicting gene essentiality across diverse essentiality studies (**Figure 2I**) (Zhang, 2004; Piya et al., 2023). These results indicate that Evo 2, even after expanding its scope to incorporate eukaryotic training data, has retained its predictive performance in prokaryotic tasks.

Expanding our analysis to whole organism fitness effects in eukaryotes, we evaluated Evo 2’s ability to predict noncoding gene essentiality in human cells. Using data from a recent long noncoding RNA (lncRNA) essentiality study (Liang et al., 2024), we found that both Evo 2 models substantially outperformed Nucleotide Transformer and other sequence-based metrics when assessing the impact of artificial disruptions—specifically, by scrambling 100 bp subsequences within the lncRNA gene (**Figure 2J**). This enhanced predictive performance was consistently observed across the five cell lines tested in the original study and was even more pronounced for lncRNAs deemed essential in multiple cell lines (**Figure S3H,I**), indicating that Evo 2 effectively captures the breadth of fitness contributions from functional lncRNAs.

In total, these results demonstrate that Evo 2 has captured a remarkable breadth of information across biological modalities and domains of life. Notably, the 7B and 40B models expand predictive capabilities without compromising the prokaryotic insights captured by Evo 1. The utility of both zero-shot likelihoods and simple classifiers trained on Evo 2 embeddings for a variety of predictive tasks across prokaryotic and eukaryotic genomes indicates that Evo 2 provides a strong foundation model for many downstream applications in computational biology.

### 2.3. Evo 2 enables accurate human clinical variant effect prediction

Variant effect prediction (VEP) evaluations have emerged both as a foundational application of and litmus test for genome language models (Benegas et al., 2025; Ji et al., 2021; Dalla-Torre et al., 2024), assessing their ability to capture functional constraints, predict pathogenicity, and generalize across diverse genomic contexts. Genome language models can perform variant effect predictions for both coding and noncoding DNA, zero-shot, by considering the predicted changes in sequence likelihoods.

We benchmarked the performance of Evo 2 against the annotations of experimental and clinical variants to assess its ability to predict biologically significant sequence variation. We first investigated the ability of Evo 2 to predict the pathogenic effects of human genome variants across diverse variant classes (**Figure 3A**). Specifically, we compared the zero-shot performance of both Evo 2 models with several other models, including state-of-the-art VEP models such as AlphaMissense (Cheng et al., 2023) and GPN-MSA (Benegas et al., 2025), in predicting pathogenic ClinVar variants (**Figure 3B, Table S7**). For single-nucleotide variants (SNVs) in coding regions, the 40B and 7B models ranked fourth and fifth, respectively, behind AlphaMissense, ESM-1b, and GPN-MSA. For coding variants that were not SNVs (e.g., insertions and deletions), which models such as AlphaMissense and GPN-MSA fail to evaluate, both Evo 2 models outperformed the other models in zero-shot classification. For noncoding variants, Evo 2 also surpassed other models for both SNVs and non-SNVs (**Figure 3C**).

**Figure 3.**
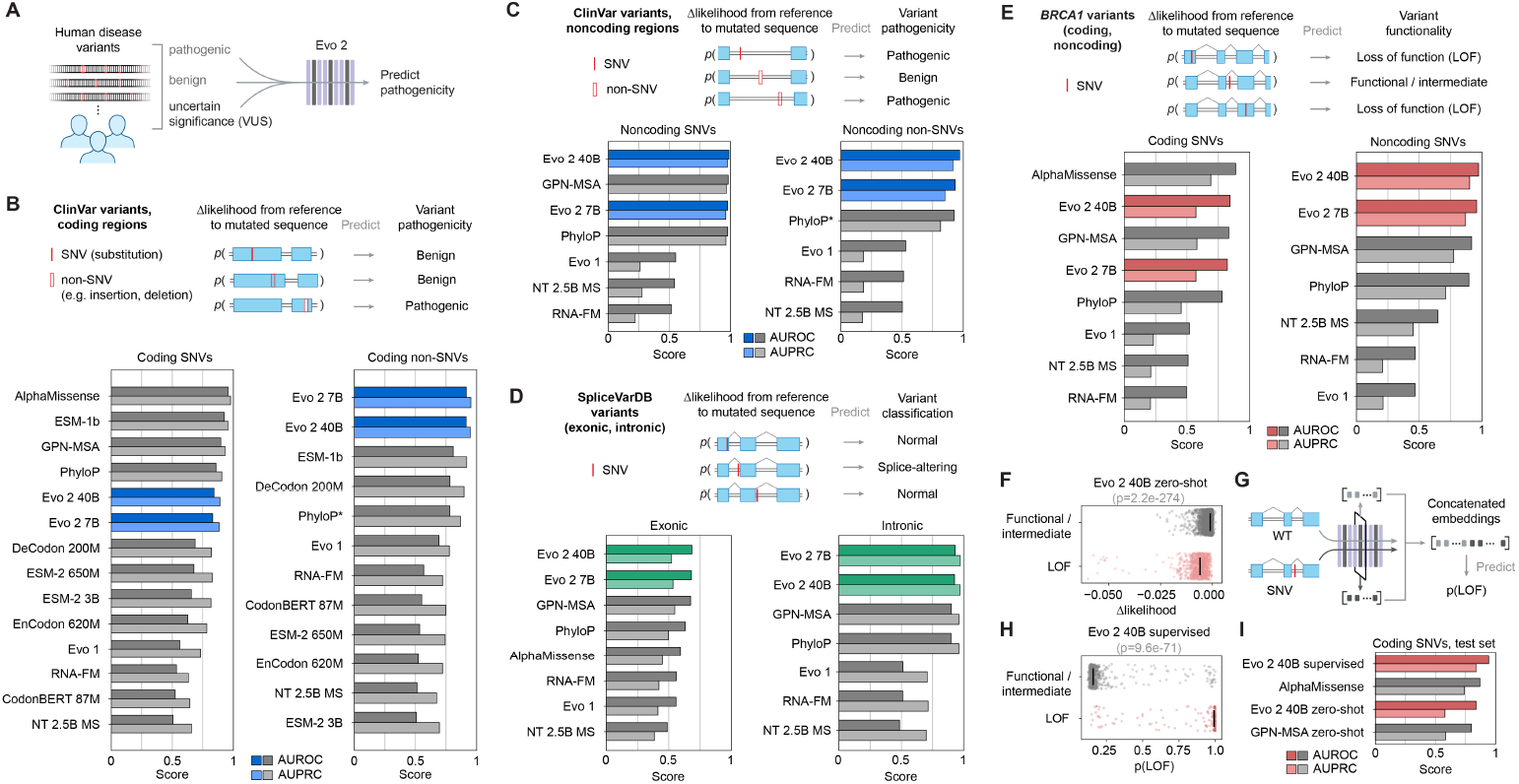
Evo 2 enables accurate human clinical variant effect prediction. (**A**) Overview of zero-shot variant effect prediction using Evo 2. Evo 2 assigns likelihood scores to human disease variants, distinguishing pathogenic and benign variants in both coding and noncoding regions. (**B, C**) Zero-shot evaluation of variant pathogenicity within the coding (**B**; *N* = 14,319 SNVs, *N* = 1,236 non-SNVs) and noncoding (**C**; *N* = 34,761 SNVs; *N* = 3,894 non-SNVs) regions. Shown are the AUROCs and AUPRCs for classifying pathogenic and benign variants, from ClinVar, across models. For non-SNV evaluations, we used a modified version of PhyloP (**Methods**). (**D**) Zero-shot evaluation on splice-altering variants in SpliceVarDB, split by exonic (*N* = 1,181) and intronic (*N* = 3,769) scoring. (**E**) Model predictions evaluated against *BRCA1* saturation mutagenesis data, comparing classification of loss-of-function (LOF) versus functional/intermediate variants in both coding (*N* = 2,077 SNVs) and noncoding (*N* = 1,125 SNVs) regions. (**F**) Evo 2 zero-shot likelihood scores plotted for LOF versus functional/intermediate variants (*N* = 3,893), demonstrating the model’s ability to separate these classes. *P*-value calculated by two-sided Wilcoxon rank sum test. (**G**) A schematic for supervised variant effect prediction classifiers: Evo 2 embeddings are extracted and concatenated to train a supervised classifier for *BRCA1* variant effect prediction. (**H**) Predictions of the supervised classifier on functional/intermediate variants compared with true LOF variants on the test set (*N* = 779), with prediction scores on the horizontal axis. *P*-value calculated by two-sided Wilcoxon rank sum test. (**I**) Comparing a supervised version of Evo 2 on the *BRCA1* test set against zero-shot baselines and AlphaMissense shows the value of using Evo 2 embeddings for predictive models.

Although ClinVar is a valuable benchmark, its coverage is biased toward well-studied variants that often have established clinical significance. To evaluate model performance on a broader and potentially more challenging set of functional noncoding variants, we turned to SpliceVarDB, a repository of experimentally validated splicing effects. For both exonic and intronic splice variant effect prediction, Evo 2 models achieved the highest zero-shot performance (**Figure 3D**). Together, these results highlight the competitive performance of Evo 2 in predicting the pathogenic effects of human coding SNVs against specialized models like AlphaMissense and GPN-MSA, while establishing a new state of the art for zero-shot scoring of non-SNV, noncoding, and splice-associated variants.

To further limit curation bias and incomplete experimental coverage of coding and noncoding variants, we focused on a dataset measuring functional consequences of variants across both coding and noncoding regions of the *BRCA1* gene (Findlay et al., 2018). Similar to our previous observations, zero-shot predictions with Evo 2 performed strongly for coding SNVs and set a new state-of-the-art for *BRCA1* noncoding SNVs. Importantly, it outperformed all other models when coding and noncoding variants were evaluated combined together (**Figures 3E** and **S4A**).

A recently released *BRCA2* variant dataset with experimental measurements allowed us to extend this analysis to *BRCA2* (Huang et al., 2025). Consistent with our findings for *BRCA1*, Evo 2 surpassed existing benchmarks from specialized models like GPN-MSA when predicting coding and noncoding variants together (**Figure S4B**). This strong performance on scoring both coding and noncoding variants for *BRCA1* and *BRCA2*, along with Evo 2’s robust results on non-SNV benchmarks, indicates that Evo 2 is a well-calibrated zero-shot predictor of a diverse range of functional human variants. This is particularly important given that many noncoding variants are routinely excluded from variant analysis reports. Notably, the only human genome in OpenGenome2 is the reference genome, and Evo 2 achieves this zero-shot performance without any human variant training, instead leveraging multi-species variation as a proxy for evolutionary constraints.

While zero-shot evaluation provides an initial measure of a model’s inherent ability to predict functional effects (**Figure 3F**), the model-derived embeddings can also be leveraged in supervised classifiers to systematically refine predictions by learning task-specific decision boundaries, potentially enhancing sensitivity and specificity. To demonstrate this, we evaluated a supervised deep neural network trained specifically on *BRCA1* variants to see whether it could outperform zero-shot methods, thereby illustrating the potential of Evo 2 as a foundation for improved variant classification models (**Figure 3G**). Because different layers of large language models encode different levels and types of abstraction, we systematically extracted sequence embeddings from each block of the Evo 2 40B model to determine which layer provided the most relevant information for variant classification (**Methods**; **Figures S4C,D**).

Our best-performing supervised model achieved a clear separation between loss-of-function variants and all other variants (**Figure 3H**), outperforming all benchmarks on the *BRCA1* coding SNV test set (AUROC = 0.94, AUPRC = 0.84) (**Figure 3I**). The model also excelled for all *BRCA1* SNVs in the test set (AUROC = 0.95, AUPRC = 0.86) (**Figure S4E**). These results further underscore how Evo 2 embeddings can be harnessed to train models aimed at more specialized tasks, including those of high clinical relevance.

Our evaluations across ClinVar, SpliceVarDB, and *BRCA1/2* variant datasets demonstrate that Evo 2 models achieve state-of-the-art performance for noncoding variants while remaining competitive on coding variants.

Their zero-shot capability highlights the strength of their learned sequence representations, and leveraging these representations in a supervised setting illustrates how they can serve as a powerful foundation for downstream variant effect prediction tasks. In sum, these findings confirm the versatility of Evo 2 as a genome-scale language model for both unsupervised and supervised applications, offering a robust framework for variant interpretation across diverse genomic contexts.

### 2.4. Feature interpretation in Evo 2 from molecular to genome scale

Evo 2 learns complex representations of genomic sequences without relying on explicit biological labels or annotations. Contrary to the common critique of large language models as black box systems, recent advances in mechanistic interpretability have demonstrated that sparse autoencoders (SAEs) can reveal latent dimensions that correspond to semantically meaningful features in natural language (Cunningham et al., 2023; Bricken et al., 2023; Templeton et al., 2024). To probe what Evo 2 is capturing, we trained SAEs on its representations, or neuron firing patterns, to decompose the model into sparse, high-dimensional representations in which individual latent dimensions often exhibit human-interpretable patterns (**Figures 4A**).

**Figure 4.**
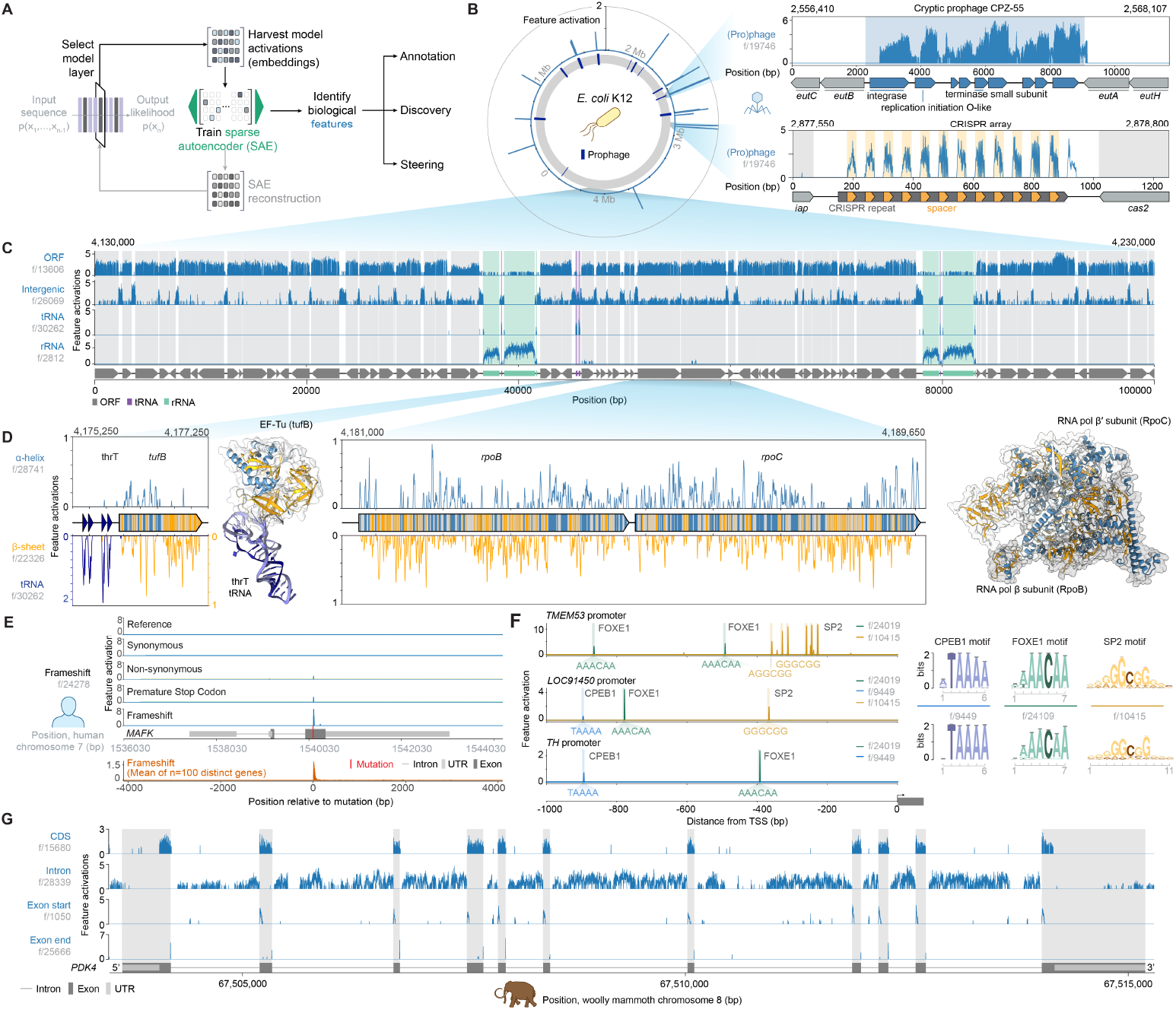
Mechanistic interpretability of Evo 2 reveals DNA, RNA, protein, and organism level features. (**A**) Sparse autoencoders (SAEs) were trained on Evo 2 to extract SAE features associated with interpretable biological function that can be used for annotation, discovery, and steering of sequence generations. (**B**) Phage-associated feature activates preferentially on RefSeq-annotated prophages (left and top right) in the *E. coli* K12 MG1655 genome and fires on phage-derived spacer sequences within CRISPR arrays (bottom right). (**C**) Activations of features associated with open reading frames (ORFs), intergenic loci, tRNAs, and rRNAs, in a 100 kb region in *E. coli* K12 MG1655. (**D**) Activations of features associated with α-helices, β-sheets, and tRNAs at an *E. coli* K12 MG1655 locus containing tuf B and a tRNA array ending with thrT (left) and the rpoB-rpoC locus (right). AlphaFold 3 (AF3) structure predictions with feature activations overlaid, of EF-Tu in complex with thrT tRNA (left) and of RpoB and RpoC in complex (right). (**E**) A feature in the human genome preferentially activates immediately after frameshift mutations over less deleterious mutation types. (**F**) Features activating on DNA motifs in the human genome that correspond to transcription factor-binding motifs. (**G**) Features associated with exons, introns, and their boundaries in the human genome can be used to annotate the woolly mammoth genome.

We trained a Batch-TopK SAE (Bussmann et al., 2024) on Evo 2’s representations at layer 26, as preliminary analysis indicated that most features of interest were represented at this point (**Figure S5**). The SAE was trained on representations from 1 billion tokens evenly split across complete eukaryotic and prokaryotic genomes (**Figures S5A,B**). We found alignments between learned SAE latent dimensions, also known as features, and known biological concepts by finding features that were enriched in sequence segments containing a particular biological concept of interest, a process we refer to as contrastive feature search (**Figure S6A**).

A close inspection of the learned latent dimensions revealed diverse features that not only align with known biological concepts and genomic building blocks but also capture evolutionary signals embedded within genomes. For example, we made the intriguing observation that Evo 2 has developed internal representations capturing evolutionary signatures of mobile genetic elements. Feature f/19746 is closely associated with prophage regions across prokaryotes (**Figure S6B**) and activates on annotated prophages in the *E. coli* genome including the cryptic prophage CPZ-55 (**Figure 4B**). We further observed that this feature activates on spacer sequences within a CRISPR array, which are integrated during CRISPR adaptation from foreign genetic material such as phage DNA (**Figure 4B**), as well as after the last CRISPR direct repeat and on synthetic, scrambled spacer sequences, suggesting that Evo 2 associates CRISPR spacers with phage sequences as opposed to directly memorizing phage sequences (**Figures 4B** and **S6C**). This feature also activated on other regions not annotated as phage by geNomad (Camargo et al., 2024) which contained genes associated with prophages such as integrases and invertases (**Figure S6D**). Together, these results highlight the potential for mechanistic interpretability of biological language models to enhance genome annotation and to mine for unexplored biological systems.

Next, we sought to identify features that are associated with canonical biological annotation types that reflect genome organization. Through contrastive feature search, we identified diverse features corresponding to open reading frames (ORFs), intergenic regions, tRNAs, and rRNAs in the *E. coli* genome (**Figures 4C** and **S6E,F**). Since these genomic sequences also encode proteins, we further probed for structural signatures at the protein level. Encouragingly, we also identified features linked to protein secondary structures such as α-helices and β-sheets (**Figures 4D** and **S6G**). These associations highlight the multimodal nature of genome language modeling, capturing higher-order structural information beyond the level of DNA alone.

We extended our analysis from *E. coli* to the human genome in search of features that capture the unique regulatory and coding complexities of eukaryotes. By introducing mutations into thousands of human coding sequences and applying contrastive feature search on a eukaryotic-only SAE, we identified a mutation-sensitive feature (f/24278) that preferentially activates on frameshifts and premature stop mutations (**Figures 4E** and **S7A,B**). This suggests that Evo 2 contains latent features beyond simple gene structure that can inform on mutational severity. In addition, with the layer 26 mixed prokaryotic-eukaryotic SAE, we observed other activations on DNA motifs in the promoter region of human genes (**Figure 4F**, left) that closely resemble the known binding sites of human transcription factors (**Figure 4F**, right) (Vorontsov et al., 2023). Collectively, these results suggest that Evo 2 not only recognizes coding sequences but also discerns regulatory elements through distinct internal representations.

Finally, we also identified features closely associated with the exon and intron architecture of the human genome, including features that activate preferentially on coding regions (f/15680), introns (f/28339), the first bases of an exon following an intron (f/1050), and the last base of an exon followed by an intron (f/25666) (**Figure S7C-E)**. The coding region feature also activates on bacterial ORFs, suggesting a learned universal representation of coding sequences (**Figure S6E,F**). To assess whether these features more broadly generalize beyond the human genome, we next applied them to annotate the genome sequence of the woolly mammoth (**Figure 4G**) (Sandoval-Velasco et al., 2024), which was not present in Evo 2’s training corpus. The successful mapping of exon-intron architecture in the woolly mammoth genome underscores the robustness and generalizability of these latent features.

Overall, we demonstrate that Evo 2 latent representations capture a broad spectrum of biologically relevant signals, from mobile genetic elements and regulatory motifs to protein secondary structure and mutational severity. Since conceptual features for natural language can capture abstract concepts, other Evo 2 SAE features likely represent more complex biological patterns (**Figure S5E**). To enable the community to explore these higher-order concepts that bridge mechanistic interpretability with biological mechanisms, we release an accompanying visualization tool for Evo 2 mechanistic interpretability across 104 genomes at https://arcinstitute.org/tools/evo/evo-mech-interp.

### 2.5 Genome-scale generation across the domains of life

Evo 2 is fundamentally a generative model trained to predict the next base pair in a sequence. We sought to generate DNA sequences from different organsism with Evo 2 and assess the quality (**Figure 5A**). In previous work, we demonstrated that the first-generation Evo models can respond to prompt sequences to design and diversify novel biological sequences (Merchant et al., 2024). To evaluate Evo 2’s ability to respond to genomic prompts, we first assessed performance across six phylogenetically diverse species, spanning archaea, prokaryotes, and four eukaryotic lineages (fungi, protists, plants, and animals). For each species, we selected highly conserved representative genes and provided Evo 2 with a context consisting of 1,000 base pairs of upstream sequence plus the first 500-1000 base pairs of the target gene. We found that Evo 2 achieves high accuracy in gene sequence completion, indicating that the model responds to prompts to enable in-context sequence design. Amino acid recovery improved with scale, and Evo 2 40B and 7B demonstrated improved performance compared with Evo 1 (**Figures 5B**. Importantly, Evo 2 also maintained consistently high sequence recovery throughout the context extension midtraining phase (**S8A**).

**Figure 5.**
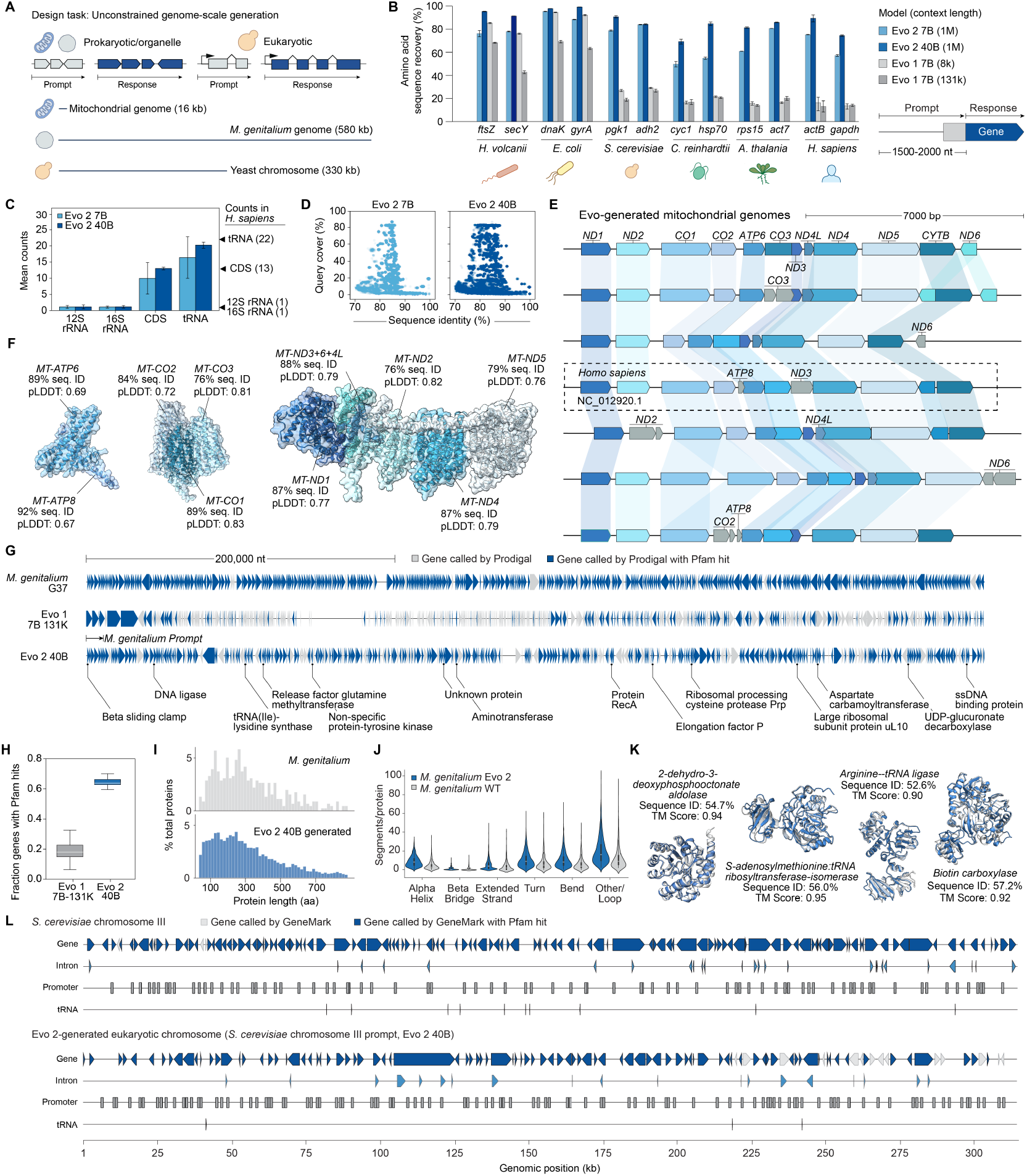
Genome-scale generation across the domains of life. (**A**) We used Evo 2 to generate chromosome- and genome-scale sequences using unconstrained autoregressive generation. We prompted the model with portions of the *H. sapiens* mitochondrial genome, *M. genitalium* genome, and *S. cerevisiae* chromosome III and generate DNA sequences with similar lengths as those of the native sequence. (**B**) We prompted Evo 2 with genomic context as well as a portion of a highly conserved protein and generate, measuring the sequence recovery of the Evo 2 generated gene against the natural gene. (**C**) Predicted rRNA, CDS, and tRNA counts in Evo 2 generated mitochondrial sequences using MitoZ compared with the natural *H. sapiens* mitochondrial genome values. (**D**) Query cover versus sequence identity of generated mitochondrial sequences against nucleotide BLAST hits in the core_nt database with expect threshold of 0.05. (**E**) Visualizations of Evo 2 generated sequences when prompted with a 3 kb sequence from the *H. sapiens* mitochondrial genome, demonstrating variation that still retains natural synteny patterns of coding sequences. (**F**) AlphaFold 3 structure predictions of multimeric complexes from an Evo 2-generated mitochondrial genomes, labeled by the sequence similarity of Evo 2-generated proteins to natural proteins based on a BLASTp query. (**G**) We prompted Evo 2 with the beginning of the *M. genitalium* genome and generated ~ 600 kb long sequences. Genes are annotated with Prodigal and colored based on statistically significant sequence similarity to natural proteins (HHpred E-value < 0.001). (**H**) The fraction of Prodigal annotated genes with HHpred hits between Evo 2 40B and *M. genitalium* generated by Evo 1. (**I**) Distribution of Prodigal annotated genes from Evo 2 generated *M. genitalium* compared with the natural genome. (**J**) Distribution of secondary structure from Evo 2 generated proteins compared to natural *M. genitalium*. (**K**) AlphaFold 3 structure predictions of example proteins found on Evo 2-generated prokaryotic genomic sequences show high structural similarity to natural proteins while diversifying the sequence composition. (**L**) The native genome sequence from *S. cerevisiae* chromosome III and an Evo 2-generated DNA sequence of similar length, which was generated by prompting the model with a 10 kb sequence from *S. cerevisiae* chromosome III, are visualized alongside predicted homologous yeast gene, exon, promoter, and tRNA annotations.

Consistent with poor performance on the language modeling task and on function prediction downstreams for viruses that infect humans, Evo 2 also has poor performance on generating proteins from human viruses (**Figure S2C**). Even when directly trying to elicit a viral protein, Evo 2 had essentially random performance in sequence recovery, effectively preventing Evo 2 from human viral generation.

To test Evo 2’s ability to perform generation at the scale of entire genomes, we first used it to generate mitochondrial DNA and assessed its ability to produce all of the known components of natural mitochondria. The human mitochondrial genome (~ 16 kb) encompasses 22 tRNA genes, 13 protein-coding genes, and 2 rRNA genes, whose protein products form complexes with nuclear-encoded proteins. We prompted Evo 2 40B with portions of human mitochondrial DNA, generating 250 unique mitochondrial sequences of 16 kb each (**Methods**).

We found that Evo 2 can generate mitochondrial genomes with the correct number of coding sequences (CDS), tRNA, and rRNA genes when annotated using MitoZ (Meng et al., 2019) (**Figure 5C**). A BLASTp analysis revealed that Evo 2 created diverse mitochondrial genes, with varying degrees of sequence identity to natural mitochondrial proteins (**Figure 5D**). Notably, individual genes showed highest homology to different organisms ranging from fish to mammals (**Table S6**), highlighting Evo 2’s ability to generate diverse combinations of genes. The generated sequences also maintained proper synteny (**Figure 5E**) while exhibiting considerable sequence diversification when compared to native sequences. Using AlphaFold 3 for protein multimer prediction, we found that the generated sequences formed structures matching expected mitochondrial protein complex folds and interactions (**Figures 5F** and **S8B,C**).

We then leveraged Evo 2’s million-base-pair context window to generate DNA sequences at the same scale as small prokaryotic genomes. For this task, we focused on *M. genitalium*, a model system of a minimal prokaryotic genome due to its short genomic length of ~ 580 kb (Gibson et al., 2008; Karr et al., 2012). Using the first 10.5 kb segment from the *M. genitalium* reference sequence as prompts, we generate ten genomes. We then performed HHpred analysis of Prodigal-predicted ORFs against the Pfam database and found that nearly 70% of Evo 2 40B genes contained significant Pfam hits, a marked improvement over Evo 1 131k (18%) (**Figure 5G,H**). Further, quality assessment of the generated proteins using ESMFold metrics, secondary structure distribution, and protein sequence identity demonstrated that the generated sequences had properties consistent with natural protein distributions, suggesting successful recapitulation of native features despite the long generation length (**Figure 5I-K**).

To assess Evo 2’s eukaryotic sequence generation capability, we prompted Evo 2 with 10.5 kb from *S. cerevisiae* chromosome III (~316 kb in length) to generate 330 kb of DNA. Evo 2 successfully generated eukaryotic-like DNA sequences with predicted tRNAs, appropriately positioned promoters, and genes exhibiting intronic structure (**Methods**; **Figures 5L** and **S8F**). Furthermore, generated proteins showed sequence and structural (**Figure S8G-I**) similarity to natural yeast genes (**Figure S8I**), highlighting Evo 2’s ability to generate realistic eukaryotic coding sequences.

While the density of tRNA and gene features was below those found in the native yeast genome (**Figures 5L** and **S8G**), we note that these genome sequences were produced by simple, unconstrained autoregressive generation. Improvements in the naturalness of generated genomes can most likely be addressed through optimized inference strategies or model improvements. Moreover, as demonstrated in our previous study (Merchant et al., 2024), genomic sequences with no significant sequence similarity to natural sequences could still be semantically valid and retain function.

Together, our results demonstrate that Evo 2 is able to generate DNA sequences that resemble organelle, prokaryotic, and eukaryotic genomes. These generated sequences contain both coding and noncoding elements, with diverse yet realistic genes that maintain both structural and sequence similarity to natural sequences.

### 2.6. Generative epigenomics via inference-time search

Beyond unconstrained autoregressive generation, we also sought to achieve guided generation of long genomic sequences with Evo 2. Eukaryotic genomes regulate gene expression through a complex system of chemical and protein-mediated modifications referred to as the epigenome. An important component of epigenomic regulation involves modifying the openness or compactness of chromatin, which in turn controls which DNA regions can be accessed by transcriptional machinery. Chromatin accessibility is modulated by both DNA sequence composition and aspects of cellular state (Brownell et al., 1996; Allis and Jenuwein, 2016).

Generating DNA sequences to have desirable chromatin accessibility profiles would allow generative genomics to design an important mechanism of functional control found in natural eukaryotic genomes and enable “generative epigenomics.” We therefore aimed to develop an approach that uses Evo 2 to generate DNA sequences for which we can specify both the location and the length of chromatin-accessible regions (**Figure 6A**). For the design tasks in this study, we focus on a binary specification of chromatin accessibility in which we aim to maximize accessibility in some genomic regions and minimize accessibility in other regions.

**Figure 6.**
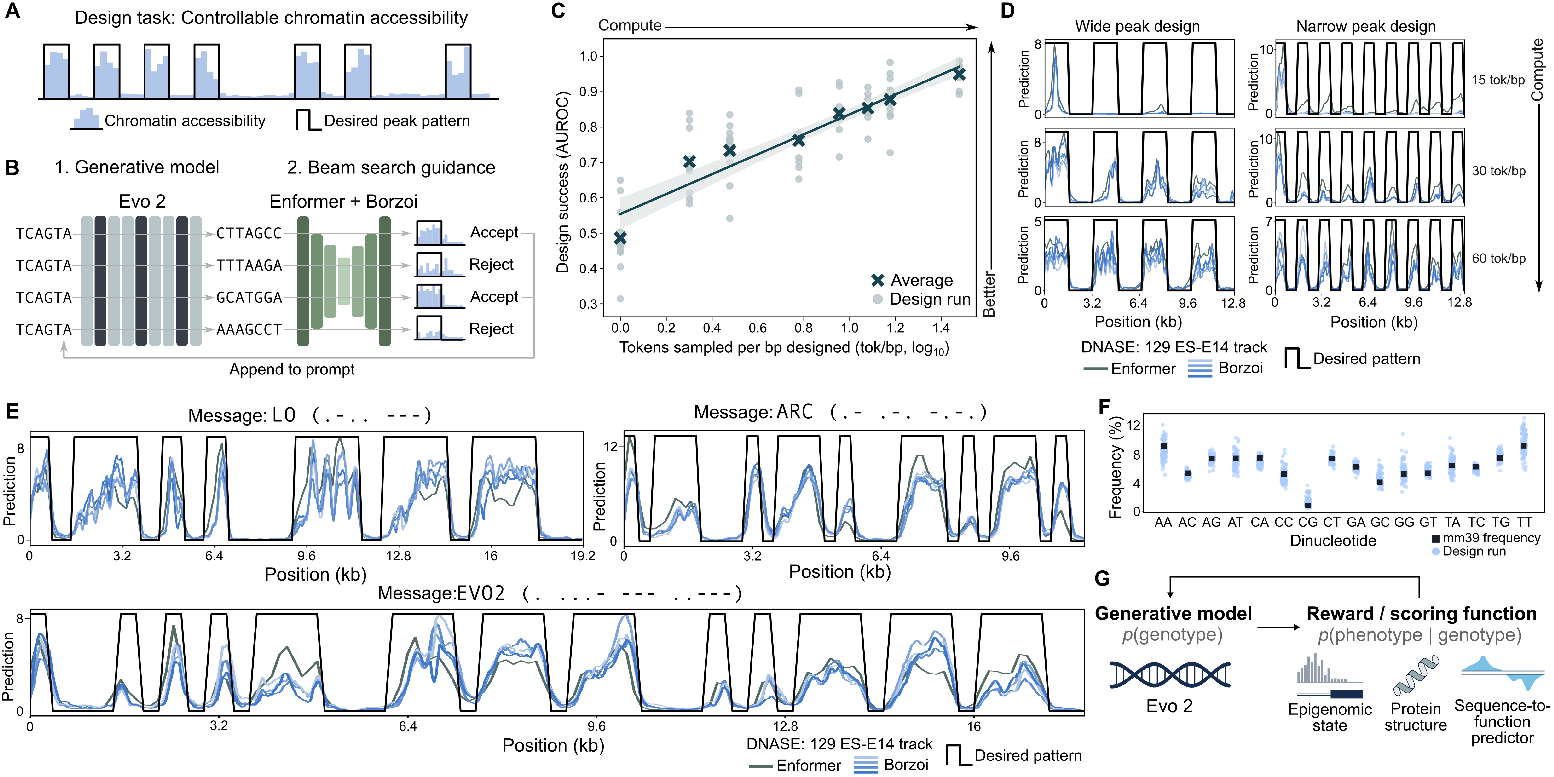
Generative epigenomics via inference-time search. (**A**) We designed multi-kilobase sequences of DNA where we sought to control both the location and length of chromatin-accessible regions. These are often visualized as “peaks” indicating the degree of chromatin accessibility at a given position in the genome. (**B**) We use autoregressive generation with Evo 2 to sample multiple 128-bp chunks of DNA given the same genomic prompt. We then use Enformer and Borzoi to predict how these chunks affect the chromatin accessibility profile of the generated sequence and score these profiles based on how well they match a desired peak pattern. Chunks that have the most desirable accessibility profiles are retained by appending to the prompt and generation proceeds to the next step, using a beam search algorithm to keep track of the best generated sequences. (**C**) Design runs corresponding to different peak patterns are plotted on the horizontal axis according to the total number of tokens sampled during the design process normalized by the length of the design in base pairs (e.g., standard autoregressive decoding would be 1 tok/bp). On the vertical axis, design runs are plotted by using the AUROC, which quantifies how well predicted accessibility profiles can distinguish our desired open- versus closed-chromatin genomic positions. As more tokens are sampled by the model, i.e., as more compute is used, designs more closely adhere to the specified peak pattern. (**D**) Two different peak patterns were designed with varying total compute budgets, with more compute leading to clearer designed peaks. (**E**) Controlling the position and width of chromatin accessibility enables us to encode Morse code messages in the epigenome. Generated DNA sequences replace the native sequence at chrX: 52,051,929–52,123,468 in the mouse genome and are scored by the predicted DNase hypersensitivity tracks in 129 ES-E14 cells according to Enformer and Borzoi. (**F**) Sampled sequences have similar dinucleotide frequencies as the baseline frequencies found in the mm39 reference genome. (**G**) The general paradigm of guiding a capable generative model with a sequence-to-structure or sequence-to-function model extends beyond chromatin accessibility design, enabling many complex biological design applications.

Although Evo 2 has no explicit mechanism for conditioning its generations based on epigenetic state, several models such as Enformer (Avsec et al., 2021) and Borzoi (Linder et al., 2025) demonstrate accurate, held-out performance in predicting chromatin accessibility profiles from DNA sequence across cell types from human and mouse. However, models like Enformer and Borzoi are not trained as generative models and assume that their sequence inputs come from natural genomes. Evo 2 captures complex rules across mammalian genomes (**Figures 2** and **3**) and is a powerful generative model that can be used to propose diverse sequences while retaining biological “naturalness” (**Figure 5**).

To achieve controllable design of chromatin accessibility, we therefore used an ensemble of Enformer and Borzoi to guide autoregressive generation with Evo 2 (**Figure 6B**). Rather than selecting designed sequences solely based on language-model likelihood, we implemented a scoring function that accepts or rejects generated sequences based on how well their predicted chromatin accessibility (using a consensus of Enformer and Borzoi) matches a desired pattern. Instead of sampling a full, multi-kilobase design and then evaluating its predicted chromatin accessibility, we improve the efficiency of the design process by conducting a beam search that scores partial generations; in particular, we reevaluate Enformer and Borzoi after each new 128-bp of sampled sequence and only continue autoregressive generation off of the most promising sequences (**Figure 6B**).

We used both Enformer and Borzoi to score our generated DNA sequences in the context of the mouse genome by replacing chrX: 52,051,929–52,123,468 with the designed sequence. We scored chromatin accessibility using the DNase hypersensitivity tracks in 129 ES-E14 cells according to Enformer and Borzoi, where Borzoi itself consists of four separate models that we ensemble alongside Enformer. Additional details of the generation pipeline are provided in **Methods**.

We first observed that increasing the width of this beam search (i.e., by sampling a greater number of 128-bp chunks given the same prompt and generating off of only the top-scoring chunks) resulted in a substantial improvement in design success (i.e., how well the final chromatin accessibility profiles matched the desired pattern). We use the AUROC metric to quantify how well the continuous-valued Enformer and Borzoi predictions separate regions or open or closed chromatin as specified by our design patterns. Across a diverse set of patterns, we observed that sampling 30 or more 128-bp chunks and selecting the top two chunks at each step of the beam search was sufficient to achieve final designs with AUROCs around 0.9. We observed a predictable log-linear relationship in which increasing the beam search width, and thereby increasing inference-time compute, resulted in better-quality designs (**Figure 6C**). When visualizing the Enformer and Borzoi predictions, we observed clear predicted peaks, indicating high chromatin accessibility, only in regions specified by our design patterns (**Figure 6D**).

To demonstrate the generality of this approach, we varied the length and location of designed peaks in order to write simple messages in Morse code, where narrow peaks indicate dots, wide peaks indicate dashes, and inaccessible regions indicate spaces. Our designed messages include “LO” (the first message transmitted over the Internet and the first word in Edmund Spenser’s *The Faerie Queene*), “ARC” (the name of the research institute in which this design run was conducted), and “EVO2” (the name of the model). We observed consistently strong design success in encoding these diverse messages (**Figure 6E**).

Importantly, using a capable generative model to propose sequences results in designs that have favorable properties that we did not directly optimize for. For example, because we prompted the model with the genomic context from the mouse genome, the dinucleotide frequencies observed in our designed sequences match those of the mm39 reference genome (**Figure 6F**). We also observed that all predictions in the ensemble of Enformer and Borzoi models largely reach a consensus prediction (**Figures 6D,E**). In contrast, when we repeated the same design pipeline except with a uniform proposal replacing Evo 2, we observed suboptimal token-matched inference-time scaling, unnatural dinucleotide distributions, and poor consensus among predictions across the Enformer and Borzoi ensemble, potentially indicating adversarial sequence examples (**Figure S9**).

This design task shows how Evo 2 can be coupled with sequence-to-function models to achieve controllable design of complex properties. In doing so, we demonstrate a clear relationship in which increasing inference-time compute predictably improves performance on a complex design task (**Figure 6C**), which is to our knowledge the first example of such a result for biological language modeling. We also show how allowing the model to propose alternative designs guided by a scoring function can efficiently sample from a functionally complex sequence space. Finally, we note that this paradigm is not exclusive to designing chromatin accessibility. We envision that other models that predict biological structure or function given genomic sequence could also be used to guide Evo 2’s generations (**Figure 6G**), enabling biological design in any downstream application for which there exists a capable predictive model.

## 3. Discussion

This work demonstrates that a generative model of the underlying genomic language enables a machine learning model to achieve generalist prediction and design capabilities across all domains of life. By learning statistical properties of DNA across 9 trillion tokens of genomic sequences, Evo 2 can predict mutational effects on protein function, ncRNA function, and organismal fitness. Evo 2 is the first alignment-free language model that robustly predicts the pathogenicity of different mutation types in ClinVar, including indels, achieving state-of-the-art performance for noncoding and splice variants. Moreover, by leveraging Evo 2 embeddings in supervised classifiers, we achieve state-of-the-art performance in classifying *BRCA1* breast cancer variants across all mutational types. Evo 2 is capable of genome-length sequence design at the scale of whole human mitochondrial genomes, minimal bacterial genomes, or yeast chromosomes. We also show that generation with Evo 2 can produce complex epigenomic patterns via inference-time search. In doing so, we also show that increasing inference-time compute can predictably improve performance on a complex design task. We note that all of these tasks are enabled by a single model.

To understand the basis for Evo 2’s strong zero-shot performance, we applied mechanistic interpretability methods for identifying representational features by decomposing Evo 2’s learned representations with sparse autoencoders (SAEs). Without any explicit biological labels, we report the first examples of SAE features that correlate with diverse elements such as exons, introns, and transcription factor motifs. We also uncovered molecular-level features corresponding to protein secondary structure and organismal-level features that correspond to prophage regions in bacterial genomes. Finally, the evolutionary links captured by these features allow for the genomic annotation of extinct species, which we demonstrated on a portion of the woolly mammoth genome.

Developing Evo 2 required significant investment in machine learning research and engineering. We overcame many important challenges when training and evaluating Evo 2 at scale, including sharding and synchronizing model parameters across tensor, pipeline, context, and data parallel ranks; coordinating events across distributed processes; developing new algorithms and a new architecture to achieve high working utilization of theoretically available GPU compute at datacenter scale; verifying numerical precision across model versions; and engineering a multi-GPU inference stack. We also tested diverse compositions of the pretraining data and found that naive long-context training across raw eukaryotic reference genomes, which mostly consist of noncoding regions, led to poor performance on downstream tasks. Instead, we trained Evo 2 on DNA tokens with high information content, including increasing the composition of training data within or near genic regions. By combining computational engineering and high-quality data curation at massive scale, we were able to achieve the generalist prediction and design capabilities of Evo 2.

Alongside this paper, we freely release a number of resources for the scientific community. Under an opensource license, we make available (i) the model parameters for Evo 2 7B and 40B, (ii) code that implements distributed pretraining and context extension of Evo 2, (iii) code that implements inference of Evo 2, and (iv) the full OpenGenome2 dataset used to train Evo 2 (**Data availability, Code and model availability**). We also release a tool for generating and scoring sequences with Evo 2 in a simple web interface at https://arcinstitute.org/tools/evo/evo-designer and a tool for exploring SAE features alongside genomic annotations at https://arcinstitute.org/tools/evo/evo-mech-interp. To our knowledge, Evo 2 is one of the largest-scale fully open models to date (including training and inference code, data, and parameters) even across other modalities such as language and vision.

Biological foundation models capable of intelligently composing novel systems will advance biomedical innovation, but as with other new biotechnologies, may also raise safety, security and ethical considerations. Aligned with the *Responsible AI x Biodesign* commitments (Responsible AI x Biodesign, 2024), we preemptively assessed and mitigated potential concerns prior to open source publication. Fully open source models enable researchers to interrogate, reproduce, and build upon AI advances. They may also be used by a broad range of actors in unanticipated ways that could lead to accident or misuse risks, as noted by experts following the publication of Evo 1 (Bloomfield et al., 2024). We collaborated with these multidisciplinary experts to reduce risks via data exclusion measures (**Methods**), safety and security evaluations, and population bias evaluations. By excluding genomic sequences of viruses that infect eukaryotes from our training data, we aimed to ensure our openly shared model did not disseminate the capability to manipulate and design pathogenic human viruses. Task-specific post-training may circumvent this risk mitigation measure and should be approached with caution. Our data exclusions had the intended outcomes of weakening language modeling performance (**Figure S2A**) and downstream mutational effect prediction (**Figure S2B**) on human viruses. Red teaming to directly elicit pathogenic human viral proteins showed generations were effectively random in this domain (**Figure S2C**). The inclusion of eukaryotic data also introduced the promising possibility of using Evo 2 to aid in the interpretation of human genetic variants. We queried whether Evo 2’s population-free design (Pathak et al., 2024) mitigated ancestry biases in model predictions, showing that Evo 2 generalizes comparably well across human populations (**Figure S2D**). Few examples of empirical risk assessment of biological foundation models exist; this work represents one of the most comprehensive evaluative efforts that considers both precaution and access to date. However, further research is needed to expand the suite of available evaluations and risk mitigation approaches.

Looking ahead, we anticipate diverse strategies for improving the quality of Evo 2’s predictions and designs. Evo 2 likely emphasizes the general evolutionary distribution of genomic sequences over specific taxonomies. Combining Evo 2 with additional features and population-scale human genomic variation (Schubach et al., 2024; Cheng et al., 2023) could enable improved pathogenicity prediction or analysis of structural variation. Leveraging mechanistic interpretability, learned features could also enhance the detection of more complex biological concepts and guide model generations through activation steering and feature clamping (Templeton et al., 2024), enabling programmable control of generations. Supervised finetuning or reinforcement learning with experimental feedback may be needed to improve the quality of Evo 2’s generated functions. Designing complex biological systems via inference-time compute, for which we provide an initial proof-of-concept in this study, could also generalize to include other properties such as alternative splicing, cell type specificity, or gene circuit functionality.

Evolution represents a unifying theory of biology from genes to populations and transmits the functional effects of natural selection through the foundational information layer of DNA. The Evo series of models lays the groundwork for biological modeling and design that unifies the diverse length scales of biology with a common representation. Future work that integrates this representation with additional modalities such as epigenomic and transcriptomic information could produce a virtual cell model that productively simulates complex cellular phenotypes in health and disease.

## 4. Methods

### 4.1. Training Evo 2

Evo 2 is trained using next-token-prediction on the byte-tokenized OpenGenome2 dataset. We train Evo 2 in two phases: a pretraining phase at 8192 token context focused more on functional elements and midtraining phase during which we extend up to 1M token context length with more entire genomes in the data mix. Evo 2 40B’s pretraining stage is further split into two stages, first training at 1024 context for 6.6T tokens before extending to 8192 context for 1.1T tokens. Additionally, we train and release a smaller, Evo 2 1B base at 8192 context length for 1T tokens. For efficiency, Evo 2 is trained using sequence packing.

**Table 1** provides an estimate of training FLOPS for Evo 2 40B and other large models spanning application domains of biology and natural language. We include Pythia Large (Biderman et al., 2023), OLMo 2 Large (Team OLMo et al., 2024), Evo 1, Falcon 1 180B Almazrouei et al. (2023), StarCoder 2 (Lozhkov et al., 2024), DeepSeek V3 (Liu et al., 2024), Llama 3.1 Large (Dubey et al., 2024), ESM 3 Large (Hayes et al., 2025) and xTrimo (Chen et al., 2024).

#### 4.1.1. Model architecture

Evo 2 uses StripedHyena 2 (Ku et al., 2025), the first multi-hybrid architecture based on input-dependent convolutions (Poli et al., 2023; Nguyen et al., 2024b). Multi-hybrids combine various different operators to balance model quality with training and inference efficiency, in line with findings of Poli et al. (2024) and Thomas et al. (2024). **Figure S1** provides a schematic of each new convolutional operator in the architecture. Self-attention layers use rotary positional embeddings (Su et al., 2024).

Model size information and hyperparameters used for Evo 2 models are shown in **Table 2**. Each Evo 2 model uses a pattern of Hyena-SE, Hyena-MR and Hyena-LI, and attention, with the number of repetitions scaling with model size. Hyena-SE uses inner filters of length 7, Hyena-MR length 128. GELU activations are only used for the first layer, followed by no activations.

**Table 2.** Model architecture configurations for Evo 2 models.

|  | Evo 2 40B | Evo 2 7B | Evo 2 1B base |
| --- | --- | --- | --- |
| Parameters | 40.3B | 6.5B | 1.1B |
| Total Layers | 50 | 32 | 25 |
| Hidden Size | 8,192 | 4,096 | 1,920 |
| FFN Size | 22,528 | 11,264 | 5,120 |
| Num Heads | 64 | 32 | 15 |
| Total Tokens | 9.3T | 2.4T | 1T |

#### 4.1.2. Loss function

Evo 2 is trained with a reweighted cross entropy loss, which weighs the loss contribution of repetitive portions of DNA by 0.1. This affects the genomic window and whole genome portions of the data which contain these annotations. This loss has been found in other DNA models to improve performance on downstream tasks and better calibrate likelihoods between repetitive and nonrepetitive DNA (Benegas et al., 2025), which we found to be true for downstream tasks in a controlled comparison (**Appendix B.1**). The loss is

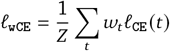

with weighting

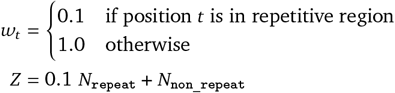

where w_*t*_ is the weight applied to each position, *N*repeats represents the number of positions in repetitive regions within a batch and *N*_non_repeat_ is the number of non repetitive regions, and *Z* is the normalization factor that ensures consistent loss scaling regardless of the proportion of repetitive regions.

For any base pair, the model is always tasked with predicting the uppercase character. For the first 3T tokens of pretraining, lowercase tokens are input to the model to add information on which portions of DNA are repetitive. This was done to further help learn different representations for interspersed repeats, which are very common in many eukaryotic genomes. For additional pretraining and for all midtraining, all inputs to the model are uppercase. Loss is masked on special tokens used to condition the model that we do not want to generate, including the stitch tokens ‘@’ and ‘#’, as well as the multi-token phylogenetic tags used during midtraining.

#### 4.1.3. Pretraining infrastructure

Evo 2 was trained on Savanna (see **Section 6**), custom training infrastructure built with components from DeepSpeed, GPT-NeoX (Andonian et al., 2023), and Transformer Engine. Our stack supports efficient pretraining of multi-hybrid models and new context parallel algorithms. We train our largest models with mixed precision, using a 3D mesh of data, tensor, and context parallelism, combined with ZeRO-3 (Rajbhandari et al., 2020). During training, we use Transformer Engine’s FP8 implementation for linear layers and RMSNorms.

#### 4.1.4. Pretraining phase

**Table 3** provides information on our pretraining configuration. We refer to the models after pretraining but before context extension as Evo 2 40B and 7B base.

**Table 3.** Pretraining hyperparameters for Evo 2 models. Each model uses the AdamW optimizer with β_1_ = 0.9, β_2_ = 0.95, and cosine learning rate decay.

|  | Evo 2 40B base | Evo 2 7B base | Evo 2 1B base |
| --- | --- | --- | --- |
| Learning Rate | 2.0e-4 | 3.0e-4 | 3.0e-4 |
| Training Batch | 16.8M | 4.2M | 2.1M |
| Total Iterations | 516K | 500K | 490K |
| Pretraining Tokens | 8.7T | 2.1T | 1T |
| Sequence length | 1024 (6.6T), 8192 (1.1T) | 8192 | 8192 |

#### 4.1.5. Midtraining phase: Context extension

We follow a multi-stage midtraining procedure, gradually extending the context length while keeping the same batch size as pretraining, adjusting model parallelism accordingly. Midtraining was performed on an adjusted data composition, including more whole genomes and with longer average sequence length (**Figure 1, Appendix S5**).

We explore two different rotary embedding-based methods to adapt to longer sequences: positional interpolation by down scaling the positional index of tokens (Chen et al., 2023) and increasing the base frequency of the RoPE embedding (Xiong et al., 2023). We decide on a combined approach using both together, with a 10× increase to the base frequency for every doubling of the context length. **Table 4** provides information on our midtraining protocol.

**Table 4.** Context extension protocol and number of tokens for Evo 2 7B and 40B.

| Context Length | Base Frequency | Scale Factor | Evo 2 7B<br>Training Tokens | Evo 2 40B<br>Training Tokens |
| --- | --- | --- | --- | --- |
| 32.8K | $1.0 \times 10^6$ | 4 | 50B | - |
| 65.5K | $1.0 \times 10^7$ | 8 | 50B | - |
| 131.1K | $1.0 \times 10^8$ | 16 | 50B | 200B |
| 262.1K | $1.0 \times 10^9$ | 32 | 50B | 200B |
| 524.3K | $1.0 \times 10^{10}$ | 64 | 50B | - |
| 1048.6K | $1.0 \times 10^{11}$ | 128 | 50B | 200B |
|  |  |  | 300B | 600B |

After extension stages, model performance was evaluated using loss, performance on short sequence DMS tasks, and performance on our long context needle-in-a-haystack evaluation to evaluate effectiveness of the extension. We did not find significant differences between different extension protocols. Based on the successful context extension results for the 7B, we reduced the number of stages and increase the number of tokens for the 40B, seeing 600B tokens during midtraining.

#### 4.1.6. Midtraining phase: Needle-in-a-haystack evaluation

We developed a novel synthetic evaluation to assess the ability of DNA language models to identify and utilize a specific sequence pattern in its context to make predictions on a repeated sequence with the same pattern. This “needle-in-haystack” evaluation quantifies a model’s capacity to retrieve sequence patterns within different context lengths. For each evaluation, we generated a random DNA sequence of 100 base pairs (bp) to serve as the “needle” sequence. A background sequence (“haystack”) was constructed by sampling DNA base pairs uniformly at random at varying lengths following powers of two, ranging from 512 bp to 1,048,576 bp. The needle sequence was systematically inserted at different relative positions within each haystack, specifically at depths corresponding to 10% through 90%, at intervals of 10%, of the total haystack length. A “query” sequence, which is an exact duplicate of the needle sequence, was then placed at the suffix of the haystack sequence.

The evaluation methodology employs a modification of the “categorical Jacobian” analysis, as originally proposed by Zhang et al. (2024), to measure the model’s use of the needle sequence to predict the query sequence. At a high-level, we mutate the needle and measure the effects on model predictions for the query as a way to assess retrieval. More formally, let **C** denote the categorical Jacobian matrix using the notation from Nguyen et al. (2024a), where the entry **C** [*i, j*] indicates the Euclidean magnitude of the difference in logits at position *j* when mutating position *i* to all tokens in the vocabulary. We compute a retrieval score

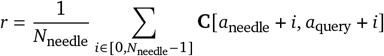

where *N*_needle_ = 100 is the length of the needle (and of the query) sequence, *a*_needle_ is the starting position of the needle and *a*_query_ is the starting position of the query. High values of *r* indicate greater retrieval strength. We use a threshold of *r* ≥ 0.8 to determine successful retrieval, which was obtained by manually inspecting categorical Jacobian matrices of synthetic sequences containing repeated motifs. We computed the retrieval score for various haystack lengths and when inserting the needle at various depths into the haystack.

#### 4.1.7. Inference infrastructure

Evo 2 inference runs on Vortex (see **Section 6**). Vortex contains infrastructure and efficient implementation for autoregressive generation with StripedHyena 2. For the new convolution operators with finite inner filters, we adopt a caching strategy similar to KV caching in self-attention. For long filters, we switch to a recurrent form. All convolution operators in the architecture can generate autoregressively with a constant memory footprint.

### 4.2. OpenGenome2 training data

We significantly expanded upon the OpenGenome pretraining dataset, used to train Evo 1, to create OpenGenome2, increasing the total number of nucleotides from 300 billion to 8.84 trillion. This included a 33% expansion of representative prokaryotic genomes from 85,205 to 113,379 (357 billion nucleotides), a total of 6.98 trillion nucleotides from eukaryotic genomes, 854 billion nucleotides of non-redundant metagenomic sequencing data, 2.82 billion nucleotides of organelle genomes, and 602 billion nucleotides of subsets of eukaryotic sequence data to focus on likely functional regions of the genomes by focusing on different windows around coding genes. We introduced these data augmentations to prioritize genes and regions around genes to improve performance on downstream tasks (**Appendix B.2**).

#### 4.2.1. Data curation

##### Reused datasets

Our previously published OpenGenome dataset was used in its entirety as part of the training data for this study (Nguyen et al., 2024a). This included representative prokaryotic genomes available through GTDB release v214.1, and curated phage and plasmid sequences retrieved through IMG/VR and IMG/PR. As previously described, the OpenGenome dataset was curated to exclude genomic sequences of viruses which infect eukaryotic hosts.

##### Updated prokaryotic genomes

New prokaryotic reference genomes made available through the GTDB release 220.0 (Parks et al., 2022) update were added to the training data for this study. New genomes were identified by selecting all species’ reference genomes that had no previously published (release 214.1) genomes within their species cluster, resulting in 28,174 additional prokaryotic genomes.

##### Eukaryotic reference genomes

All available eukaryotic reference genomes were downloaded from NCBI on May 31, 2024, excluding atypical genomes, metagenome-assembled genomes, and genomes from large multi-isolate projects. This resulted in 16,704 genomes including an estimated 10.7 trillion nucleotides. Only contigs that were annotated as ‘Primary Assembly’, ‘non-nuclear’, or ‘aGasCar1.hap1’ (an aberrant annotation that applied only to GCA_027917425.1) were retained. Mash sketch was run on each individual genome with the flag “-s 10000” and the mash distance was calculated between all genomes as an estimate for their pairwise 1-ANI (average nucleotide identity) (Ondov et al., 2016). All genomes with a mash distance < 0.01 were joined with edges in a graph (Csárdi et al., 2024), and clusters were identified by finding connected components. One representative genome per cluster was chosen, prioritizing genomes with a higher assembly level and genomes with longer total sequence length. This clustering resulted in 15,148 candidate genomes. Genomes were further filtered by removing ambiguous nucleotides at the termini of each contig, by removing regions annotated as “centromere” in an available GFF file, and by removing contigs that were less than 10 kb in total length. Finally, contigs that were composed of more than 5% ambiguous nucleotides were removed. This final filtered set included 15,032 genomes and 6.98 trillion nucleotides.

##### Metagenomes

A previously described metagenomics dataset (Durrant et al., 2024) was further curated as part of the training data. This included 41,253 metagenomes and metagenome-assembled genomes from NCBI, JGI IMG (Chen et al., 2021), MGnify (Mitchell et al., 2020), MG-RAST (Meyer et al., 2008), Tara Oceans samples (Sunagawa et al., 2015), and Youngblut et al. animal gut metagenomes (Youngblut et al., 2020). All contigs were split at consecutive stretches of ambiguous nucleotides of length 5 bp or longer, the split contigs were filtered by a minimum sequence length of 1 kb, and only contigs with at least one open reading frame as predicted by Prodigal (Hyatt et al., 2010) were kept. Contig-encoded proteins were previously clustered at 90% identity using MMseqs (Durrant et al., 2024; Steinegger and Söding, 2017). To further remove redundant sequences, contigs were sorted by descending length, and each contig was only retained if at least 90% of its respective protein clusters were not already in the sequence collection (determined using a bloom filter from the pybloom package with a capacity of 1614960255 and an error rate of 1e-6).

##### Eukaryotic organelle genomes

33,457 organelle genomes were identified and downloaded using the “NCBI Organelle” web resource. Ambiguous nucleotides at the terminal ends of the organelle genome sequences were removed. Sequences that had over 25 ambiguous nucleotides were removed. This resulted in 32,240 organelle genomes that were used for training, including 17,613 mitochondria, 12,856 chloroplasts, 1,751 plastids, 18 apicoplasts, 1 cyanelle, and 1 kinetoplast.

##### mRNA and ncRNA transcripts

Transcripts were extracted using GTF files that were available through NCBI for 4,390 reference genomes. All transcripts were extracted using these coordinates, and the longest representative transcript per gene was selected to limit sequence redundancy. Transcripts from each representative genome were then filtered to be at least 64 nucleotides in length and less than 100 kb in length, and transcripts that had consecutive stretches of ambiguous nucleotides that were 5 bp or longer were removed. Transcripts were clustered with mmseqs at 90% identity for each genome to reduce redundancy. Transcripts were then split into “mRNA” and ncRNA (not-mRNA) by using the gbkey field of the GTF file. All mRNA and ncRNA transcripts across all species were then grouped together, and separately clustered again using mmseqs at 90% identity.

##### Noncoding RNAs

ncRNA sequences were obtained from Ensembl (release 112), Rfam, and RNAcentral. For Ensembl, we used sequences explicitly annotated as “ncrna” across 338 reference genomes. For Rfam and RNAcentral, we downloaded all available sequences in each database. All ncRNA sequences were then combined into a single FASTA file and then clustered with mmseqs easy-linclust with parameters –min-seq-id 0.9, -c 0.8, and –cov-mode 0 to produce the final set of sequences for training.

##### Eukaryotic promoters (EPDnew)

Sequences from 15 organisms consisting of 600 bp representing positions *−*499 to 100 relative to the transcription start sites, which correspond to experimentally validated promoter sequences, were obtained from the EPDnew database. These sequences were then clustered with mmseqs easy-linclust with parameters –min-seq-id 0.9, -c 0.8, and –cov-mode 0 to produce the final set of sequences for training.

#### 4.2.2. Data processing and tokenization

Datasets were preprocessed differently for pretraining and midtraining to reflect the different lengths of training. We augment the data in order to focus model pretraining on more conserved, information dense regions around genes, inspired by previous approaches (Benegas et al., 2023). To enable data augmentations and stitching of multiple contigs together, we introduce two special tokens. The ‘#’ token is used to join sequences from the same species with uncertain distance to each other, while the ‘@’ token is used for sequences that are from the same contig/strand and are near each other. We perform a data ablation to test the effectiveness of our data composition compared to one with fewer augmentations (**Appendix B.2**).

For pretraining, we perform the following additional filtering, processing, and augmentation strategies. We use both ncRNA and mRNA transcript annotations to generate the augmented transcripts and gene windows data portions.

##### GTDB and IMG/PR

We reverse complement 50% of the sequences. A random set of 100 sequences was used for validation.

##### Transcripts

Sequences with 3 continuous ‘N’ nucleotides were removed, and all uracil is replaced with thymine. We reverse complement 50% of the sequences. A random set of 1000 sequences is held out for validation.

##### Augmented transcripts

**Eukaryote promoter, exon, and splice overhangs** As a separate data augmentation, the same set of clustered mRNA and ncRNA transcripts (see **Section 4.2.1**) were modified to include an additional 1,024 bp of starting sequence and an additional 32 bp around each exon for additional splice site information. These “stitched” transcript sequences were then combined together for each transcript using a special “@” token.

##### Eukaryotic genic regions

We use windows around annotated genes to enrich functional coding and cis regulatory elements in the training data. Using the same GTF gene coordinates previously retrieved from NCBI (see **Section 4.2.1**), we created an augmented collection of eukaryotic sequences that were enriched around coding and noncoding exons. All transcripts that were retained after all filtering steps of the mRNA and ncRNA transcript curation process (see **Section 4.2.1**) were identified, with 5000 bp from both sides of each exon coordinate. These coordinates were then merged using bedtools (Quinlan and Hall, 2010) in a strand-agnostic manner, and contiguous stretches of sequence were then extracted from each respective genome sequence file. All extracted sequences were then split at consecutive stretches of ambiguous nucleotides of length 5 bp or longer, and the remaining sequences were filtered to be at least 1,000 nucleotides in length. The ‘@’ token is used to join sequences from the same contig. 50% of entire joined sequences are reverse complemented.

For midtraining, we increased the effective length of sequences to take advantage of the model’s extended context window. We achieved this by stitching together sequences from the same accession with different strategies for each dataset, leading to effective sequence lengths of millions of base pairs for prokaryotic and eukaryotic genomes so that the model sees relevant sequence in its entire window during training.

##### GTDB

Sequences from the same organism are joined together with a special ‘#’ token at the gaps. This increases the median sequence length from 12 kb to 2 million base pairs. Phylogenetic tags are added every 131 kb to help condition the model.

##### IMG/VR

Phylogenetic tags are added at the beginning of every IMG/VR sequence

##### Eukaryotic genomes

Sequences from the same organism are joined together. ‘@’ is used for sequences from the same contig, while the ‘#’ is used to join sequences from different contigs. This increases the median sequence length from 15 kb to millions.

Transcripts, augmented transcripts, genomic windows, and IMGPR remain the same as for the pretraining phase.

Phylogenetic tags are included help condition the model during midtraining, and loss is ignored for these tokens. Phylogenetic tags are formatted Greengenes-style lineage strings which concatenate all taxa starting domain to species, with all uppercasing separated by semicolons. ‘|’ tokens are added at the start and end, similar to in Evo 1 (Nguyen et al., 2024a). For example, the tag for *E. coli* would be:

~~~
|D__BACTERIA;P__PSEUDOMONADOTA;C__GAMMAPROTEOBACTERIA;
O__ENTEROBACTERALES;F__ENTEROBACTERIACEAE;G__ESCHERICHIA;
S__ESCHERICHIA|
~~~

#### 4.2.3. UMAP visualization of whole genomes

We created a UMAP visualization of all prokaryotic and eukaryotic representative genomes used in the training data as an illustrative representation of their abundance and diversity. K-mer frequencies were calculated for all prokaryotic and eukaryotic genomes using jellyfish (Marçais and Kingsford, 2011) and k values 1 through 6. These were combined into a single data frame, and scikit-learn (Pedregosa et al., 2011) was used to scale the data. To better separate the domains, k-mer composition vectors were multiplied by 2 for archaeal species and by 3 for eukaryotic species. The umap.UMAP function was used to then calculate the UMAP with parameters n_neighbors=15, min_dist=0.5, and default parameters otherwise (McInnes et al., 2018).

### 4.3. Prediction evaluations

#### 4.3.1. Effect of mutations on Evo 2 likelihoods around start codons

Reference genome sequences and their annotations for 20 prokaryotic and 16 eukaryotic species were obtained from NCBI. From the annotations of each species, 1,024 protein-coding genes were randomly sampled, with the exception of *N. equitans*, for which all of its 536 protein-coding genes were selected. For each of these sampled genes, we selected genomic coordinates ranging from −20nt to +20nt from the first base of the start codon, and mutated the wildtype base of each position to each of the three alternative bases to introduce SNVs. Then, using the Evo 2 7B model, we calculated the difference in the likelihoods of the SNVs to their respective wildtype sequences, both of which included the genomic context of a 8,192nt window that is centered around the mutated nucleotide. In calculating the log likelihoods of both wildtype sequences and their SNVs, we used the average likelihoods of the original sequence and its reverse complement. These delta likelihoods were averaged across the 1,024 sampled genes per each position for each species. The same process was used for the variant effects around stop codons. The phylogenetic trees for both prokaryotic and eukaryotic species were constructed using existing literature (Hug et al., 2016; Hartmann et al., 2006; Hedges, 2002).

#### 4.3.2 Effect of prokaryotic mutations on Evo 2 likelihoods

We systematically introduced artificial mutations across different genomic regions in prokaryotic genomes. Annotated reference genomes of 20 prokarytoic species shown below were obtained through NCBI.

- *Escherichia coli* (GCF_000005845.2)
- *Bacillus subtilis* (GCF_000009045.1)
- *Synechocystis sp. PCC* (GCF_000019485.1)
- *Mycobacterium tuberculosis* (GCF_002357975.1)
- *Bacteroides thetaiotaomicron* (GCF_001314975.1)
- *Methanococcus maripaludis* (GCF_002945325.1)
- *Nitrosopumilus maritimus* (GCF_000018465.1)
- *Acinetobacter baumannii* (GCF_001628795.1)
- *Enterococcus faecium* (GCF_009734005.1)
- *Klebsiella pneumoniae* (GCF_000968155.1)
- *Neisseria gonorrhoeae* (GCF_023822665.1)
- *Neisseria meningitidis* (GCF_015679665.1)
- *Pseudomonas aeruginosa* (GCF_000626655.2)
- *Staphylococcus aureus* (GCF_003609855.1)
- *Methanocaldococcus jannaschii* (GCF_000091665.1)
- *Sulfolobus solfataricus* (GCF_000968435.2)
- *Haloferax volcanii* (GCF_010692905.1)
- *Thermococcus kodakarensis* (GCF_028471865.1)
- *Nanoarchaeum equitans* (GCF_000008085.1)
- *Streptomyces coelicolor* (GCF_008931305.1)

For each species, 5,000 positions were randomly sampled from bases annotated as coding regions, and 2,000 positions each were sampled from bases within loci annotated as rRNAs, tRNAs, and ncRNAs, respectively. Positions not annotated as CDS, rRNA, tRNA, or ncRNA were considered to be intergenic regions, from which 2,000 positions were sampled for each species. Centered around each sampled position, a 8,192 bp window was used as the genomic context for the calculation of likelihoods with Evo 7B. For each sampled position in the CDS, the wildtype base was mutated to each of the three alternative bases to introduce SNVs, and deleted for the 1 bp deletion mutant. For each sampled position in rRNA, tRNA, ncRNA, or intergenic regions, the 10 bp window surrounding the position was deleted for the 10 bp deletion variant. For the deletion variants, the 8,192 bp window was extended into the neighboring sequences to match the length of the wildtype sequence (8,192 bp) to avoid any biases in the likelihoods that arise from differences in sequence lengths. Whether an SNV in the coding region was synonymous, missense, or a nonsense mutation was determined using the standard codon table for all species.

#### 4.3.3. Effect of eukaryotic mutations on Evo 2 likelihoods

We systematically introduced artificial mutations across different genomic regions in eukaryotic genomes. Gene annotations were extracted from GFF3 and GTF files obtained through Ensembl (Harrison et al., 2024) or NCBI. Reference genome sequences for 16 eukaryotic species were used:

- *Arabidopsis thaliana* (GCF_000001735.4)
- *Caenorhabditis elegans* (GCA_000002985.3)
- *Callithrix jacchus* (GCA_011100555.1)
- *Chlamydomonas reinhardtii* (GCA_000002595.3)
- *Drosophila melanogaster* (GCF_000001215.4)
- *Danio rerio* (GCA_000002035.4)
- *Homo sapiens* (GCF_000001405.40)
- *Macaca mulatta* (GCF_003339765.1)
- *Mus musculus* (GCF_000001635.27)
- *Nicotiana attenuata* (GCA_001879085.1)
- *Oryza sativa* (GCA_001433935.1)
- *Paramecium tetraurelia* (GCA_000165425.1)
- *Saccharomyces cerevisiae* (GCA_000146045.2)
- *Thalassiosira pseudonana* (GCA_000149405.2)
- *Tetrahymena thermophila* (GCA_000189635.1)
- *Xenopus tropicalis* (GCF_000004195.4)

Mutations were generated in coding sequences, including synonymous substitutions, nonsynonymous substitutions, premature stop codons, and single nucleotide deletions (frameshift). Noncoding regions include 5′ UTRs, 3′ UTRs, introns, intergenic sequences, and noncoding RNA exons annotated as lncRNA (lncRNA, lincRNA, or lnc_RNA), snoRNA, miRNA (miRNA, pre_miRNA), snRNA, tRNA (tRNA or source=trnascan), rRNA, or ncRNA (ncRNA or ncRNA_gene). Noncoding regions were mutated with 10 bp deletions. 8,192 bp of sequence context around each mutation position were used, and additional flanking sequence was appended to the ends of the sequence in the case of deletions to keep the length of the sequences consistent. Species-specific codon tables were used to identify appropriate premature stop codons and substitutions. Up to 200,000 sequences were sampled across sequence types to maintain balanced representation. For each unique region coordinate (i.e. a specific exon or intron) up to 20 distinct mutations were retained. For each region and mutation type, the median change in likelihood was calculated as the representative change in likelihood.

#### 4.3.4. Stop codon analysis across genetic codes

To test the ability of the Evo 2 model to understand variations in the genetic code usage across different species, we introduced premature stop codons into coding sequences across 5 species: *Arabidopsis thaliana* (GCF_000001735.4, standard code), *Homo sapiens* (GCF_000001405.40, standard code), *Mycoplasma pneumoniae* (GCF_900660465.1, mycoplasma code), *Thalassiosira pseudonana* (GCA_000149405.2, ciliate code), and *Tetrahymena thermophila* (GCA_000189635.1, ciliate code). GFF3 and GTF annotation files were used to identify coding sequences. Coding sequences and mutation positions were randomly selected, filtering out pseudogenes and sequences shorter than a minimum length threshold of 10. For each selected coding sequence, randomly selected codons were replaced with seven different stop codons (TAA, TAG, TGA, AGG, TCA, AGA, TTA). Sequences were randomly subsampled to at most 200,000 per genome. The likelihood of the reference sequence and mutated sequences was calculated with the Evo2 7B model with sequence context windows ranging from 100 to 8,192 base pairs centered on the mutation site. The change in likelihood was calculated for each mutation, and the median of these values for each codon was calculated for each species. These median values were *z*-score standardized across all 7 tested codons.

#### 4.3.5. Noncoding regulatory sequence evaluations (DART-Eval)

For zero-shot evaluations in DART-Eval (Patel et al., 2024), we assessed model performance on Task 1 (CCRE), Task 2 (TF Motif), and Task 5 (Variant Effect Prediction). Tasks 1 and 2 focused on likelihood-based evaluations: Task 1 involved distinguishing candidate cis-regulatory elements (cCREs) from shuffled control sequences, using a dataset of over 2.3 million sequences derived from experimentally validated regulatory regions, while Task 2 aimed to identify transcription factor (TF) motifs by differentiating true TF binding sites from control sequences, utilizing a dataset of approximately 577,000 sequences from TF footprinting experiments.

Task 5 evaluated variant effects through both likelihood-based and embedding-based analyses. This task included two datasets: one examining chromatin accessibility QTLs (CaQTLs) across African populations, with over 219,000 variant sites, and another focusing on Yoruba dynamic sequence QTLs (dsQTLs), with approximately 28,000 sites. For the embedding-based analysis, Evo 2 embeddings were first extracted from Block 13, which was randomly selected for both the 7B and 40B models. Additionally, Block 27 was then chosen for the Evo 2 7B model while Block 20 was selected for the Evo 2 40B model due to their superior performance on the *BRCA1* variant supervised classification task (see **Section 4.3.16**). We selected these tasks to emphasize Evo 2’s zero-shot capabilities for regulatory sequence prediction across various data types. Results across all three tasks were compared to pre-computed results from testing various models in the DART-Eval paper, including GENA-LM (bert-large-t2t), HyenaDNA (large-1m), DNABERT-2 (117M), and NT 500M (v2-500m-multi-species).

#### 4.3.6. Zero-shot protein fitness prediction

We conducted zero-shot fitness prediction for protein and ncRNA sequences as described in Nguyen et al. (2024a). We previously compiled nine datasets of prokaryotic DMS datasets from ProteinGym in which the original study authors had provided readily accessible nucleotide and protein sequences: a β-lactamase DMS by Firnberg et al., a β-lactamase DMS by Jacquier et al., a CcdB DMS, a multiprotein thermostability dataset, an IF-1 DMS, an Rnc DMS, an HaeIII DMS, a VIM-2 DMS, and an APH(3′)II DMS. See ProteinGym (Notin et al., 2023) for a list of references to these studies. We also compiled six datasets of human proteins from Livesey and Marsh (2023) in which the original study authors had provided readily accessible nucleotide and protein sequences: a CBS DMS, a GDI1 DMS, a PDE3A DMS, a P53 DMS by Kotler et al., a P53 DMS by Giacomelli et al., and a BRCA1 DMS. See Livesey and Marsh (2023) for references to these studies.

For this study, we also compiled a set of 18 DMS datasets corresponding to proteins from viruses that infect humans: an HCV polymerase DMS by Qi et al., an influenza hemagglutinin DMS by Wu et al., an influenza nucleoprotein DMS and a PB1 DMS by Doud et al., an influenza PA DMS by Wu et al., an influenza neuraminidase DMS by Jiang et al., an HIV-1 TAT DMS and an HIV-1 REV DMS by Fernandes et al., an HIV-1 envelope DMS by Haddox et al., an HIV-1 envelope DMS by Duenas-Decamp et al., an influenza hemagglutinin DMS by Lee et al., an HIV-1 envelope DMS by Haddox et al., an influenza PB2 DMS by Soh et al., a Zika envelope DMS by Sourisseau et al., a SARS-CoV-2 spike RBD DMS by Starr et al., a coxsackievirus capsid DMS by Mattenberger et al., an AAV2 capsid DMS by Sinai et al., a SARS-CoV-2 Mpro DMS by Flynn et al., a dengue virus NS5 DMS by Suphatrakul et al., and an influenza PB1 DMS by Li et al. See ProteinGym (Notin et al., 2023) for a list of references to these studies.

For comparing nucleotide and protein language models on bacterial and human DMS datasets, we utilized the complete set of unique nucleotide sequences and their corresponding fitness values exactly as reported in the original studies. When discrepancies arose between fitness values reported for nucleotide sequences versus their protein counterparts, we used the fitness values for the nucleotide sequences; in these cases, we evaluated the protein language models using the translated sequence. For mutations involving stop codons, which were reported in some studies, we included these sequences when evaluating the nucleotide language models but excluded them from the protein language model benchmark. For human viral proteins, we used the wildtype protein sequence and substitutions as reported by ProteinGym to evaluate protein language models.

To evaluate the nucleotide language models on human viral proteins, we manually retrieved the nucleotide sequence of each protein from the GenBank entry associated with each protein’s UniProt entry. For each amino acid substitution in ProteinGym, we used the most frequently used nucleotide codon in the human codon table corresponding to the mutant amino acid value.

To assess model performance, we calculated the Spearman correlation between the experimental fitness values and the model-derived sequence scores. For autoregressive language models, we used sequence likelihood as the score, while for masked language models, we used sequence pseudolikelihood. We compared the Evo 2 40B, Evo 2 7B, Evo 1, GenSLM (Zvyagin et al., 2023), Nucleotide Transformer (Dalla-Torre et al., 2024), and RNA-FM (Chen et al., 2022) nucleotide language models and the CARP-640M (Yang et al., 2024), ESM-1v (Meier et al., 2021), ESM-2 650M, ESM-2 3B (Lin et al., 2023), ProGen2-large, and ProGen2-xlarge (Nijkamp et al., 2023) protein language models.

#### 4.3.7. Zero-shot ncRNA fitness prediction

We previously compiled nine datasets of DMS datasets on ncRNA (Nguyen et al., 2024a): a ribozyme DMS by Kobori et al., a ribozyme DMS by Andreasson et al., a tRNA DMS by Domingo et al., a tRNA DMS by Guy et al., a ribozyme DMS by Hayden et al., a ribozyme DMS by Pitt et al., and a rRNA mutagenesis study by Zhang et al. See Nguyen et al. (2024a) for a list of references to these studies. To assess model performance, we calculated the Spearman correlation between the experimental fitness values and the model-derived sequence scores. For autoregressive language models, we used sequence likelihood as the score, while for masked language models, we used sequence pseudolikelihood. We compared the Evo 2 40B, Evo 2 7B, Evo 1, GenSLM, Nucleotide Transformer, RNA-FM, RNAErnie (Wang et al., 2024), and RiNALMo (Penić et al., 2024) nucleotide language models.

#### 4.3.8. Zero-shot mRNA decay evaluation

For this evaluation, we used a human cell line dataset that leverages metabolic RNA labeling to estimate mRNA decay rates across the transcriptome. We used the average values across the reported lines as a measure of overall mRNA decay rates. We then used the Evo 2 40B and 7B models, along with Nucleotide Transformer, RNA-FM, and RiNALMo to compare sequence scores to mRNA decay rates. For inputs to each model, we either used the full-length mRNA or the longest context allowed by the model from the end of the transcript. Since model scores may be confounded by sequence length, we first applied loess regression to correct for variations in transcript length.

#### 4.3.9. Exon/intron classification

To evaluate model embeddings’ ability to classify genomic positions as exonic or intronic across diverse eukaryotes, we selected 94 available eukaryotic species from PANTHER19.0 (Mi et al., 2013). We partitioned organisms into training (80%), hyperparameter optimization (10%), and test (10%) sets, with *Homo sapiens, Mus musculus*, and *Danio rerio* manually assigned to the test set after random partitioning. *Trichomonas vaginalis* and *Leishmania major*, originally in the test set, were excluded from evaluation due to insufficient intronic annotations. All computations were performed on a single NVIDIA H100 Tensor Core GPU. Model versions used were evo2_7b_gen for Evo 2, evo-1-8k-base for Evo 1 (Nguyen et al., 2024a), and nucleotide-transformer-2.5b-multi-species for Nucleotide Transformer (Dalla-Torre et al., 2024).

For each species, we randomly sampled positions from NCBI RefSeq-annotated (O’Leary et al., 2016) long noncoding and protein-coding genes in an unbiased manner. The latest version of RefSeq FASTA and GTF files were downloaded on November 18th, 2024. All exon annotations from RefSeq GTF files were included, without distinguishing between constitutive and alternative exons. For each position, we extracted both forward and reverse strand sequences from reference genomes up to each model’s maximum length (8192 bp for Evo 2 and Evo 1; 5994 bp for Nucleotide Transformer), with the target position at the 3′ end of each. We concatenated the forward and reverse strand embeddings as classifier input after keeping all embedding dimensions and only the final sequence position.

We evaluated both linear classifiers and single-hidden layer perceptrons. For initial optimization, we sampled 150 positions per species and extracted model embeddings from each model’s top level layers. Using weighted binary cross-entropy loss, we trained classifiers for all layers and selected the best performing layer based on validation accuracy. We then optimized hyperparameters (number of hidden layers (0 or 1), hidden layer dimension, learning rate, batch size, and weight decay) using Tree-structured Parzen Estimators (TPE) on the selected layer, choosing the configuration that maximized validation accuracy. For Evo 2 - layer: blocks.26, number of hidden layers: 1, hidden layer dimension: 1024, learning rate: 5 × 10^−5^, batch size: 16, and weight decay: 2 ×10^−4^. For Nucleotide Transformer - layer: 24, number of hidden layers: 1, hidden layer dimension: 1024, learning rate: 5 × 10^−4^, batch size: 64, and weight decay: 2× 10^−7^. For Evo 1 - layer: blocks.3, number of hidden layers: 0, learning rate: 5 ×10^−5^, batch size: 64, and weight decay: 1.5 ×10^−6^. Final classifiers were trained on 1500 positions per species using these optimal parameters.

#### 4.3.10. Gene essentiality

We conducted zero-shot gene essentiality prediction as previously described (Nguyen et al., 2024a). We obtained binary essentiality data (labeled as “essential” or “nonessential”) for 56 bacterial genomes from the DEG database (Zhang, 2004). Additionally, we incorporated genome-wide essentiality data for two phage genomes, lambda and P1, from Piya et al. (2023), using the binary labels that the study authors assigned based on their CRISPRi screen results.

To conduct our in silico gene essentiality screen, we accessed the complete bacterial genomes using DEG-provided RefSeq IDs. For the phage genomes, we used RefSeq: NC_001416 (lambda phage) and RefSeq: NC_005856 (P1 phage) as reference genomes. We provided the model with gene sequence plus symmetric context totaling 8,192 bp (equal distribution on both sides). For genes exceeding 8,192 bp in length, we utilized only the first 8,192 bp of the sequence.

We calculated scores by determining the difference in log-likelihoods between mutated and wildtype sequences. Our mutation strategy involved inserting multiple stop codons (“TAATAATAATAGTGA”) at a 12-nucleotide offset into the coding sequence. We evaluated the Evo 2 40B, Evo 2 7B, and Evo 1 131k models using this strategy. We also used the gene’s linear position in the reference genome as a predictive value for essentiality to control for potential positional bias in the model’s predictions. We assessed gene conservation as another control. We first extracted all protein sequences from the OpenGenome1 dataset, performed an all-by-all sequence search (using mmseqs easy-search with default parameters) between proteins and OpenGenome proteins, counted the number of significant hits (E-value threshold of 1 × 10^−2^), and used higher hit counts as an approximation of greater conservation and potential essentiality. We evaluated the predictive power of the log-likelihood changes (and control experiments) for binary gene essentiality using the AUROC score.

#### 4.3.11. lncRNA essentiality

Essential lncRNA genes were recently identified in a Cas13 knockdown experiment of lncRNA transcripts in 5 different cell lines (HAP1, HEK293FT, K562, MDA-MB-231, and THP1) (Liang et al., 2024). This included a total of 778 genes that were essential in one or more cell lines, and 46 that were essential in all 5 cell lines. Human lncRNA gene annotations were collected from a previous publication (Sarropoulos et al., 2019). Cas13 guide sequence binding sites were used as mutation positions, with 100 bp of sequence being scrambled around the genomic Cas13 mutation position, and up to 6,000 bp (Nucleotide Transformer) or 8,192 bp (Evo 2 models) of surrounding flanking sequence were extracted. 97.7% (48,310 out of 49,441) of Cas13 guide sequences were mapped to their corresponding lncRNA gene genomic position. The lncRNA transcripts were also scrambled at guide positions and analyzed separately. The difference in log-likelihoods for reference and mutated sequences were then calculated using Evo 2 7B, Evo 2 40B, and nucleotide transformer 2.5B multispecies. The average difference in log-likelihood for each gene was used as the final change in log-likelihood value for each gene. These values were then used as a predictive variable in a logistic regression model of gene essentiality, and directly compared to simple genetic metrics such as GC content and transcript length. Gene age values from the original lncRNA essentiality study (Sarropoulos et al., 2019) were used where available as an additional control.

#### 4.3.12. Zero-shot variant scoring

We compare different models’ ability to score mutations, zero-shot, by taking the delta between the predicted mutant and reference log likelihoods. DNA models use the DNA sequence around the variant while protein models are scored using the amino acid sequence of the gene. For indels, we also normalized to the reference sequence, and for DNA models to keep the change in likelihood invariant to length we maintain the total length of sequence scored to be the same regardless of the length of the indel by centering on the indel and adding or removing context nucleotides on the edges of the window. Variants assigned a more negative log-likelihood change from reference are considered to be more deleterious. Across the various evaluations, we used the following models for comparisons: CADD (Schubach et al., 2024), GPN-MSA (Benegas et al., 2023), PhyloP (Pollard et al., 2010), Nucleotide Transformer 2.5B multi-species (Dalla-Torre et al., 2024), Evo 1 (Nguyen et al., 2024a), RNA-FM (Shen et al., 2024), EnCodon/DeCodon (Naghipourfar et al., 2024), CodonBERT (Li et al., 2023), ESM-1b (Rives et al., 2021), ESM-2 (Lin et al., 2023), and AlphaMissense (Cheng et al., 2023).

To score a variant with Evo 2, we take a genomic window of length 8,192 around the variant and calculate the likelihood of the variant sequence divided by the likelihood of the reference sequence at the same position. For encoder models, we applied a position-wise masking to calculate the pseudolikelihood of the reference sequence normalized to pseudolikelihood of the reference alleles. We use the change in pseudolikelihood (Brandes et al., 2023) to score non-SNVs using encoder models such as ESM-1b, ESM-2, and Encodon, but cannot include indels for models with fixed coordinates like GPN-MSA or AlphaMissense. For PhyloP, we use PhyloP100 which is built on alignments from 100 vertebrate species. To score non-SNV variants, we take the total of the PhyloP scores at each affected position in the reference. For each benchmark we score the same variants with all models.

#### 4.3.13. ClinVar variant effect prediction

We use ClinVar release 2024.02.28, a database of expert annotated human disease variants (Landrum et al., 2013). We remove variants which affect more than 64 base pairs of reference or alternate allele sequence and remove variants of unknown significance from the evaluation. We include only variants on the nuclear genome, with a review status of at least two stars (i.e., at least multiple submitters provided evidence or an expert panel reviewed the variant), and subset to loci with matched transcript_ids in GRCh38.p14 GTF file.

Using model scores, we classify Pathogenic/Likely Pathogenic variants from Benign/Likely Benign variants, evaluating using AUROC and AUPRC. We calculate statistics for coding, noncoding, both for SNV and non-SNV, to enable comparison with specialized models that only support coding variants or only support scoring SNV.

#### 4.3.14. SpliceVarDB variant effect prediction

We used SpliceVarDB, a database of experimentally validated splice variants in humans (Sullivan et al., 2024), to classify mutations that cause aberrant splicing based on zero-shot prediction of various models. We excluded variants labeled as ‘low-frequency’ in SpliceVarDB.

#### 4.3.15. BRCA1/2 zero-shot classification

For all *BRCA1* and *BRCA2* SNVs with reported functional scores and classifications, we parsed sequences of a 8,192bp window around the variant site from the human reference genome used by the respective original studies (Findlay et al., 2018; Huang et al., 2025). For both genes, we used the classification of SNVs made in the original studies to label the SNVs: for *BRCA1*, SNVs labeled as “LOF” were classified as loss of function variants (*N* = 823), while SNVs labeled as “FUNC” or “INT” were labeled as functional/intermediate variants (*N* = 3,070); for *BRCA2*, SNVs labeled as “P strong”, “P moderate”, and “P supporting” were classified as loss of function variants (*N* = 1,156), while SNVs labeled as “B strong”, “B moderate”, “B supporting” were classified as functional/intermediate variants (*N* = 5,681).

#### 4.3.16. BRCA1 supervised classification

For all *BRCA1* SNVs, we again parsed sequences of a 8,192 bp window around the mutation site using the human reference genome. To identify the best block of Evo 2 40B to extract embeddings from for this task, we took embeddings from the pre-normalization layer of each block of the Evo 2 40B model for the reference sequence and the SNV. These embeddings were averaged across tokens to yield vectors of length 8,192 for the reference and the variant sequences. These two vectors were concatenated to yield vectors of length 16,384, which were used as inputs to train a classification model that consists of a feed-forward neural network with three hidden layers. The neural network takes an input of dimension *D* and processes it through fully connected layers of sizes 512, 128, and 32 neurons, respectively, before outputting a probability that the given SNV is pathogenic. Each hidden layer is followed by ReLU activation, batch normalization, and dropout (*p* = 0.3). The final layer uses sigmoid activation to produce binary classification probabilities. The complete architecture is:

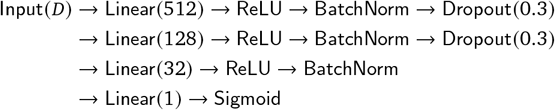

20% of all BRCA1 SNVs were withheld from training as the test set. 20% of the remaining SNVs were further sequestered as the validation set. The model was trained on the training set using Adam optimization (learning rate: 3 × 10^−4^, batch size: 128). Training employed early stopping (patience: 100), learning rate reduction on plateau (factor: 0.5, patience: 20, min_lr: 1×10^−6^), and gradient clipping (max norm: 1.0). The model was trained for up to 500 epochs using binary cross-entropy loss, with the best-performing model on validation data selected for final evaluation. The model trained with the embeddings from block 20 performed the best on the test set (AUROC = 0.92), which led us to choose this layer for the next step.

In the next step, for each SNV, we created a feature vector by extracting embeddings from four sequences: the reference sequence, its reverse complement, the variant sequence, and its reverse complement. For each of these four sequences, we focused on the narrower loci surrounding the variant site and calculated average embeddings across different window sizes (ranging from 16 to 8,192 nucleotides in powers of 2). We then concatenated the vectors from all four sequences into a single 32,768-dimensional feature vector for each SNV. These feature vectors were used to train a classification model using the same architecture and parameters as the model described previously. Among the different window sizes for averaging, we found that using a 128nt window produced the best results, achieving an AUROC of 0.95 on the test set. This is the model we use for comparison with zero-shot methods throughout our figures.

### 4.4. Mechanistic interpretability with sparse autoencoders

#### 4.4.1. SAE training and dataset composition

We trained a BatchTopK sparse autoencoder (Bussmann et al., 2024), a variant of the TopK sparse autoencoder (Makhzani and Frey, 2014; Gao et al., 2024a) on activations from the Evo 2 residual stream following layer 26 (a Hyena-MR block) for an intermediate checkpoint of the 7B model that was context-extended to 262,144 tokens. Sparse autoencoders take as input model activations *x* and autoencode them into a sparse feature vector *f*, before predicting a reconstruction of the inputs 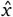. The conventional SAE architecture (which we also make use of here) is a very wide MLP with one hidden layer:

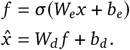

The nonlinearity σ (·) that we use is the Batch-TopK activation function. The Batch-TopK activation function is a version of the TopK activation function, which zeros out all but the *k* largest elements of a vector. The BatchTopK activation function with an equivalent value of *k* and a batch of size *B* zeros out all but the *kB* largest elements of the input batch. This allows the SAE to flexibly allocate capacity to higher-complexity inputs. Evo 2 activations have dimensionality *d*_model_ = 4, 096 and our SAE has feature dimensionality *d*_feature_ = 8 × 4, 096 = 32, 768. We used *k* = 64 for our SAE training.

Batch-TopK SAEs are trained with a loss ℒ= ℒ _recon_ +α ℒ _aux_ that combines a reconstruction loss ℒ _recon_ and uxiliary loss ℒ_aux_. The reconstruction loss simply the mean squared reconstruction error:

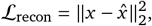

and the auxiliary loss ℒ _aux_ predicts the residual 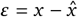 using only “dead features” (see Gao et al. (2024a) and Bussmann et al. (2024) for details). A dead feature is a feature *f*_*i*_ which has not fired (i.e., *f*_*i*_ = 0) for some large number of inputs during training. We call a feature dead if it has not fired for 10 million training inputs.

Our main SAE (which we refer to as the mixed-data SAE) was trained on a dataset of prokaryotic and eukaryotic genomes, representing a small subset of OpenGenome2. For prokaryotes, genomes were collected from GTDB release 220.0, filtering for genomes annotated as GTDB and NCBI representative genomes, and assembly level annotated as “complete”, totaling 2752 genomes and 10.97 billion base pairs. For eukaryotic genomes, 16 representative species were selected (see **Section 4.3.3**), and only regions with high gene content (genic regions, see **Section 4.2.2**) were used for SAE training, totaling 4.91 billion base pairs.

We subsampled this dataset to 1 billion activations from sequence chunks of length 16,384 to provide SAE training data. These activations were then globally shuffled so that activations from the same input sequence were unlikely to occur in the same SAE training batch. We trained the SAE using the Adam optimizer with a learning rate of 5 × 10^−5^, default values of β_1_ and β_2_, and a batch size of 16,384 for a single epoch. We used a trapezoidal learning rate schedule, ramping up from zero for the first 5% of training and ramping down to zero for the last 5%. We also trained an SAE solely on eukaryotic data (which we refer to as the eukaryotic data SAE). The eukaryotic data SAE was identically trained to the mixed-data SAE, except that instead of subsampling 1B tokens from the combined eukaryotic and prokaryotic datasets, we subsampled only from the eukaryotic dataset described above.

#### 4.4.2. SAE metrics and feature embedding calculation

To compute some basic statistics for the trained mixed-data SAE, activations were computed for all features across the *E. coli* K12 MG1655 genome (NCBI reference sequence NC_000913.3) and a length-matched segment of human chromosome 17 (NCBI reference sequence NC_000017.11, bases 40,019,967-44,661,618) in 50 kb non-overlapping sequence chunks. Activation density for each feature was then computed as the fraction of non-zero activations and the mean non-zero activation computed as expected across all sequence chunks from the relevant genome for both prokaryotes and eukaryotes. To visualize the features in an embedding, the (4096 × 32768) weights matrix was column-normalized and then embedded using UMAP (McInnes et al., 2018) with 2 components and random seed 1 and visualized by coloring each point by the difference in prokaryotic and eukaryotic activation density for each feature.

#### 4.4.3. Prophage feature

To identify a prophage-associated feature, we used contrastive feature search on 100 kb sequences that are centered on geNomad-annotated (Camargo et al., 2024) phage regions and include flanking bacterial regions. These sequences were from 100 randomly selected genomes from GTDB. The feature with the highest mean differential activation (f/19746) was selected. To evaluate the predictive value of this feature, we analyzed its mean activation values in prophage regions. We then compared these values against length-matched bacterial (non-phage) sequences from the same genome, sampled without replacement.

To analyze this feature at the genome scale, we computed feature activations on 50 kb non-overlapping chunks of the *E. coli* K12 MG1655 genome, and compared them with RefSeq-annotated phage regions. In the genome-scale plot in **Figure 4B**, we de-noise feature activations by setting a position’s activation to 0 if it does not have a neighboring position with a nonzero activation. From the dataset of 100 randomly selected GTDB genomes, we selected 3 loci with the highest f/19746 activation values outside of phage-annotated regions for visualization in **Figure S6D**.

As feature f/19746 also activated on the region downstream of the last CRISPR direct repeat in *E. coli*, we sought to determine whether Evo 2 is learning to associate sequences downstream of CRISPR direct repeats (like CRISPR spacers) with phage sequences or directly memorizing phage sequences. We investigated this by performing ablations on the sequence. Using the same CRISPR locus displayed in **Figure 4B**, we first computed feature activations on a synthetic sequence where all spacer sequences in the CRISPR array were independently scrambled. Observing that this did not result in a change in the feature activation pattern, we then computed feature activations on a synthetic sequence in which all the CRISPR direct repeats were replaced with a constant scrambled sequence, while keeping the spacer sequences unchanged from the natural sequence. Lastly, we repeated the previous test but replaced CRISPR direct repeats with nonconstant, scrambled sequences.

#### 4.4.4. E. coli genomic loci features

Features reflecting genome organization were identified using a combination of contrastive feature search and manual examination of features with large total activations over sequence chunks of the *E. coli* K12 MG1655 genome (NCBI reference sequence NC_000913.3), as annotated by RefSeq. Feature activations for the 100 kb segment displayed in the figures were computed over the full 100 kb segment as one batch.

To quantify mean activations over annotations, activations were computed over non-overlapping 50 kb chunks of the *E. coli* K12 MG1655 genome. Each nucleotide in the genome was grouped into ORF, tRNA, rRNA, or intergenic annotations based on existing RefSeq annotations, with the intergenic annotations including all sequences that were not categorized as ORF, tRNA, or rRNA. ORF annotations were further categorized into (+) ORF and (-) ORF depending on their directionality. For each annotation, mean activation for each feature was computed by extracting and averaging the activations across all nucleotides within the annotation.

#### 4.4.5 Protein secondary structure features

We identified α-helix and β-sheet associated features using contrastive feature search on all coding sequences in the *E. coli* K12 MG1655 genome. Secondary structure annotations were obtained using the DSSP algorithm (Kabsch and Sander, 1983; Kunzmann et al., 2023) on structures from the AlphaFold Protein Structure Database (Tunyasuvunakool et al., 2021). For each feature, we computed the Pearson correlation between its activation pattern on a coding sequence and the secondary structure annotations of the encoded protein, for all coding sequences. The features with the highest mean Pearson correlation with α-helix regions (f/28741) and β-sheet regions (f/22326) were selected.

Structures of EF-Tu in complex with thrT tRNA, and of RpoB and RpoC in complex, were obtained using AlphaFold 3 (Abramson et al., 2024) through the AlphaFold server. To project the features onto the structures, we first preprocessed the feature activation values. Except for thrT tRNA, we first mean collapsed feature activations per codon, as feature activations were computed at the genome level. Feature activations were then smoothed using a Gaussian kernel with a window size of 10. We then overlaid the smoothed feature activations on the structures, using linear interpolation coloring and clipping values at a maximum threshold of 0.2, visualized using ChimeraX (Meng et al., 2023).

#### 4.4.6. Frameshift and premature stop feature

Using the same set of artificially mutated coding sequences generated previously for the human genome, contrastive feature search was used with the eukaryotic data SAE to identify features that seemed to specifically correspond with frameshift and premature stop codon mutations, but not synonymous or nonsynonymous substitutions. A subset of 100 loci was first used to calculate average feature activations across all mutation types in 8192 bp windows. The difference in mean feature activations between the mutated and reference sequence was used to prioritize features. Top 20 features for each mutation type were identified. Features that were among the top 20 predictors for both frameshift and premature stop codon mutations were identified. Features that were also in the top 20 for synonymous or nonsynonymous substitutions were then excluded from this subset. This resulted in 3 remaining features: f/24278, f/29870, and f/18585. f/24278 was then selected for further analysis. A separate subset of 100 sequences, each of length 8192 bp, was used to analyze 24,278 feature activations on a per-nucleotide, and this subset was used to determine the average feature activation pattern near the mutation site. A separate subset of 500 sequences was then used to estimate the precision, recall, and the F1 score of the feature. For calculating these metrics, any non-zero activation of the feature within 100 bp after the mutation position was considered a true positive.

#### 4.4.7. Transcription factor motif features

From the GRCh38 human reference genome (Schneider et al., 2017), a random sample of 1,000 promoter sequences, defined as 1 kb upstream of a transcription start site, was selected and passed through the model, in batches of 100 sequences, to extract the SAE feature activations. All feature activations above the sparsity threshold of 1 ×10^−4^ were stored in sparse matrices from these activations. Following this selection, 31 bp windows centered on each position with a feature activation greater than the sparsity threshold were extracted.

To ensure robust motif detection, a selection pipeline for good features was established. First, features were required to have activations across at least 50 promoter sequences to be considered for further analysis. For each feature activation window, sequence complexity was assessed using Shannon entropy, calculated as *H* = −∑*π* log_2_ (*π*) where *π* represents the frequency of each nucleotide in the sequence. To ensure highquality motif detection, sequences below the 25th percentile of complexity scores were filtered out. Position Weight Matrices (PWMs) were constructed from the remaining sequences, and information content (IC) was calculated for each position as 2 *−H* of the nucleotide frequencies in the PWM. The motif scoring system incorporated multiple metrics: maximum IC, IC variance (to identify sharp conservation peaks), peak width (number of positions with IC > 0.5), and peak spacing. Penalties were applied for deviating from expected transcription factor binding site characteristics: a width penalty for diverging from the optimal 6-8 bp width, and a peak penalty for having significantly more or fewer than 1-2 distinct peaks. The final motif score was calculated as (maximum IC * 2 + IC variance) * (1 / (1 + width penalty + peak penalty)), with features requiring a maximum IC > 0.3 to be considered valid motifs. After this filtering, motifs were scored based on the maximum information content, number of high-information positions, information content variance, and sequence complexity. For some motifs, an additional scoring metric, the number of zeros appearing in the count matrices, was also incorporated into the scoring. Sequence logos were generated using the Logomaker package (Tareen and Kinney, 2019) for visual inspection, and the JASPAR count matrices were passed through the MEME suite TOMTOM Motif Comparison Tool (v5.5.7) (Gupta et al., 2007) with the “Human and Mouse (HOCOMOCO V12 CORE)” dataset (Vorontsov et al., 2023) to match the motifs to known transcription factor binding motifs.

#### 4.4.8. Exon-intron features

SAE features associated with eukaryotic coding sequences, introns, and exon-intron boundaries were found through contrastive feature search on the annotated sequence of chromosome 1 of the GRCh38 human reference genome (Schneider et al., 2017). 50 exon-rich segments of length 8,192 nt were sampled from this reference sequence, and feature activations were collected from each segment. Then, we searched for features with the top-20 highest values of the weighted sum

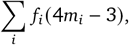

where *f*_*i*_ is the activation at base *i*, and *m*_*i*_ is 1 if base *i* is annotated as the given genomic component (CDS, intron, exon start, exon end), and 0 if not.

For features associated with exon starts and ends, the first 50 bases and the last 50 bases of each exon were used as the loci where *m*_*i*_ is 1, respectively. The features that are presented most frequently in these 50 segments were visually inspected to determine the features that activate the most consistently at the respective loci, and a representative feature was chosen for CDS, introns, exon starts, and exon ends. To calculate the F1, precision, and recall scores for each feature, we sampled 1,000 genes with 5 or more exons from the human genome, and counted the bases where the features have a nonzero activation on or off the genomic regions corresponding to each feature. The reference genome annotations for CDS and introns were considered to be ground truth in calculating the numbers of true/false positives/negatives. For the exon start feature (f/1050), we considered the first 25 bases of each exon following an intron to be the ground truth, in consideration of the firing patterns of the feature. For the exon end feature (f/25666), we considered the last one base of each exon followed by an intron to be the ground truth label. To calculate the mean activations of features for each type of genomic loci, we used the same set of 1,000 genes and calculated the average activations of each feature for each individual genomic locus identified by the ground truth labels – a single exon, a single intron, first 25 bases of a single exon, and the last base of a single exon.

The activations for the same set of features were collected for the *PDK3* gene loci of the woolly mammoth genome (Sandoval-Velasco et al., 2024). The exon coordinates of the mammoth genome were derived from homologs to the *Loxodonta africana* genome. Note that while the *Loxodonta africana* genomic sequence was part of the training corpus for Evo 2, it was not part of the training data used to collect activations for the SAE.

#### 4.4.9. SAE feature visualization web viewer

We manually curated 104 prokaryotic genomes of microbiological, medical, or agricultural relevance from NCBI. These genomes were annotated using the Bakta v1.10.3 pipeline (Schwengers et al., 2021), putative prohage regions were annotated using geNomad (Camargo et al., 2024), and secondary structure annotations were obtained using the DSSP algorithm (Kabsch and Sander, 1983; Kunzmann et al., 2023) on structures from the AlphaFold Protein Structure Database (Tunyasuvunakool et al., 2021). Activations were computed for all features without using the Batch-TopK activation function, and the top 50 largest activations per token were retained. Building on igv.js (Robinson et al., 2022), features identified as mapping to prokaryotic concepts are plotted and interactively displayed at https://arcinstitute.org/tools/evo/evo-mech-interp.

### 4.5. Unconstrained generation with Evo 2

#### 4.5.1. Gene completion

DNA sequences were collected around genes from *Haloferax volcanii, Escherichia coli, Saccharomyces cerevisiae, Chlamydomonas reinhardtii, Arabidopsis thaliana*, and *Homo sapiens*, covering archaeal, prokaryotic, and eukaryotic species. For each gene, a prompt of 1,000 base pairs upstream of the gene and the first 500 bp (*Haloferax volcanii, Escherichia coli, Saccharomyces cerevisiae*) or 1000 bp (*Chlamydomonas reinhardtii, Arabidopsis thaliana*, and *Homo sapiens*) of the gene sequence were input into the model. The language model generated the remaining sequence using a temperature of 0.7 and top-*k* of 4. We performed 10 generations for each prompt and evaluated results by converting to amino acids, aligning using BioPython, and calculating the percent protein recovery. We compared Evo 1 and Evo 1 131k with the different Evo 2 models.

#### 4.5.2. Mitochondrial genome generation

We prompted with 3,000 base pairs of the human mitochondrial genome and generated 250 unique mitochondrial sequences (16 kb each) using top-k of 4 and temperature of 1.0 or 0.7. Prompts were tested using positions 0-3,000 of the human mitochondrial genome and the reverse complement of positions 13,500–16,500. Generated sequences were annotated using MitoZ (Meng et al., 2019) to assess rRNA, tRNA, and CDS counts. Synteny analysis was performed using LoVis4u (Egorov and Atkinson, 2024). Sequence diversity was evaluated through BLAST searches against the nt/nr database to quantify query coverage and sequence identity. For structural analysis, we selected generated sequences containing the complete human complement of mitochondrial ND, CO, and ATP genes. Protein complexes were predicted using AlphaFold 3 through the AlphaFold server, with identical analysis performed on human mitochondrial genes for comparison. Generated structures were assessed through alignment against wild-type structures. Gene homology was analyzed using BLASTp with default settings, with percent sequence identity calculated as the product of query coverage and percent identity. Protein structures were visualized using ChimeraX (Meng et al., 2023).

#### 4.5.3. Mycoplasma genitalium genome generation

Generation was initiated using the first 10.5 kb of the reference *M. genitalium* genome as prompt. Using Evo 2 40B, we generated 35 sequences of 580 kb each with temperature 1.0 and top-*k* of 4. We compared these against previously generated *M. genitalium* sequences from Evo 1 131k and the reference natural *M. genitalium* genome. Protein-coding regions were predicted using Prodigal (Hyatt et al., 2010) and analyzed for homology using HHpred (Söding et al., 2005) against the Pfam database (Mistry et al., 2021). The density of significant hits (E-value < 0.001) was calculated for generations from each model. Proteins with significant Pfam hits were folded using ESMfold (Lin et al., 2023). Protein quality metrics evaluated included pLDDT scores, which were derived from ESMfold, and secondary structure distributions, which were obtained using DSSP (Kabsch and Sander, 1983). As a point of comparison, annotated proteins from the reference *M. genitalium* genome were folded using ESMFold and compared to those from Evo 2 generated *M. genitalium* using the same metrics. Protein structures were visualized using ChimeraX (Meng et al., 2023).

#### 4.5.4. Yeast chromosome generation

Sequences were generated using the first 10.5 kb of *S. cerevisiae* chromosome III as prompt, with temperature 1.0 and top-*k* of 4 settings in Evo 2 40B. Twenty artificial chromosomes of 330 kb were generated. Gene prediction was performed using GeneMark-S with Prot-hint based on the Fungi proteins from OrthoDB v12.0. As with the prokaryotic generations, predicted protein sequences were analyzed for homology against natural proteins using HHpred (E-value < 0.001) against the Pfam database. Proteins containing significant Pfam hits were subsequently folded using ESMFold and evaluated for pLDDT and secondary structure (DSSP). To predict tRNAs and promoters, generated DNA sequences were annotated using the Yeast Genome Annotation Pipeline (Proux-Wéra et al., 2012) and Promoter 2.0 (Knudsen, 1999) respectively (likelihood score > 0.8). Feature density analysis included quantification of predicted tRNAs, promoters, and genes with intronic structure. As a point of comparison, this process was repeated for the reference *S. cerevisiae* chromosome III genome.

### 4.6. Generative epigenomics via inference-time search

#### 4.6.1. Beam search algorithm

We implemented a beam search algorithm for efficiently sampling a sequence autoregressively while guided by a scoring function. At a high level, we sample autoregressively from a language model to obtain multiple chunks of tokens that are all continuations of the same prompt. We then use a given scoring function to select the best chunks, which are appended to the prompt for the next round of sampling.

More formally, let us denote a sequence as **x** = {*x*_1_, …, *x*^*L*^}∈ 𝒳 ^*L*^ where ^*L*^ is the sequence length and 𝒳 is the vocabulary (e.g., DNA base pairs). Let

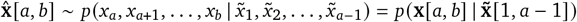

denote a sampled sequence from a distribution *p*, which we parameterize with an autoregressive language model (e.g., Evo 2), and where *a, b* ∈ [^*L*^] define the start and stop indices for a sampled sequence chunk such that *a* < *b*. We define *C* = *b −a* +1 as the length of a sampled sequence chunk. We can obtain samples from *p* via standard autoregressive decoding. On each iteration *t* of the beam search algorithm, we sample *K* chunks

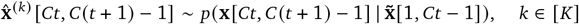

off the prompt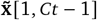.

Now, we are also given a scoring function *f* : 𝒳^*L*^ → ℝ_≥ 0_ that takes in a sequence and outputs a nonnegative score, where a lower value of the score is better. When we pick the prompt for iteration *t* + 1, we can pick the chunk with the lowest score to append to the prompt at iteration *t*, i.e.,

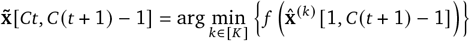

where

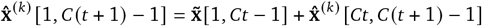

and + denotes a string concatenation operator.

Note that instead of taking only the best chunk, we can instead use the best *K*′ ≤ *K* chunks as prompts for sampling at the next iteration, i.e.,

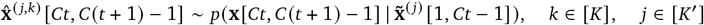

where 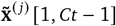 includes one of the best-*K*′ chunks (i.e., with the lowest scores) according to *f* obtained in the previous iteration. The algorithm iterates until all tokens ^*L*^ have been sampled. For the first chunk, we sample off a sequence 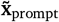. We assume that *C* is constant and that ^*L*^ is an integer multiple of *C*, i.e., all chunks sampled throughout the procedure, including the final chunk, are of the same length.

#### 4.6.2. Beam search for generative epigenomics

For the main design runs shown in **Figure 6E**, we used a chunk length *C* = 128, a total designed sequence length of ^*L*^ = 19, 968, sampled *K* = 42 chunks per prompt, and retained *K*′ = 2 prompts per iteration (i.e., we sampled 2 × 42 = 84 chunks per iteration). We used Evo 2 7B to parameterize *p*, from which we sample using standard autoregressive decoding with a temperature of 1.0 and top-*k* of 4. For 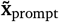, we used a sequence of length 40,960 upstream of the genomic region chrX: 52,051,929–52,123,468 in the mm39 reference genome.

For our scoring function, we used an ensemble of Enformer (Avsec et al., 2021) and Borzoi (Linder et al., 2025), where Borzoi itself contains an ensemble of four replicate models. We used the implementation of Enformer at https://github.com/lucidrains/enformer-pytorch and the Flashzoi reimplementation of Borzoi (Hingerl et al., 2024).

We note that Enformer takes in a sequence **x** ∈ 𝒳^196,608^ and outputs scores 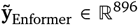, where each dimension of 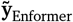 corresponds to 128 bp (representing output predictions for 896 × 128 = 114, 688 bp centered in the input), thereby not making predictions for 40,960 bp of left and right flanking context. Our scoring function requires a user-defined binary pattern **y**_Enformer_ ∈ {0, 1} ^896^, which specifies, for example, a

Morse code peak pattern. To score with Enformer, we simply divide all entries in 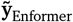 by the maximum value in 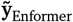 (i.e., so all values are in the interval [0, 1]), which we denote **ŷ**_Enformer_, and report the score as the *𝓁*_1_ norm

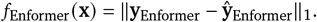

As input to Enformer, we concatenate: (i) 40,960 bp of sequence upstream of chrX: 52,051,929–52,123,468, (ii) the designed sequence, and (iii) enough DNA sequence downstream of chrX: 52,051,929–52,123,468 to achieve a total input sequence to Enformer of length 196,608. As beam search progresses, we mask the loss such that it is defined only on the sequence designed by Evo 2.

Borzoi makes predictions similar to Enformer but with longer sequence lengths and higher output resolution, taking in a sequence **x** ∈ 𝒳^524,288^ and outputting scores 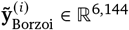, where *i* ∈ [4] refers to one Borzoi model replicate in an ensemble of four and each dimension corresponds to 32 bp (representing output predictions for 6, 144 × 32 = 196, 608 bp centered in the input), thereby not making predictions for 163,840 bp of left and right flanking context. We modify **y**_Enformer_ to have higher resolution by expanding its dimensionality to 6,144 by repeating each entry 128/32 = 4 times to produce a vector **y**_Borzoi_ ∈ {0, 1}^6,3 144^. We normalize the Borzoi predictions 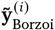 by dividing by the maximum value across all dimensions and across all predictions in the ensemble, to produce 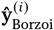. We then combine scores across the Borzoi ensemble by computing a “lower confidence bound”

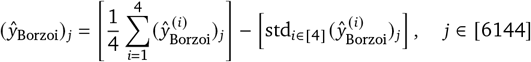

where 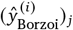 indicates the *j*th entry in 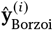 and std(·) is the standard deviation. Our Borzoi loss is then defined as

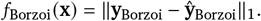

Analogous to the Enformer setting, our input sequence to Borzoi is a concatenation of: (i) 163,840 bp of sequence upstream of chrX: 52,051,929–52,123,468, (ii) the designed sequence, and (iii) enough DNA sequence downstream of chrX: 52,051,929–52,123,468 to achieve a total input sequence to Borzoi of length 524,288. As beam search progresses, we mask the loss such that it is defined only on the sequence designed by Evo 2

Our final scoring function simply takes the average of the Enformer and Borzoi scoring functions

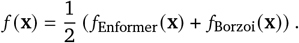

We use this scoring function to guide the beam search algorithm as described above.

#### 4.6.3. Design patterns and tasks

We designed six peak patterns:

- **Long square wave:** Alternating between 1664 bp of open chromatin and 1664 bp of closed chromatin.
- **Medium square wave:** Alternating between 768 bp of open chromatin and 768 of closed chromatin.
- **Short rectangular wave:** Alternating between 384 bp of open chromatin and 1152 bp of closed chromatin.
- **“LO” morse code:** Encoding LO `(.-.. ---)` in Morse code. One dot length corresponds to 768 bp.
- **“ARC” morse code:** Encoding ARC `(.- .-. -.-.)` in Morse code. One dot length corresponds to 384 bp.
- **“EVO2” morse code:** Encoding EVO2 `(. …- --- ..---)` in Morse code. One dot length corresponds to 384 bp.

We follow the Morse code specification in which a dash is three times the width of a dot, dots and dashes within a character are separated by a single dot length, and characters within a word are separated by three dot lengths. Dots and dashes are encoded by open chromatin; spaces are encoded by closed chromatin.

#### 4.6.4. Inference-time scaling experiments

We also conducted scaling analyses in which we vary the number of chunks sampled per beam-search iteration, thereby increasing the amount of inference-time compute required to complete a single design run. For each of the six patterns above, we ran beam search with the following configurations:

- *K*′ = 1, *K* = 1 (1 toK/bp)
- *K*′ = 1, *K* = 2 (2 toK/bp)
- *K*′ = 1, *K* = 3 (3 toK/bp)
- *K*′ = 1, *K* = 6 (6 toK/bp)
- *K*′ = 1, *K* = 9 (9 toK/bp)
- *K*′ = 1, *K* = 12 (12 toK/bp)
- *K*′ = 1, *K* = 15 (15 toK/bp)
- *K*′ = 2, *K* = 15 (30 toK/bp)
- *K*′ = 2, *K* = 30 (60 toK/bp)

where tok/bp indicates the number of tokens sampled per base pair designed. All other design parameters were kept the same. We assessed the quality of a design with the AUROC metric, in which the “true” values were the desired peak patterns **y**_Enformer_ and **y**_Borzoi_, the “predicted” values were the outputs **ŷ**_Enformer_ and **ŷ**_Borzoi_, and the AUROC compares the true and predicted values separately for Enformer and Borzoi. The final reported AUROC score for a design is the average of the Enformer and Borzoi AUROCs. We conducted these scaling laws using either Evo 2 7B or a model that samples uniformly over the nucleotide vocabulary as the proposal distribution *p*.

## 5. Data availability

The OpenGenome2 dataset used to train Evo 2 is available at:

https://huggingface.co/datasets/arcinstitute/opengenome2

## 6. Code and model availability

We make code and tools for model exploration available at the following links:

- Top-level code repository: https://github.com/arcinstitute/evo2
- Pretraining, midtraining and finetuning code: https://github.com/zymrael/savanna
- Inference code: https://github.com/zymrael/vortex
- Evo Designer, an interactive user interface for generation and scoring with Evo 2:
- https://arcinstitute.org/tools/evo/evo-designer
- Evo Mech Interp Visualizer, an interactive user interface for exploring SAE features:
- https://arcinstitute.org/tools/evo/evo-mech-interp
- NVIDIA Evo 2 NIM (generation): https://build.nvidia.com/nvidia/evo2-protein-design
- NVIDIA Evo 2 NIM (forward): https://build.nvidia.com/arc/evo2-40b
- NVIDIA BioNeMo version of Evo 2 code: https://github.com/NVIDIA/bionemo-framework

We make the model parameters available on Hugging Face:

- Evo 2 40B: https://huggingface.co/arcinstitute/evo2_40b
- Evo 2 7B: https://huggingface.co/arcinstitute/evo2_7b
- Evo 2 40B base: https://huggingface.co/arcinstitute/evo2_40b_base
- Evo 2 7B base: https://huggingface.co/arcinstitute/evo2_7b_base
- Evo 2 1B base: https://huggingface.co/arcinstitute/evo2_1b_base

### 7. Acknowledgments

We thank Euan Ashley, Joseph Caputo, Gabriel Filsinger, Michael Fischbach, Julia Kazaks, Anshul Kundaje, Milan Macek Jr., Scott Newins, Rachel Park, Sudarshan Pinglay, Brian Plosky, Alden Woodrow, Ben Viggiano, and Michael Zhang for helpful discussions and assistance with manuscript preparation. We thank the following people at NVIDIA for behind-the-scenes help: Dani Traphagen, Alissa Gordon, Haleh Lewis, Henry Estela, Timur Rvachov, Dong Ahn, Seungjun Nah, Christina Adams, Xiaowei Ren, Seonmyeong Bak, and Dennis Chang. The following people at NVIDIA provided additional support: Chris Dallago, Kyle Tretina, Johnny Israeli, Neha Tadimeti, Abe Stern, Dawn Voss, Eddie Calleja, Cory Ye, Rick Izzo, Maciej Bala, Stefania Alborghetti, Vipin Sirohi, Vikas Mehta, Pallab Bhattacharya, Jason Sewall, Alexandre Milesi, Dorota Toczydlowska, Jonathan Mitchell, Tim Moon, Viji Balas, Ashwath Aithal, Paniz Karbasi, Caroline Xuan, Glody Guo, Jared Wilber, Marshal Uhls, Matthew Harwood, Neel Patel, Ohad Mosafi, Risto Haukioja, Seth Poulos, and Sidney Bryson. G.B., A.T.M., and S.H.K. acknowledge funding support from the National Science Foundation Graduate Research Fellowship Program. D.B.L. acknowledges funding support from the Fannie and John Hertz Foundation. A.T.M. acknowledges funding support from the Knight-Hennessy Graduate Scholarship Fund. H.G. is an Arc Core Investigator and acknowledges funding support from Arc Institute. P.D.H. acknowledges funding support from Arc Institute, Yosemite, Rainwater Foundation, Curci Foundation, Rose Hill Innovators Program, V. and N. Khosla, S. Altman, and anonymous gifts to the Hsu Lab. B.L.H. acknowledges funding support from Arc Institute, Stanford Institute for Human-Centered Artificial Intelligence (HAI) Hoffman-Yee Research Grants, V. Gupta, and R. Tonsing.

## 8. Author contributions

P.D.H. and B.L.H. conceived the project. D.P.B., H.G., P.D.H., and B.L.H. supervised the project. M.P. and E.N. designed the model architecture. M.P. developed the first version of the training code. G.Bri., J.K., G.Bro., D.W.R., E.N., A.T., B.Y., and B.L.H. contributed to the training code. M.P. developed the first version of the inference code. G.Bri., A.V., and B.Y. contributed to the inference code. G.Bri., M.G.D., and B.L.H. curated and processed the pretraining dataset. G.Bri. conducted training data composition and loss experiments. J.K., M.P., and E.N. conducted model pretraining. G.Bri., J.K., and M.P. conducted model midtraining and context extension. M.G.D., J.K., and B.L.H. developed and conducted needle-in-a-haystack and other midtraining evaluations. M.G.D., G.S., A.X.L., K.Z., and B.L.H. performed zero-shot mutational effect prediction. H.G. performed zero-shot human mRNA decay rate prediction. M.G.D. and J.C.S. conceived and implemented the exon/intron classifier. G.Bri. and M.N. performed zero-shot human variant effect prediction. G.G. and G.S. performed zero-shot and supervised *BRCA1/2* analyses. M.G.D., D.C., and D.B.L. curated and processed SAE training datasets. L.G. and T.M. implemented and trained SAEs. M.D., N.N., N.K.W., D.B., E.H., and T.M. developed the SAE feature visualization tool. M.G.D., D.C., G.G., D.B.L., G.S., L.G., N.K.W., D.B., P.D.H., and B.L.H. performed SAE feature discovery. A.T.M. conducted short-sequence prompt completion experiments. G.Bri., S.H.K., and D.G. sampled and analyzed mitochondrial genomes. G.Bri., S.H.K., and A.T.M. sampled and analyzed *M. genitalium* genomes. G.Bri., S.H.K., and A.T.M. sampled and analyzed yeast genomes. B.L.H. implemented the beam search guidance algorithm. B.L.H. conducted the generative epigenomics design runs and scaling analyses. G.Bri., E.A., R.I., R.M., J.S., and D.P.B. designed and implemented the Evo Designer tool. S.D., M.Y.N., J.P., and T.H.-B. conducted the safety, security, and ethics investigation. G.Bri., M.G.D., J.K., M.P., D.C., G.G., S.H.K., D.B.L., A.T.M., M.N., C.R.-T., G.S., H.G., P.D.H., and B.L.H. wrote the first draft of the manuscript. All authors wrote the final draft of the manuscript.

## Supplementary Material

### A. Supplementary figures

**Figure S1.**
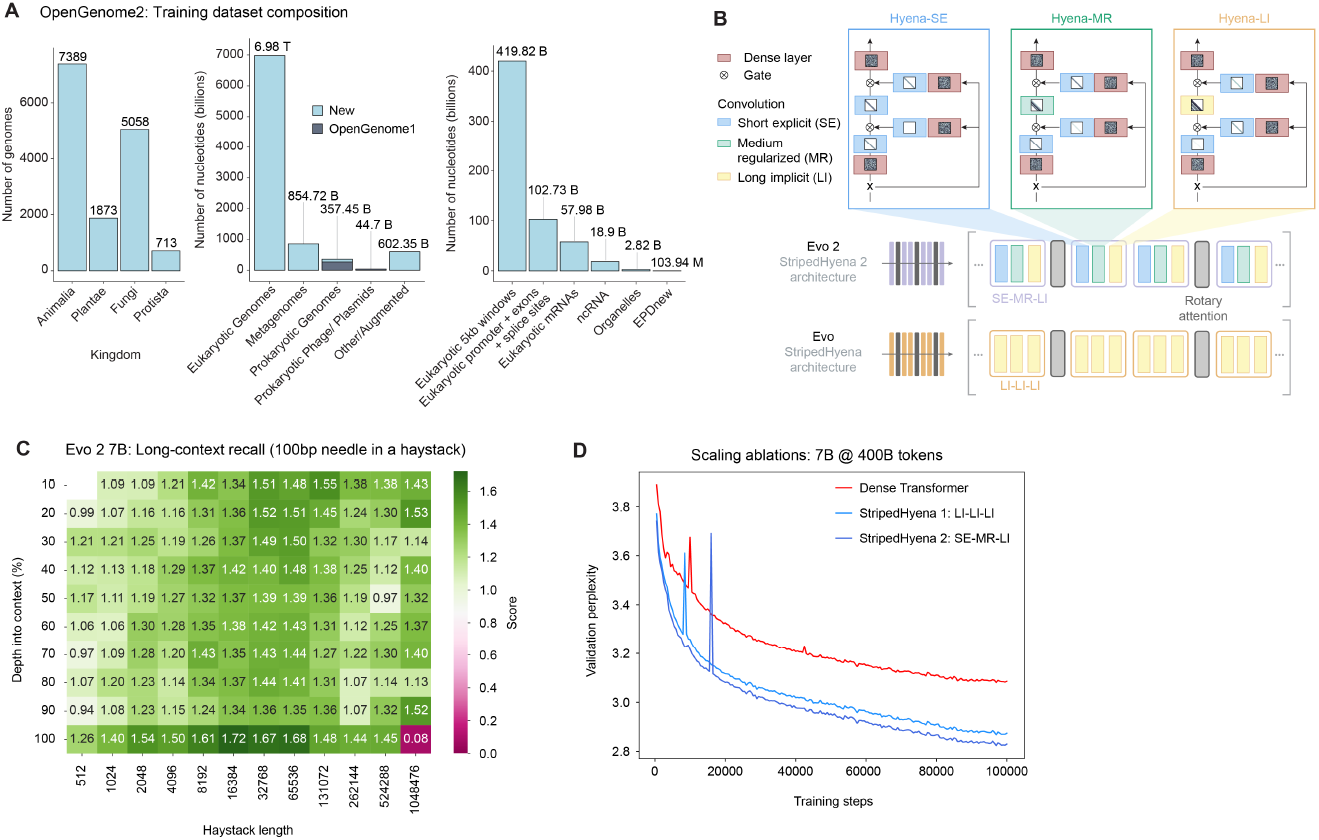
Overview of model architecture, training procedure, datasets, and evaluations for Evo 2. (**A**) Data composition of OpenGenome2. Showing total eukaryotic genomes per kingdom (left), total base pairs per training data subset (middle), and detailed breakdown of other/augmented training data subset (right). (**B**) Core input-dependent convolution operators in StripedHyena 2, with a diagram showing their composition in the architecture. (**C**) Needle-in-a-haystack performance of Evo 2 7B, spanning input contexts of 512 to 1 million tokens. (**D**) Scaling ablations on OpenGenome2, showing the loss convergence of multi-hybrids compared to previous generation hybrids and transformers. Models of 7 billion parameters are compared after pretraining with the same 400 billion tokens.

**Figure S2.**
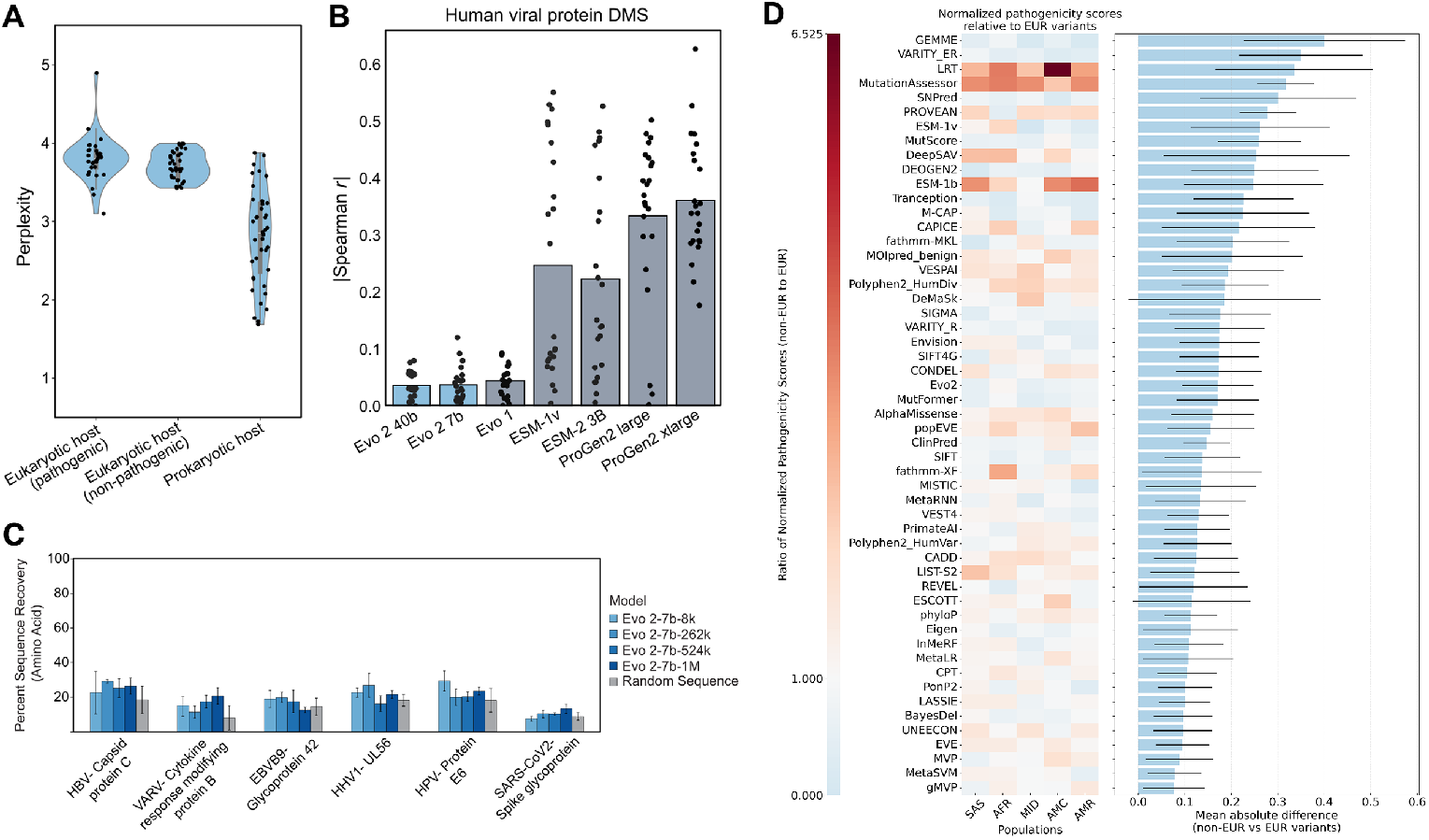
Biosecurity and ethics evaluations for model generation and scoring. To decrease dual use risks, safety filtering was performed on the training data to remove viral sequences that can infect eukaryotic hosts. Evo 2 is less performant on eukaryotic viruses, as intended. (**A**) Perplexity scores for viral sequences from the USDA Select Agents and Toxins List consistently demonstrate elevated perplexity values compared to non-pathogenic viruses and prokaryotic viruses. Blue violin plots show the distribution of scores, with individual data points overlaid representing 512-bp chunks sampled uniformly at random across viral genomes. (**B**) Correlation of language model likelihood with experimental deep mutational scanning (DMS) fitness measurements for human viral proteins. Gray bars represent mean correlation coefficients, with individual data points corresponding to DMS datasets from ProteinGym. Results indicate poor predictive capability on viral protein mutational effects for Evo 2 and Evo 1 models. (**C**) Comparative analysis of protein sequence generation success rates across different model conditions. Bar heights represent percentage amino-acid sequence recovery in the response sequences when prompted with a portion of a viral protein, with error bars showing standard deviation across multiple responses to the same prompt. Models were tested with various prompting proteins (shown on the horizonal axis) with different Evo 2 models (indicated by color). Random sequence generations are included as a control condition. (**D**) Analysis of ancestry bias for Evo 2 as a variant effect predictor compared to baselines, with protein mutations converted to DNA codons. Baseline performance data is taken from Pathak et al. (2024). Most variant effect predictors have ancestry bias, and score non-European ancestry variants as more pathogenic. Evo 2 has similar ancestry bias as other population-free methods, examined by taking both the ratio (heatmap) and mean difference (barplot) of min-max scaled scores of each population subgroup to the European subgroup.

**Figure S3.**
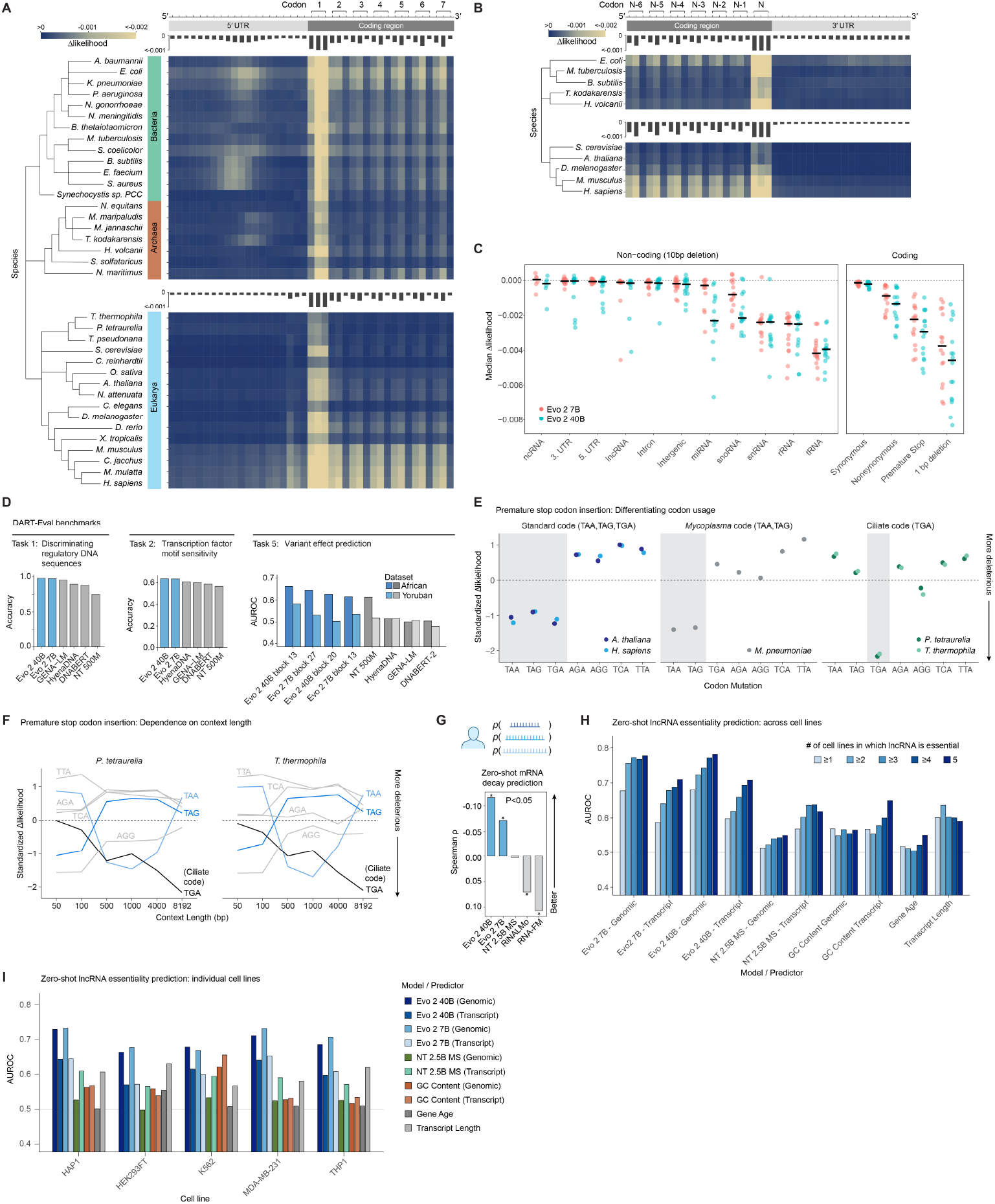
Evo 2 understands mutational effects on protein, RNA, and organismal fitness across all domains of life. *(caption continued on next page)* (**A**) Evo 2 predicts mutations to be unlikely in the start codons of protein-coding genes, the first two bases of each codon of the coding region, and the ribosome-binding sites of the 5′ UTR, across 20 prokaryotic and 16 eukaryotic model species. (**B**) Evo 2 predicts mutations to be unlikely in the stop codons of protein-coding genes and the first two bases of each codon of the coding region before the stop codon. (**C**) Evo 2 40B predicts lower likelihoods for deletions in miRNA and snoRNA loci compared to Evo 2 7B. Red points in (**C**) are the same as is shown in **Figure 2D**. The same sequences were analyzed with both models. (**D**) DART-Eval results for three tasks focused on regulatory DNA sequence elements. Task 1 evaluates models on their ability to distinguish candidate cis-regulatory elements (cCREs) from shuffled sequences, Task 2 tests their ability to identify transcription factor (TF) motifs by distinguishing true TF binding sites from control sequences, and Task 5 predicts variant effects for chromatin-accessibility QTLs (caQTLs, African) and dynamic sequence QTLs (dsQTLs, Yoruban). Sequence likelihoods were computed under each model and used to measure classification accuracy. Across these tasks, Evo 2 7B and Evo 2 40B outperformed other baselines. For task 5, while we did not see strong signal across all models when using the zero-shot log-likelihoods, we observed that the Evo 2 embeddings were predictive of noncoding variant effects. (**E**) Evo 2 differentiates between genomic sequences of model organisms that use different stop codons. Show z-score standardized median Δlikelihood values across 5 species, median calculated across ~ 4,100 randomly selected mutation loci. (**F**) Evo 2 requires >4kb of sequence context to recognize codon tables. Showing median z-score standardized median Δlikelihood values for two ciliate genomes across 6 sequence context lengths. (**G**) Length-adjusted Evo 2 likelihoods of human mRNA sequences showed a negative correlation with their experimentally measured decay rates. (**H**) Evo 2 predictions for lncRNA essentiality improve for lncRNAs that are essential in multiple cell lines. Comparing effect of scrambled mutations in non-essential genes (*N* = 5,417), and genes that are essential in ≥ 1 (*N* = 778), ≥ 2 (*N* = 301), ≥ 3 (*N* = 164), ≥ 4 (*N* = 93), and 5 (*N* = 46) cell lines. (**I**) Predictions for lncRNA essentiality across different models/predictors, shown for each individual cell line. Comparing effect of scrambled mutations in non-essential genes (*N* = 5,417) versus essential genes in HAP1 (*N* = 283), HEK293FT (*N* = 267), K562 (*N* = 278), MDA-MB-231 (*N* = 270), and THP1 (*N* = 284).

**Figure S4.**
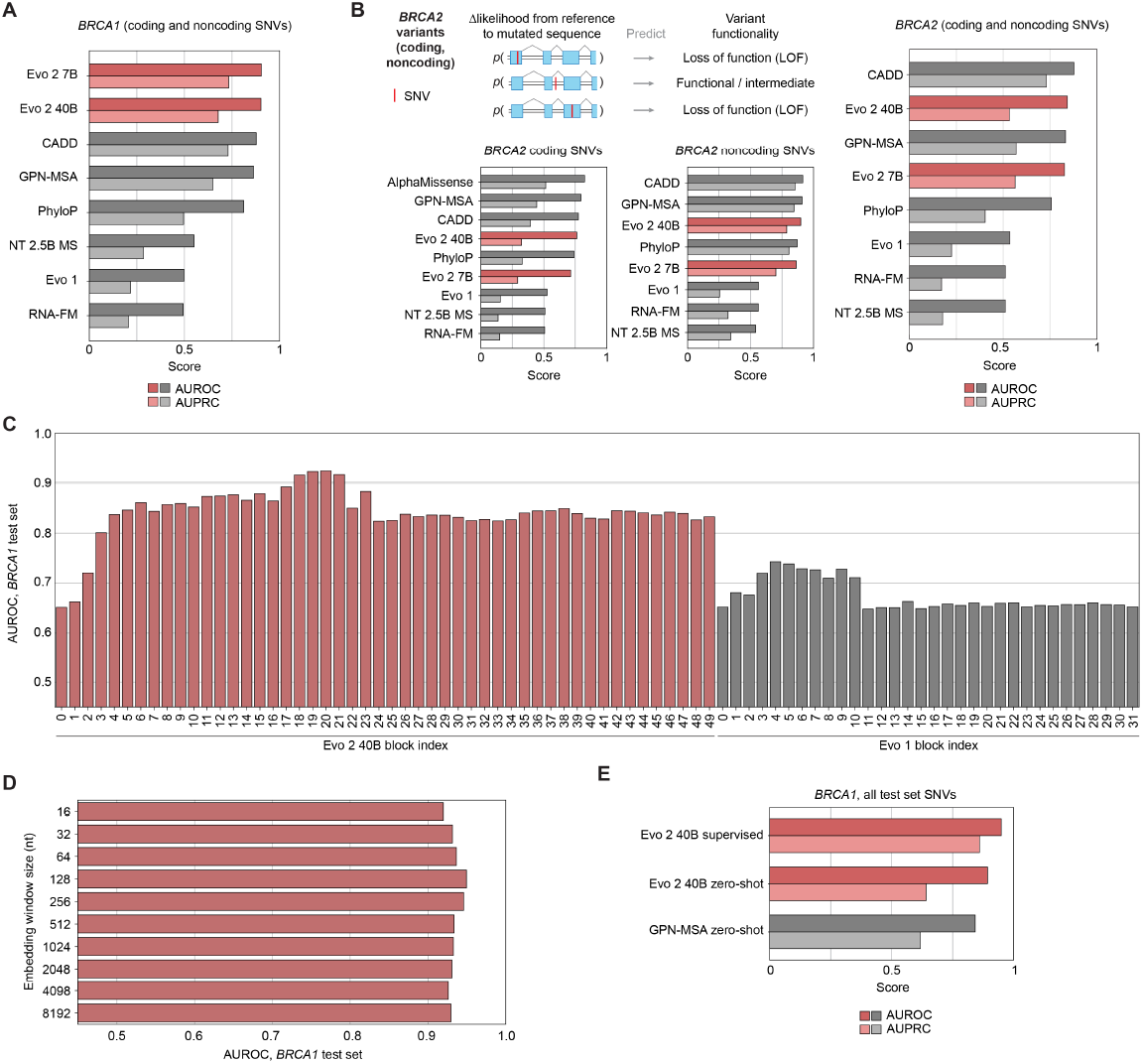
Evo 2 enables accurate human clinical variant effect prediction. (**A**) Zero-shot prediction of *BRCA1* variant pathogenicity for coding and noncoding SNVs evaluated in aggregate (*N* = 3,893), showing AUROC and AUPRC scores across models. (**B**) Zero-shot prediction of *BRCA2* variant pathogenicity for coding (left, *N*=3,446), noncoding (center, *N* = 506), and aggregated (right, *N* = 5,219) SNVs, showing AUROC and AUPRC scores across models. (**C**) Comparison of AUROCs for the supervised classification of test set *BRCA1* SNVs trained with embeddings from different blocks of Evo 2 40B and Evo 1. The best-performing layer (block 20 of Evo 2) was chosen for the next step. (**D**) Comparison of AUROCs for the supervised classification of test set *BRCA1* SNVs, using different window sizes around the variant site to average embeddings for. (**E**) Comparison of supervised Evo 2 model predictions to zero-shot Evo 2 and GPN-MSA baselines for all test set SNVs (*N* = 779).

**Figure S5.**
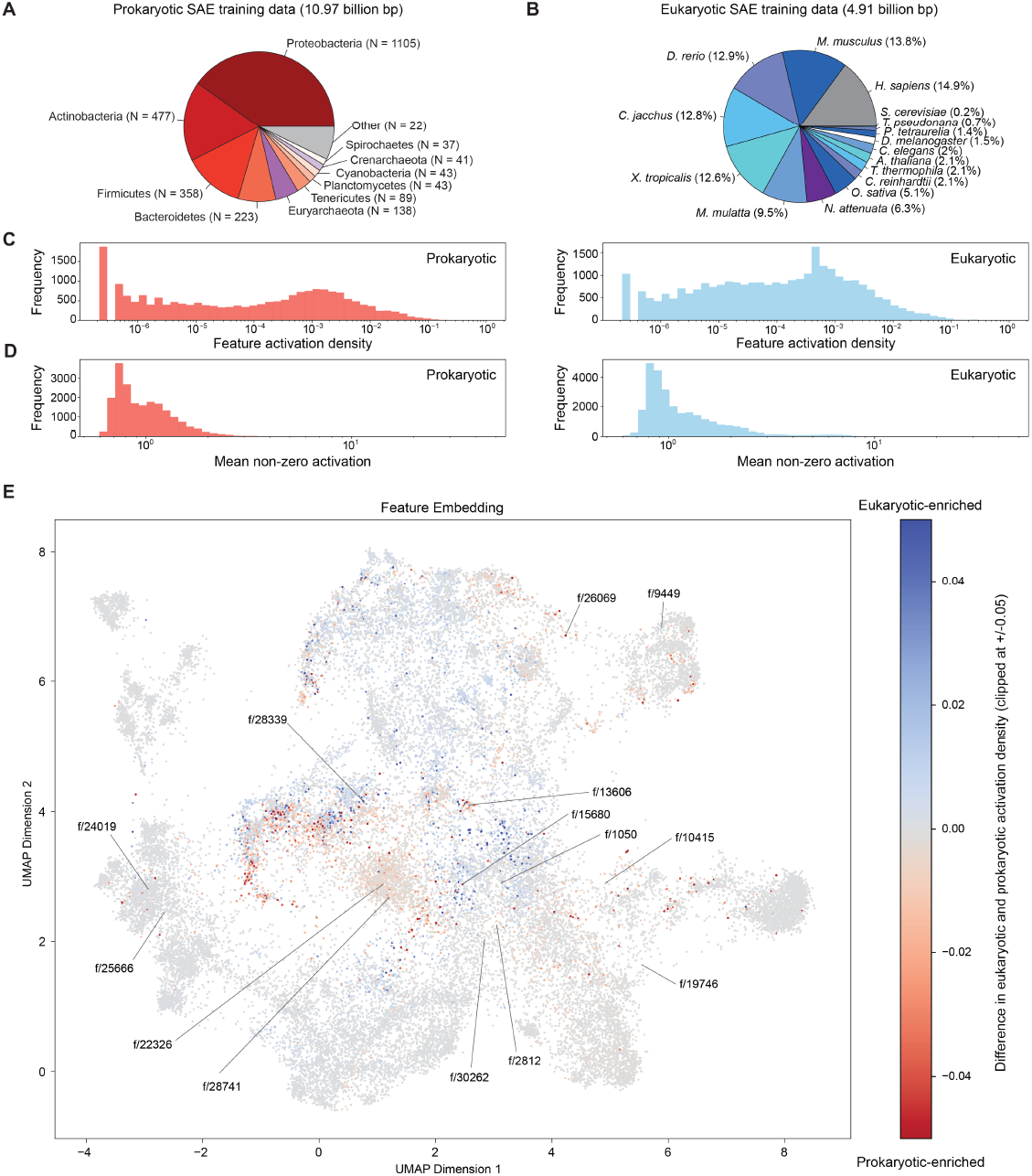
SAE overview with training data, metrics, and feature embedding. (**A**) Composition of the prokaryotic sequences randomly subsampled for SAE training. (**B**) Composition of the eukaryotic sequences randomly subsampled for SAE training. (**C**) Activation density for all layer 26 SAE features over the *E. coli* K12 MG1655 genome (left) or for a length-matched segment of human chromosome 17 (right). (**D**) Mean non-zero activation for all layer 26 SAE features over the *E. coli* K12 MG1655 genome (left) or for a length-matched segment of human chromosome 17 (right). (**E**) UMAP embedding of layer 26 SAE feature weights colored by activation density difference between eukaryote and prokaryotic sequence for each feature, with features presented in **Figure 4** labeled.

**Figure S6.**
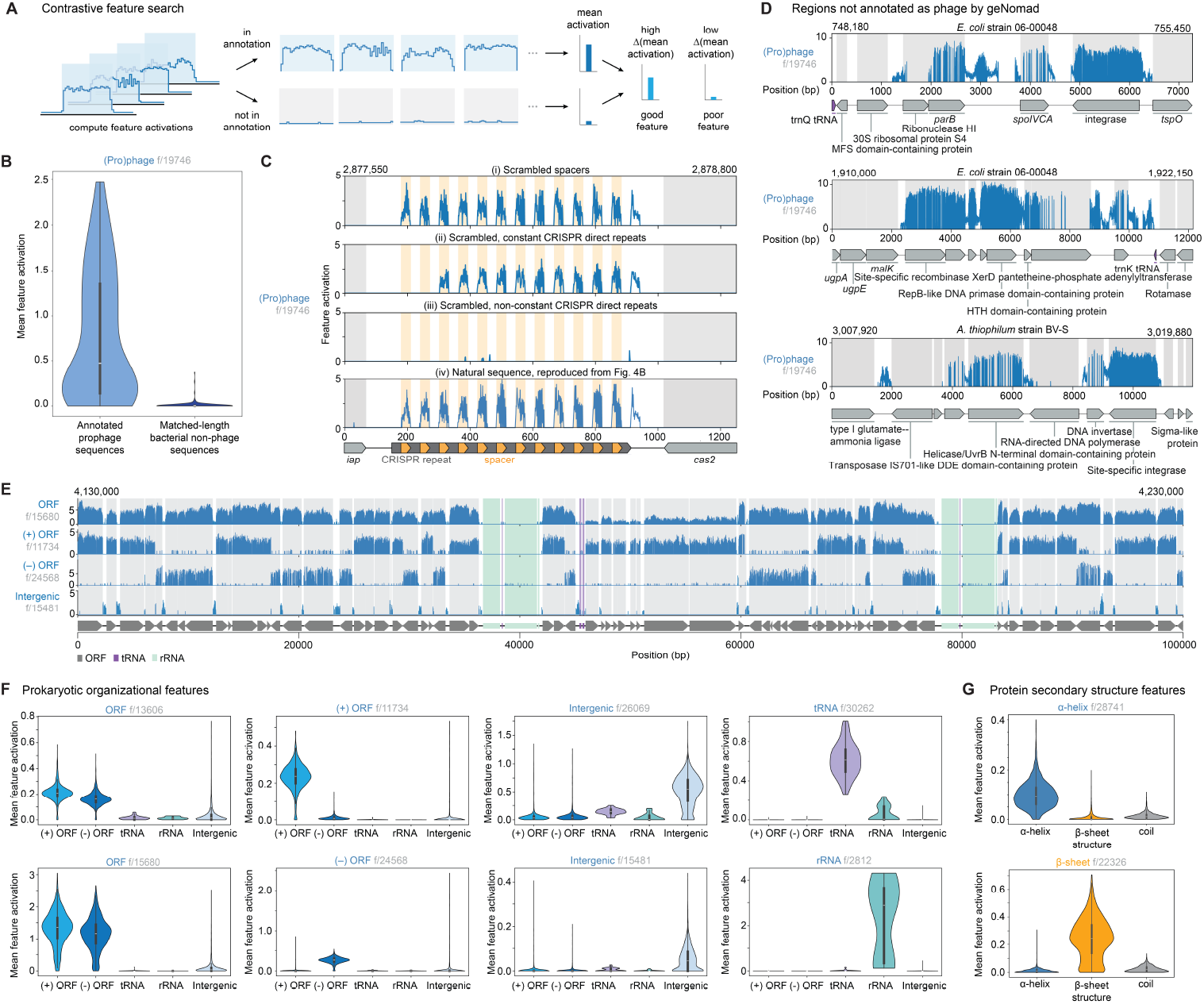
SAE features reveal semantic, structural, and organizational details of prokaryotic genomes. Diagram depicting the contrastive feature search strategy used to identify and quantify selected features. Mean activations of the phage feature across phage regions annotated by geNomad and matched-length bacterial non-phage sequences, across 100 randomly selected GTDB genomes. (**C**) (i) Scrambling spacer sequences does not ablate the phage feature activation pattern on spacer sequences. (ii) Using a constant scrambled CRISPR direct repeat sequence ablates phage feature activation for the first two spacers. (iii) Using different scrambled sequences instead of CRISPR direct repeats ablates the phage feature activation pattern. Natural sequence activation pattern, as in **Figure 4B**. (**D**) Additional examples of sequences not annotated as phage sequences by geNomad which the phage feature activates on. (**E**) Activations of additional features associated with open reading frames (ORFs), plus strand or minus strand ORFs ((+) ORF and (-) ORF), and intergenic loci in a 100 kb region in *E. coli* K12 MG1655. (**F**) Mean activations for prokaryotic organizational features on different annotation types across the *E. coli* K12 MG1655 genome. (**G**) Mean activations for protein secondary structure features on different secondary structure types across ORFs in the *E. coli* K12 MG1655 genome.

**Figure S7.**
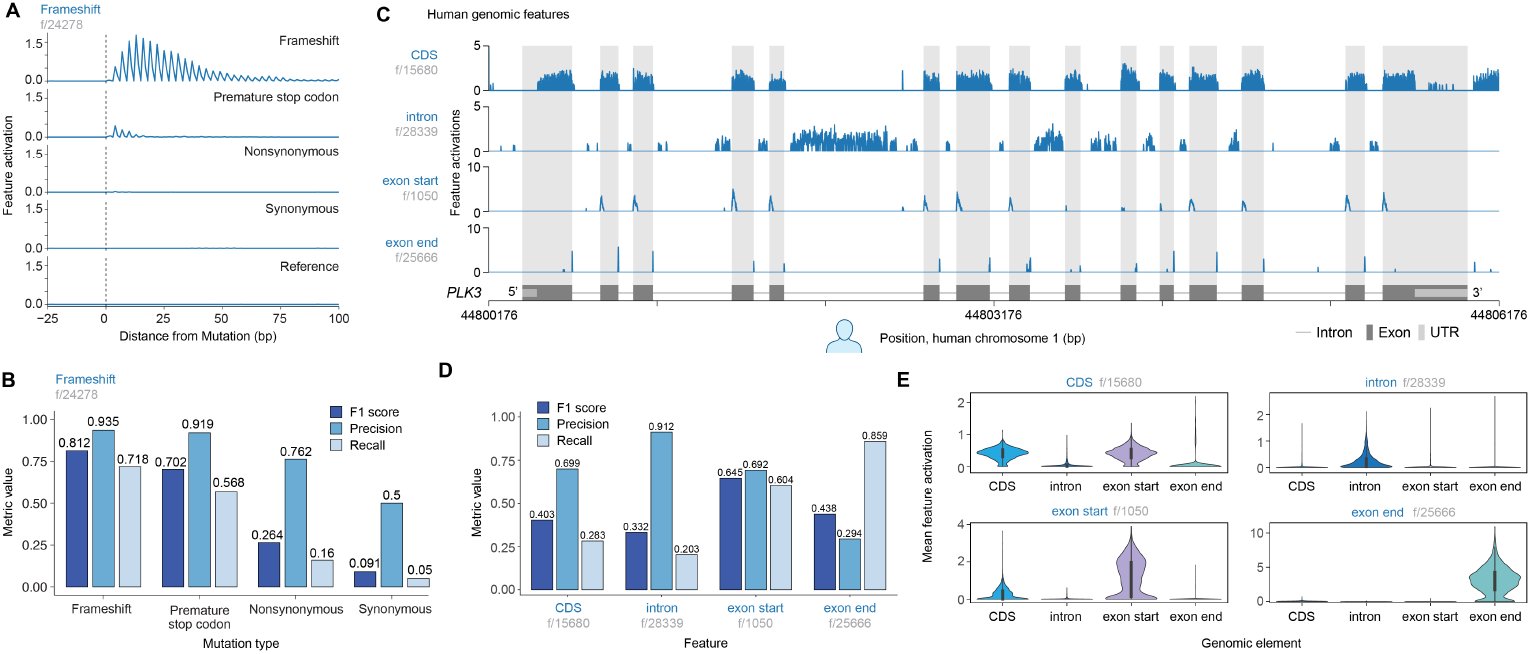
SAE features reveal semantic, structural, and organizational details of eukaryotic genomes. Activations of a frameshift-associated feature in a 100 bp region following different mutation types. (**B**) F1, precision, and recall scores across mutation types for feature shown in (**A**). (**C**) Activations for SAE features associated with exons, introns, and their boundaries in the human genome, shown for a 6000 bp region in chromosome 1. (**D**) F1, precision, and recall scores for each SAE feature shown in (**C**) to its corresponding genomic element. (**E**) Mean activations for each SAE feature shown in (**C**) on different annotation types across the human genome.

**Figure S8.**
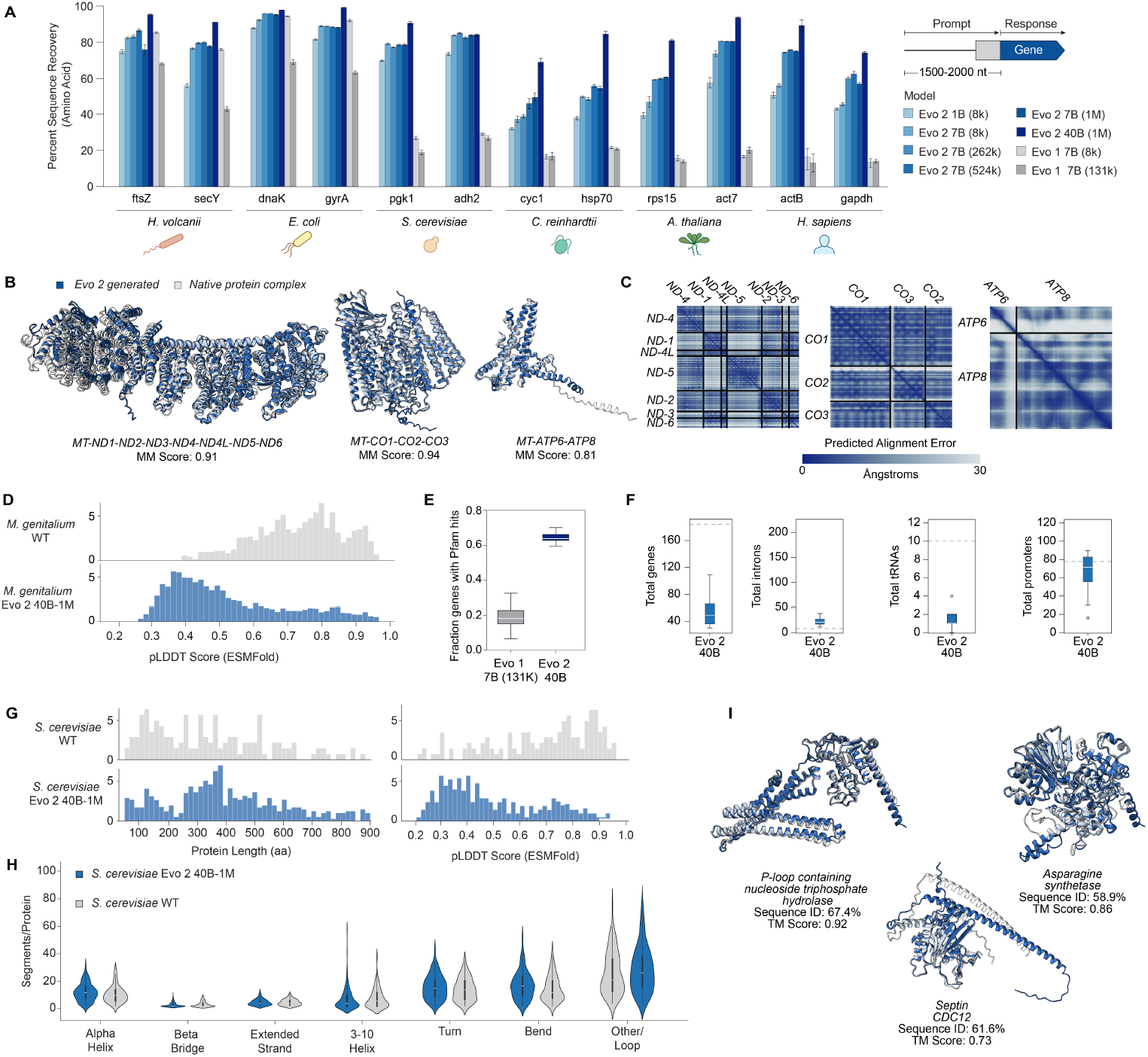
Additional results for unconstrained generation at genome and chromosome scale. (**A**) Evaluating the amino acid recovery for different genes across Evo 2 models. (**B**) AlphaFold 3 complexes of native human mitochondria and of Evo 2 generated human, aligned together. (**C**) PAE of the Evo 2 generated mitochondrial complexes from AlphaFold 3. (**D**) ESMFold pLDDT distributions of natural and Evo 2 generated *M. genitalium* genes called by Prodigal. (**E**) Fraction of Prodigal annotated genes that have Pfam hits comparing Evo 2 40B with Evo 1 7B-131K *M. genitalium* generations. (**F**) Distribution of genes, introns, tRNAs, and promoters on Evo 2 generated *S. cerevisiae* chromosome compared with natural (gray line). (**G**) Distribution of gene length and pLDDT of Evo 2 generated and natural *S. cerevisiae* genes, annotated by GeneMark-ES. (**H**) Distribution of different secondary structures in Evo 2 generated and *S. cerevisiae* chromosome III wild type genes. (**I**) Protein structure of genes from Evo 2 generated *S. cerevisiae* chromosome compared with the wild type gene structure and sequence.

**Figure S9.**
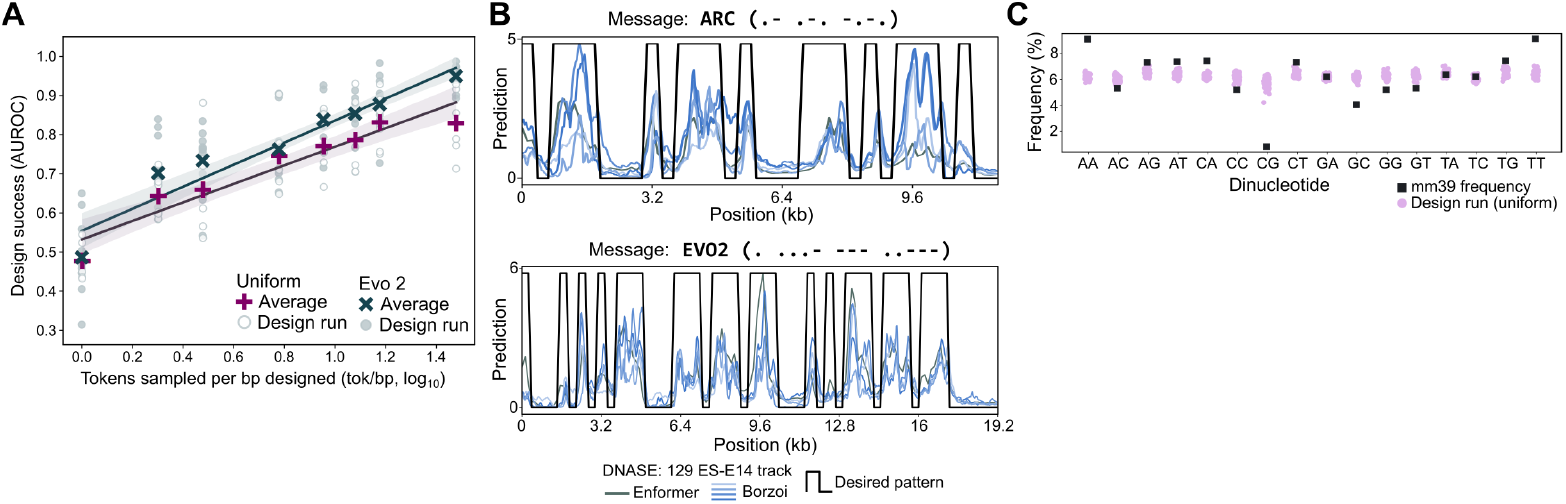
Generative epigenomics with a uniform proposal. (**A**) Token-matched performance has worse inference-time scaling when using a random sequence proposal compared to Evo 2-generated sequences, as shown across varying token sampling rates. (**B**) Reduced concordance between Enformer and Borzoi predictions suggests exploration of potentially adversarial model regimes, demonstrated for two representative Morse code designs (“ARC” and “EVO2”). (**C**) Dinucleotide frequencies in randomly proposed sequences after beam search filtering with Enformer and Borzoi still show significant deviation from the baseline mm39 frequency.

## B. Appendix

### B.1. Repeat Down Weighting Ablation

We conducted experiments to assess the impact of repeat loss reweighting on downstream performance using two Striped Hyena 1 models (550M parameters, 8192 sequence length). Both models were trained on a differently weighted ablation dataset, which was enriched for complete eukaryotic genomes to better evaluate the effects of repeat down-weighting (**Table S5**). The models differed only in their treatment of repetitive sequences: one applied a loss reweighting factor of 0.1 to lowercase sequences, while the other used no reweighting.

Performance was evaluated using the ClinVar pathogenic versus benign classification benchmark. The model without loss reweighting achieved an AUROC of 0.63 at 40,000 training steps, compared to 0.73 for the model with reweighting. Training of the non-reweighted model was discontinued after we observed no improvement in ClinVar performance beyond 30,000 steps, while the reweighted model continued to show improvements until the experiment concluded at 100,000 steps, reaching 0.82 AUROC. Based on these results, we adopted a loss reweighting factor of 0.1 for repetitive sequences in subsequent experiments.

**Table S1.** ClinVar AUROC at 40,000 Steps.

| Model | AUROC |
| --- | --- |
| No reweighting | 0.63 |
| Repeat Reweighting 0.1 | 0.73 |

#### B.2. Data Composition Effect on Downstream Tasks

We compare Evo 2 7B base with a 7B parameter StripedHyena 2 model trained on an ablation data composition (**Table S5**) at 8192 context length. This ablation dataset includes mRNA with the same weight as the final pretraining dataset, but used whole genomes instead of the eukaryotic genomic windows and augmented mRNA. We trained for 1.9T tokens until the loss plateaued. We compare the data ablation model with Evo 2 7B base model on zero-shot prediction of ClinVar, SpliceVarDB, and *BRCA1*, following the same protocol as before for these evaluations (**Section 4.3.12**).

Zero-shot evaluations demonstrated improved performance for the final pretraining data composition compared to the data ablation, suggesting that focusing pretraining on functionally enriched regions around genes can significantly improve downstream performance. Zero-shot AUROC improves for *BRCA1* variants from 0.793 to 0.891, ClinVar SNV from 0.933 to 0.957, ClinVar non-SNV from 0.897 to 0.939, and SpliceVarDB (intronic and exonic) from 0.791 to 0.826. We improvements for all subsets except for ClinVar coding SNVs, which slightly decrease. While the data ablation has more weight to noncoding regions, the high effect non-coding variants considered by these benchmarks are better modeled by the Evo 2 7B base model which was pretrained on datasets focused around genes. Importantly, the final model learned to better calibrate these proximal non-coding and coding effects as shown by the improvements when considering coding and non-coding together. At the same time, Evo 2 7B base performs within 0.01 at DART-eval zero shot evaluations of the data ablation 7B model (task 1: 0.97 vs 0.98, task 2: 0.63 vs 0.64), suggesting that added pretraining on whole genomes did not help substantially even for tasks on enhancer regions, consistent with findings from recent analysis of DNA language models (Patel et al., 2024). These results highlight the importance of data engineering in DNA language models.

#### B.3. Evo 2 Mitochondrial Generation BLASTp

We analyze the generated mitochondrial genes of the generated mitochondria shown in **Figure 5F** using BLASTp with default parameters and find alignments to a diversity of different animal species.

**Table S2.** Comparing Evo 2 7B base and data ablation model on zero-shot ClinVar classification prediction.

| Metric | Subset | 7B data ablation | Evo 2 7B base | Difference |
| --- | --- | --- | --- | --- |
| AUROC | All SNV | 0.933 | 0.957 | +0.024 |
| AUPRC | All SNV | 0.910 | 0.927 | +0.017 |
| AUROC | All non-SNV | 0.897 | 0.939 | +0.042 |
| AUPRC | All non-SNV | 0.846 | 0.898 | +0.052 |
| AUROC | Coding SNV | 0.851 | 0.829 | -0.022 |
| AUPRC | Coding SNV | 0.908 | 0.883 | -0.025 |
| AUROC | Coding non-SNV | 0.893 | 0.918 | +0.025 |
| AUPRC | Coding non-SNV | 0.946 | 0.954 | +0.008 |
| AUROC | Noncoding SNV | 0.943 | 0.976 | +0.033 |
| AUPRC | Noncoding SNV | 0.911 | 0.956 | +0.045 |
| AUROC | Noncoding non-SNV | 0.867 | 0.929 | +0.062 |
| AUPRC | Noncoding non-SNV | 0.726 | 0.825 | +0.099 |

**Table S3.** Comparing Evo 2 7B base and data ablation model on zero-shot SpliceVarDB classification.

| Metric | Subset | 7B data ablation | Evo 2 7B base | Difference |
| --- | --- | --- | --- | --- |
| AUROC | All | 0.791 | 0.826 | +0.035 |
| AUPRC | All | 0.845 | 0.873 | +0.028 |
| AUROC | Exonic | 0.666 | 0.687 | +0.021 |
| AUPRC | Exonic | 0.510 | 0.540 | +0.030 |
| AUROC | Intronic | 0.893 | 0.923 | +0.030 |
| AUPRC | Intronic | 0.956 | 0.968 | +0.012 |

**Table S4.**
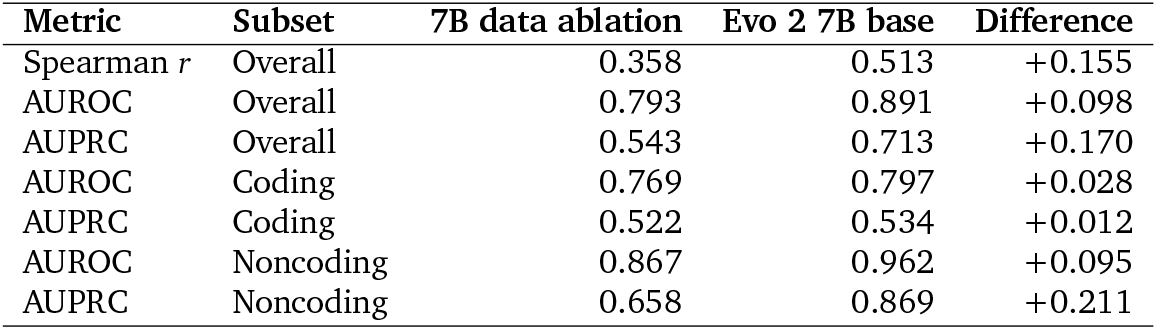
*BRCA1* zero-shot comparison between models.

| Metric | Subset | 7B data ablation | Evo 2 7B base | Difference |
| --- | --- | --- | --- | --- |
| Spearman $r$ | Overall | 0.358 | 0.513 | +0.155 |
| AUROC | Overall | 0.793 | 0.891 | +0.098 |
| AUPRC | Overall | 0.543 | 0.713 | +0.170 |
| AUROC | Coding | 0.769 | 0.797 | +0.028 |
| AUPRC | Coding | 0.522 | 0.534 | +0.012 |
| AUROC | Noncoding | 0.867 | 0.962 | +0.095 |
| AUPRC | Noncoding | 0.658 | 0.869 | +0.211 |

**Table S5.**
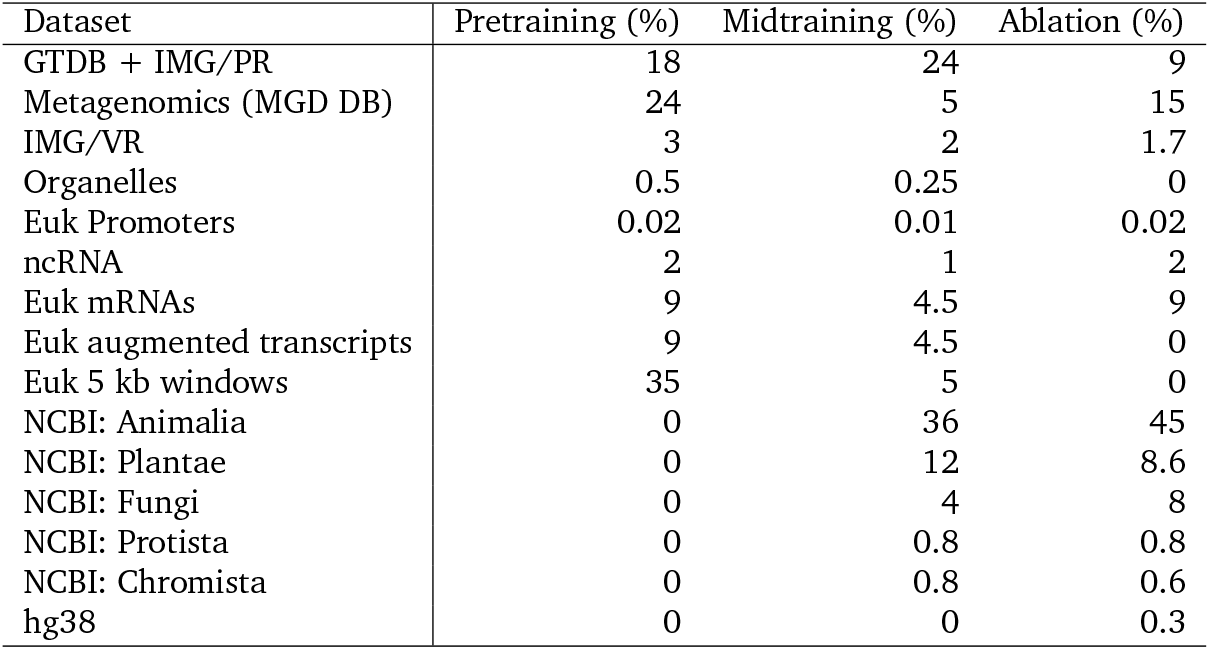
Pretraining, midtraining data composition, and data ablation experiment composition.

| Dataset | Pretraining (%) | Midtraining (%) | Ablation (%) |
| --- | --- | --- | --- |
| GTDB + IMG/PR | 18 | 24 | 9 |
| Metagenomics (MGD DB) | 24 | 5 | 15 |
| IMG/VR | 3 | 2 | 1.7 |
| Organelles | 0.5 | 0.25 | 0 |
| Euk Promoters | 0.02 | 0.01 | 0.02 |
| ncRNA | 2 | 1 | 2 |
| Euk mRNAs | 9 | 4.5 | 9 |
| Euk augmented transcripts | 9 | 4.5 | 0 |
| Euk 5 kb windows | 35 | 5 | 0 |
| NCBI: Animalia | 0 | 36 | 45 |
| NCBI: Plantae | 0 | 12 | 8.6 |
| NCBI: Fungi | 0 | 4 | 8 |
| NCBI: Protista | 0 | 0.8 | 0.8 |
| NCBI: Chromista | 0 | 0.8 | 0.6 |
| hg38 | 0 | 0 | 0.3 |

**Table S6.** BLASTp results for genes from Evo 2 generated mitochondria. The amino acid percent sequence ID is obtained by multiplying the query cover with the sequence identity.

| Annotation | Sequence ID % | Nearest Species |
| --- | --- | --- |
| mt-Co1 | 86.94 | *Muntiacus reevesi* |
| mt-Atp8 | 77.27 | *Myotis albescens* |
| mt-Nd4 | 84.95 | *Semilabeo notabilis* |
| mt-Nd1 | 84.59 | *Phocoena phocoena* |
| mt-Co3 | 77.31 | *Alces alces* |
| MT-ND4L | 86.73 | *Bos grunniens* |
| mt-Nd5 | 78.99 | *Schizothorax argentatus* |
| MT-ND2 | 68.45 | *Rusa unicolor* |
| mt-Co2 | 88.54 | *Ovis aries* |
| mt-Atp6 | 91.59 | *Alcelaphus buselaphus* |
| mt-Nd3 | 87.11 | *Bos indicus* |
| mt-Cytb | 75.08 | *Davidijordania poecilimon* |
| mt-Nd6 | 94.43 | *Neolissochilus hendersoni* |

**Table S7.**
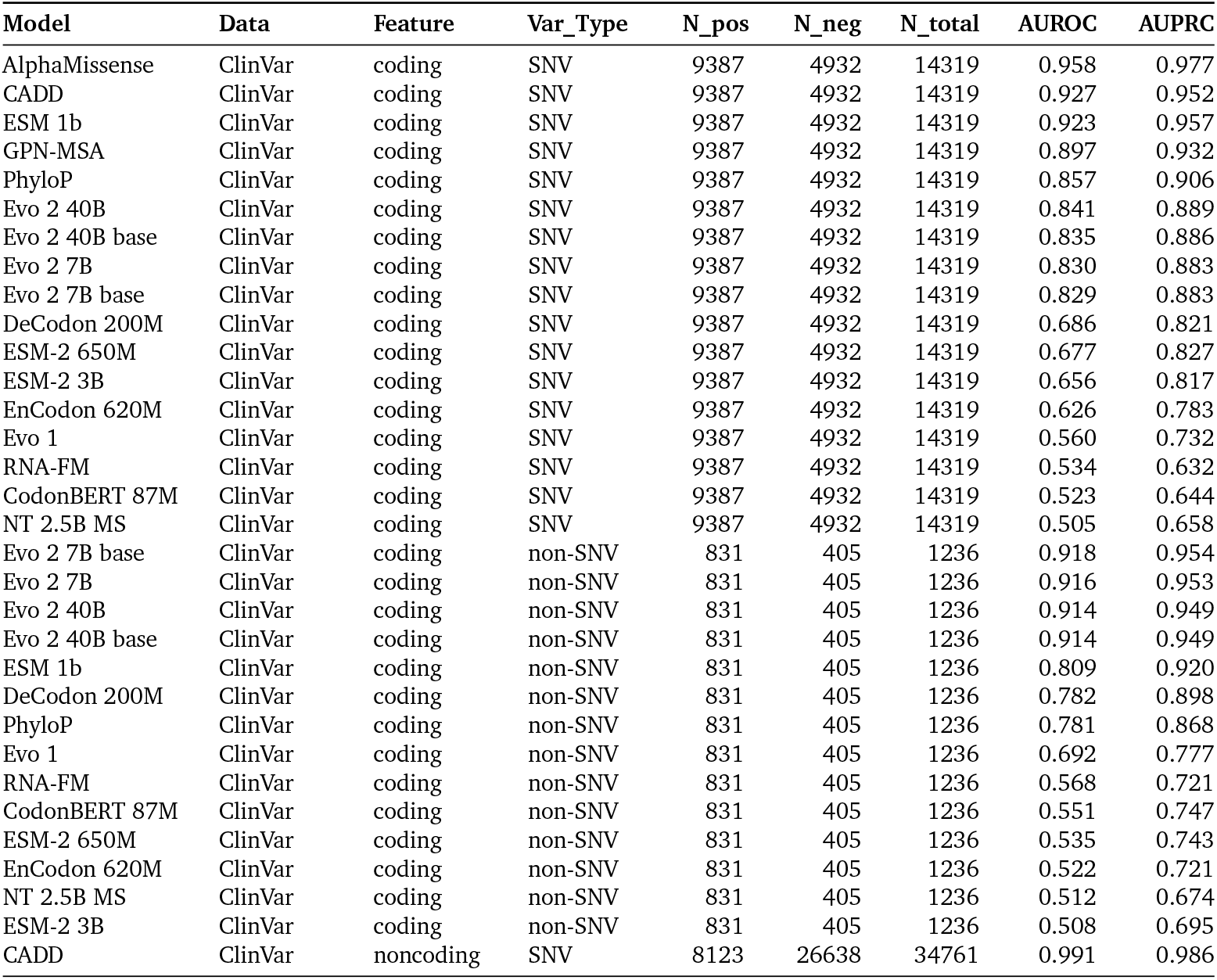
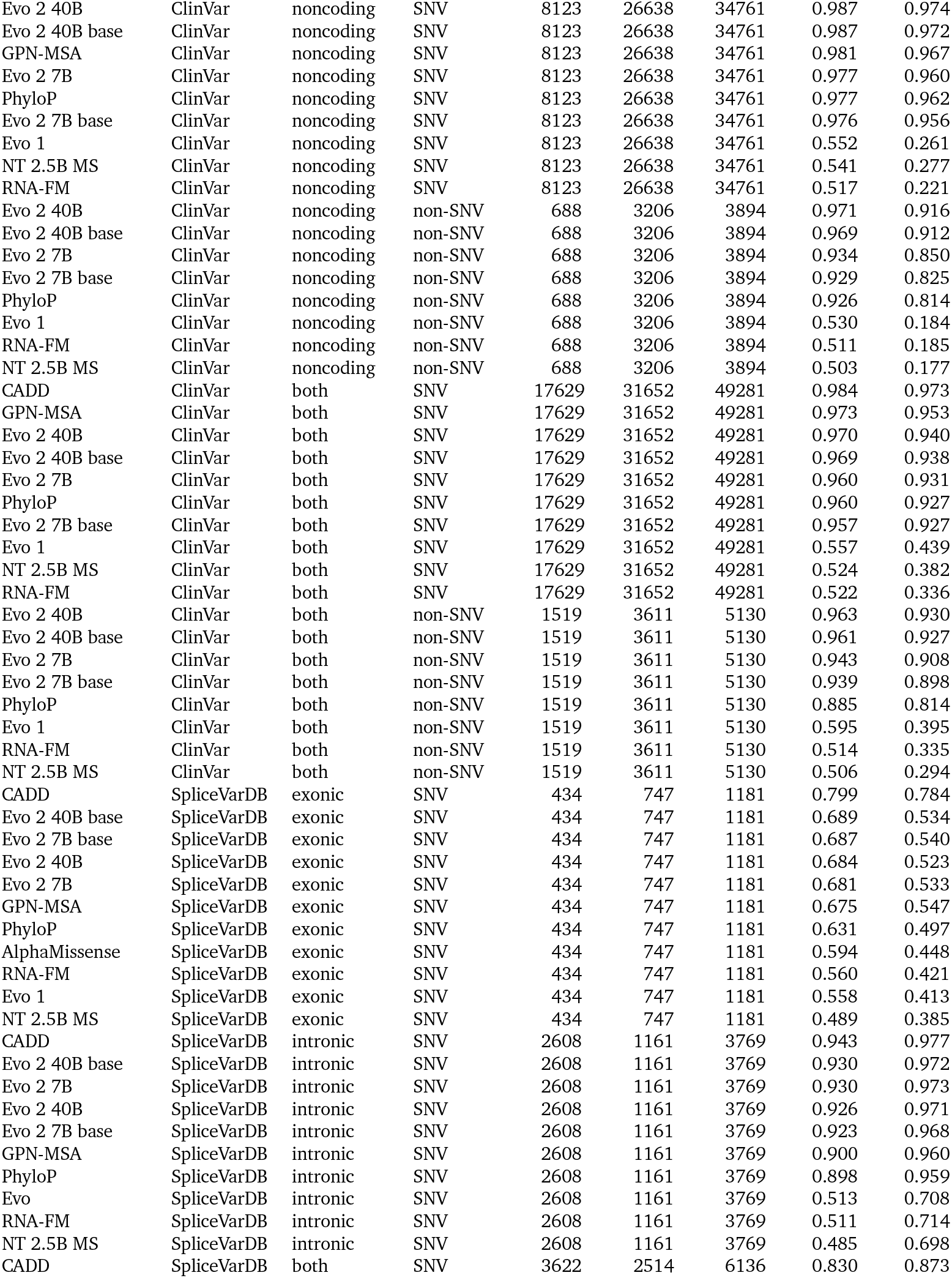
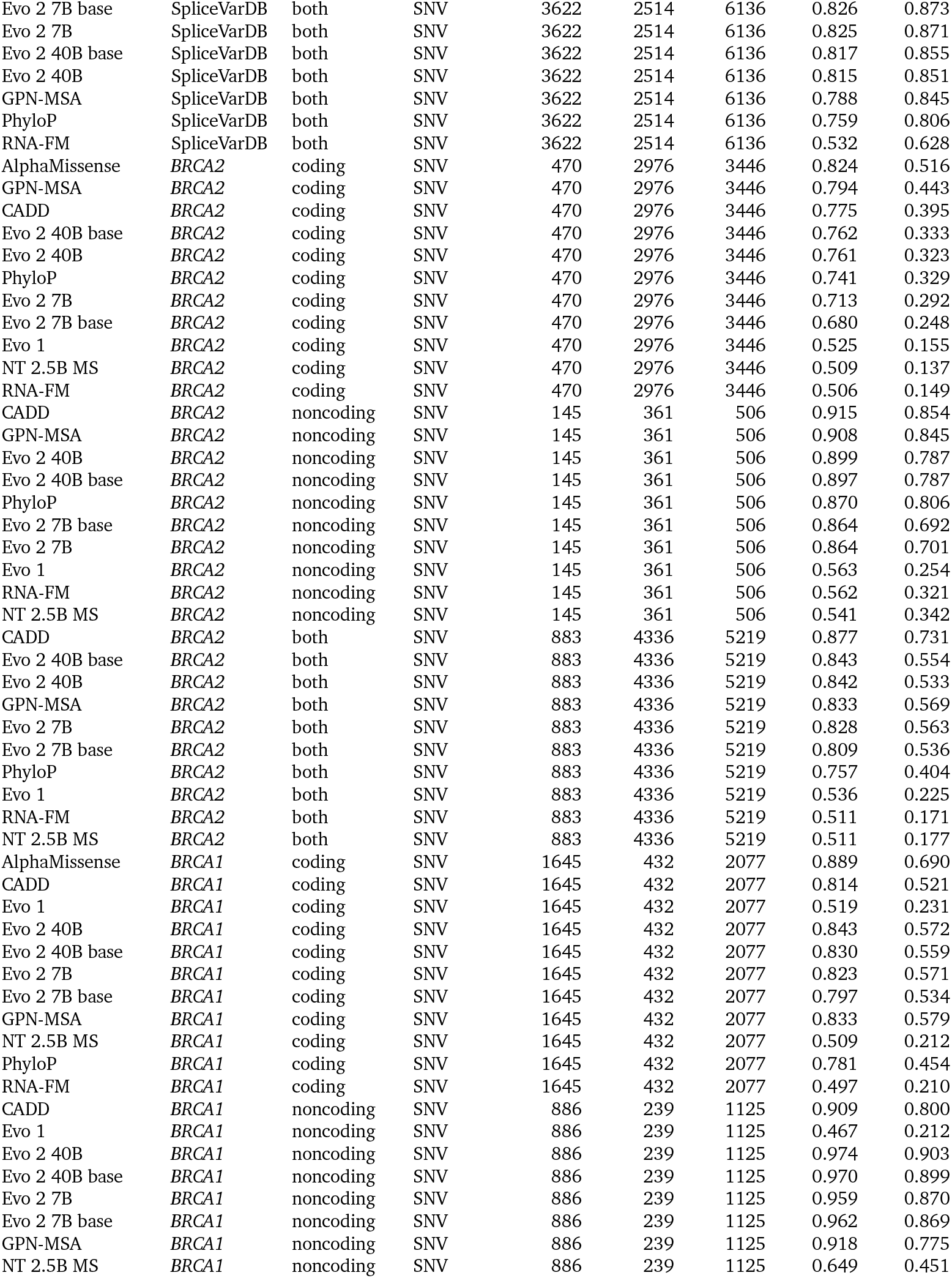
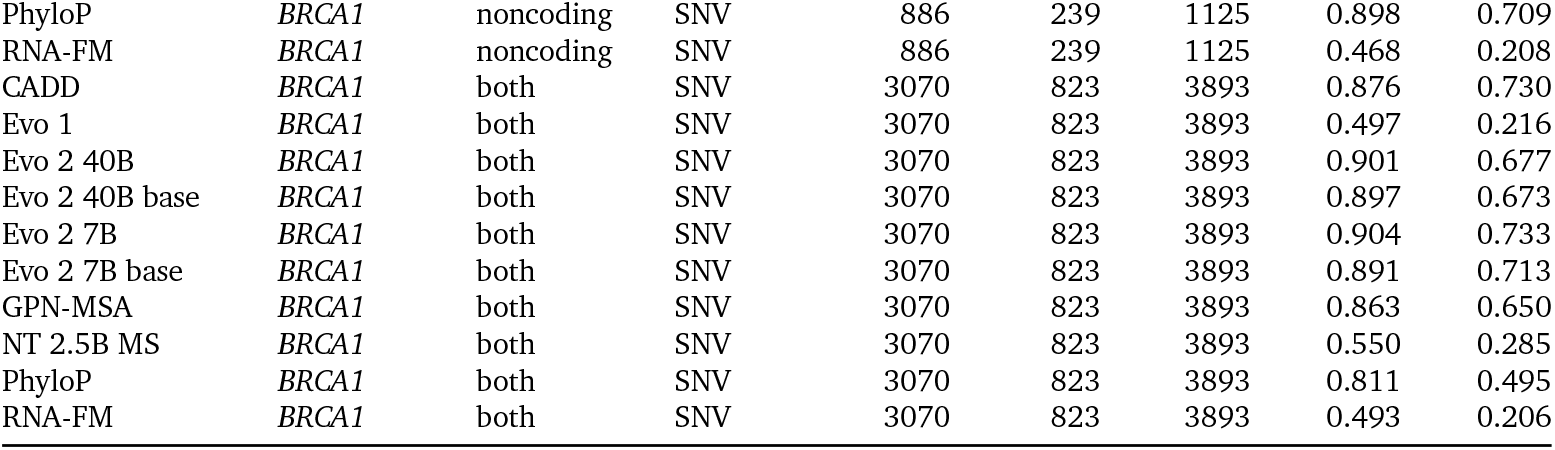
Comparison of zero-shot variant effect predictions across models, variant types (SNV and non-SNV), and associated genomic features (coding vs noncoding, intronic vs exonic). In addition to AUROC and AUPRC performance metrics, counts of variants falling into either positive (N_pos) or negative (N_neg) classes for each evaluation are also included.

| Model | Data | Feature | Var_Type | N_pos | N_neg | N_total | AUROC | AUPRC |
| --- | --- | --- | --- | --- | --- | --- | --- | --- |
| AlphaMissense | ClinVar | coding | SNV | 9387 | 4932 | 14319 | 0.958 | 0.977 |
| CADD | ClinVar | coding | SNV | 9387 | 4932 | 14319 | 0.927 | 0.952 |
| ESM 1b | ClinVar | coding | SNV | 9387 | 4932 | 14319 | 0.923 | 0.957 |
| GPN-MSA | ClinVar | coding | SNV | 9387 | 4932 | 14319 | 0.897 | 0.932 |
| PhyloP | ClinVar | coding | SNV | 9387 | 4932 | 14319 | 0.857 | 0.906 |
| Evo 2 40B | ClinVar | coding | SNV | 9387 | 4932 | 14319 | 0.841 | 0.889 |
| Evo 2 40B base | ClinVar | coding | SNV | 9387 | 4932 | 14319 | 0.835 | 0.886 |
| Evo 2 7B | ClinVar | coding | SNV | 9387 | 4932 | 14319 | 0.830 | 0.883 |
| Evo 2 7B base | ClinVar | coding | SNV | 9387 | 4932 | 14319 | 0.829 | 0.883 |
| DeCodon 200M | ClinVar | coding | SNV | 9387 | 4932 | 14319 | 0.686 | 0.821 |
| ESM-2 650M | ClinVar | coding | SNV | 9387 | 4932 | 14319 | 0.677 | 0.827 |
| ESM-2 3B | ClinVar | coding | SNV | 9387 | 4932 | 14319 | 0.656 | 0.817 |
| EnCodon 620M | ClinVar | coding | SNV | 9387 | 4932 | 14319 | 0.626 | 0.783 |
| Evo 1 | ClinVar | coding | SNV | 9387 | 4932 | 14319 | 0.560 | 0.732 |
| RNA-FM | ClinVar | coding | SNV | 9387 | 4932 | 14319 | 0.534 | 0.632 |
| CodonBERT 87M | ClinVar | coding | SNV | 9387 | 4932 | 14319 | 0.523 | 0.644 |
| NT 2.5B MS | ClinVar | coding | SNV | 9387 | 4932 | 14319 | 0.505 | 0.658 |
| Evo 2 7B base | ClinVar | coding | non-SNV | 831 | 405 | 1236 | 0.918 | 0.954 |
| Evo 2 7B | ClinVar | coding | non-SNV | 831 | 405 | 1236 | 0.916 | 0.953 |
| Evo 2 40B | ClinVar | coding | non-SNV | 831 | 405 | 1236 | 0.914 | 0.949 |
| Evo 2 40B base | ClinVar | coding | non-SNV | 831 | 405 | 1236 | 0.914 | 0.949 |
| ESM 1b | ClinVar | coding | non-SNV | 831 | 405 | 1236 | 0.809 | 0.920 |
| DeCodon 200M | ClinVar | coding | non-SNV | 831 | 405 | 1236 | 0.782 | 0.898 |
| PhyloP | ClinVar | coding | non-SNV | 831 | 405 | 1236 | 0.781 | 0.868 |
| Evo 1 | ClinVar | coding | non-SNV | 831 | 405 | 1236 | 0.692 | 0.777 |
| RNA-FM | ClinVar | coding | non-SNV | 831 | 405 | 1236 | 0.568 | 0.721 |
| CodonBERT 87M | ClinVar | coding | non-SNV | 831 | 405 | 1236 | 0.551 | 0.747 |
| ESM-2 650M | ClinVar | coding | non-SNV | 831 | 405 | 1236 | 0.535 | 0.743 |
| EnCodon 620M | ClinVar | coding | non-SNV | 831 | 405 | 1236 | 0.522 | 0.721 |
| NT 2.5B MS | ClinVar | coding | non-SNV | 831 | 405 | 1236 | 0.512 | 0.674 |
| ESM-2 3B | ClinVar | coding | non-SNV | 831 | 405 | 1236 | 0.508 | 0.695 |
| CADD | ClinVar | noncoding | SNV | 8123 | 26638 | 34761 | 0.991 | 0.986 |
| Evo 2 40B | ClinVar | noncoding | SNV | 8123 | 26638 | 34761 | 0.987 | 0.974 |
| Evo 2 40B base | ClinVar | noncoding | SNV | 8123 | 26638 | 34761 | 0.987 | 0.972 |
| GPN-MSA | ClinVar | noncoding | SNV | 8123 | 26638 | 34761 | 0.981 | 0.967 |
| Evo 2 7B | ClinVar | noncoding | SNV | 8123 | 26638 | 34761 | 0.977 | 0.960 |
| PhyloP | ClinVar | noncoding | SNV | 8123 | 26638 | 34761 | 0.977 | 0.962 |
| Evo 2 7B base | ClinVar | noncoding | SNV | 8123 | 26638 | 34761 | 0.976 | 0.956 |
| Evo 1 | ClinVar | noncoding | SNV | 8123 | 26638 | 34761 | 0.552 | 0.261 |
| NT 2.5B MS | ClinVar | noncoding | SNV | 8123 | 26638 | 34761 | 0.541 | 0.277 |
| RNA-FM | ClinVar | noncoding | SNV | 8123 | 26638 | 34761 | 0.517 | 0.221 |
| Evo 2 40B | ClinVar | noncoding | non-SNV | 688 | 3206 | 3894 | 0.971 | 0.916 |
| Evo 2 40B base | ClinVar | noncoding | non-SNV | 688 | 3206 | 3894 | 0.969 | 0.912 |
| Evo 2 7B | ClinVar | noncoding | non-SNV | 688 | 3206 | 3894 | 0.934 | 0.850 |
| Evo 2 7B base | ClinVar | noncoding | non-SNV | 688 | 3206 | 3894 | 0.929 | 0.825 |
| PhyloP | ClinVar | noncoding | non-SNV | 688 | 3206 | 3894 | 0.926 | 0.814 |
| Evo 1 | ClinVar | noncoding | non-SNV | 688 | 3206 | 3894 | 0.530 | 0.184 |
| RNA-FM | ClinVar | noncoding | non-SNV | 688 | 3206 | 3894 | 0.511 | 0.185 |
| NT 2.5B MS | ClinVar | noncoding | non-SNV | 688 | 3206 | 3894 | 0.503 | 0.177 |
| CADD | ClinVar | both | SNV | 17629 | 31652 | 49281 | 0.984 | 0.973 |
| GPN-MSA | ClinVar | both | SNV | 17629 | 31652 | 49281 | 0.973 | 0.953 |
| Evo 2 40B | ClinVar | both | SNV | 17629 | 31652 | 49281 | 0.970 | 0.940 |
| Evo 2 40B base | ClinVar | both | SNV | 17629 | 31652 | 49281 | 0.969 | 0.938 |
| Evo 2 7B | ClinVar | both | SNV | 17629 | 31652 | 49281 | 0.960 | 0.931 |
| PhyloP | ClinVar | both | SNV | 17629 | 31652 | 49281 | 0.960 | 0.927 |
| Evo 2 7B base | ClinVar | both | SNV | 17629 | 31652 | 49281 | 0.957 | 0.927 |
| Evo 1 | ClinVar | both | SNV | 17629 | 31652 | 49281 | 0.557 | 0.439 |
| NT 2.5B MS | ClinVar | both | SNV | 17629 | 31652 | 49281 | 0.524 | 0.382 |
| RNA-FM | ClinVar | both | SNV | 17629 | 31652 | 49281 | 0.522 | 0.336 |
| Evo 2 40B | ClinVar | both | non-SNV | 1519 | 3611 | 5130 | 0.963 | 0.930 |
| Evo 2 40B base | ClinVar | both | non-SNV | 1519 | 3611 | 5130 | 0.961 | 0.927 |
| Evo 2 7B | ClinVar | both | non-SNV | 1519 | 3611 | 5130 | 0.943 | 0.908 |
| Evo 2 7B base | ClinVar | both | non-SNV | 1519 | 3611 | 5130 | 0.939 | 0.898 |
| PhyloP | ClinVar | both | non-SNV | 1519 | 3611 | 5130 | 0.885 | 0.814 |
| Evo 1 | ClinVar | both | non-SNV | 1519 | 3611 | 5130 | 0.595 | 0.395 |
| RNA-FM | ClinVar | both | non-SNV | 1519 | 3611 | 5130 | 0.514 | 0.335 |
| NT 2.5B MS | ClinVar | both | non-SNV | 1519 | 3611 | 5130 | 0.506 | 0.294 |
| CADD | SpliceVarDB | exonic | SNV | 434 | 747 | 1181 | 0.799 | 0.784 |
| Evo 2 40B base | SpliceVarDB | exonic | SNV | 434 | 747 | 1181 | 0.689 | 0.534 |
| Evo 2 7B base | SpliceVarDB | exonic | SNV | 434 | 747 | 1181 | 0.687 | 0.540 |
| Evo 2 40B | SpliceVarDB | exonic | SNV | 434 | 747 | 1181 | 0.684 | 0.523 |
| Evo 2 7B | SpliceVarDB | exonic | SNV | 434 | 747 | 1181 | 0.681 | 0.533 |
| GPN-MSA | SpliceVarDB | exonic | SNV | 434 | 747 | 1181 | 0.675 | 0.547 |
| PhyloP | SpliceVarDB | exonic | SNV | 434 | 747 | 1181 | 0.631 | 0.497 |
| AlphaMissense | SpliceVarDB | exonic | SNV | 434 | 747 | 1181 | 0.594 | 0.448 |
| RNA-FM | SpliceVarDB | exonic | SNV | 434 | 747 | 1181 | 0.560 | 0.421 |
| Evo 1 | SpliceVarDB | exonic | SNV | 434 | 747 | 1181 | 0.558 | 0.413 |
| NT 2.5B MS | SpliceVarDB | exonic | SNV | 434 | 747 | 1181 | 0.489 | 0.385 |
| CADD | SpliceVarDB | intronic | SNV | 2608 | 1161 | 3769 | 0.943 | 0.977 |
| Evo 2 40B base | SpliceVarDB | intronic | SNV | 2608 | 1161 | 3769 | 0.930 | 0.972 |
| Evo 2 7B | SpliceVarDB | intronic | SNV | 2608 | 1161 | 3769 | 0.930 | 0.973 |
| Evo 2 40B | SpliceVarDB | intronic | SNV | 2608 | 1161 | 3769 | 0.926 | 0.971 |
| Evo 2 7B base | SpliceVarDB | intronic | SNV | 2608 | 1161 | 3769 | 0.923 | 0.968 |
| GPN-MSA | SpliceVarDB | intronic | SNV | 2608 | 1161 | 3769 | 0.900 | 0.960 |
| PhyloP | SpliceVarDB | intronic | SNV | 2608 | 1161 | 3769 | 0.898 | 0.959 |
| Evo | SpliceVarDB | intronic | SNV | 2608 | 1161 | 3769 | 0.513 | 0.708 |
| RNA-FM | SpliceVarDB | intronic | SNV | 2608 | 1161 | 3769 | 0.511 | 0.714 |
| NT 2.5B MS | SpliceVarDB | intronic | SNV | 2608 | 1161 | 3769 | 0.485 | 0.698 |
| CADD | SpliceVarDB | both | SNV | 3622 | 2514 | 6136 | 0.830 | 0.873 |
| Evo 2 7B base | SpliceVarDB | both | SNV | 3622 | 2514 | 6136 | 0.826 | 0.873 |
| Evo 2 7B | SpliceVarDB | both | SNV | 3622 | 2514 | 6136 | 0.825 | 0.871 |
| Evo 2 40B base | SpliceVarDB | both | SNV | 3622 | 2514 | 6136 | 0.817 | 0.855 |
| Evo 2 40B | SpliceVarDB | both | SNV | 3622 | 2514 | 6136 | 0.815 | 0.851 |
| GPN-MSA | SpliceVarDB | both | SNV | 3622 | 2514 | 6136 | 0.788 | 0.845 |
| PhyloP | SpliceVarDB | both | SNV | 3622 | 2514 | 6136 | 0.759 | 0.806 |
| RNA-FM | SpliceVarDB | both | SNV | 3622 | 2514 | 6136 | 0.532 | 0.628 |
| AlphaMissense | BRCA2 | coding | SNV | 470 | 2976 | 3446 | 0.824 | 0.516 |
| GPN-MSA | BRCA2 | coding | SNV | 470 | 2976 | 3446 | 0.794 | 0.443 |
| CADD | BRCA2 | coding | SNV | 470 | 2976 | 3446 | 0.775 | 0.395 |
| Evo 2 40B base | BRCA2 | coding | SNV | 470 | 2976 | 3446 | 0.762 | 0.333 |
| Evo 2 40B | BRCA2 | coding | SNV | 470 | 2976 | 3446 | 0.761 | 0.323 |
| PhyloP | BRCA2 | coding | SNV | 470 | 2976 | 3446 | 0.741 | 0.329 |
| Evo 2 7B | BRCA2 | coding | SNV | 470 | 2976 | 3446 | 0.713 | 0.292 |
| Evo 2 7B base | BRCA2 | coding | SNV | 470 | 2976 | 3446 | 0.680 | 0.248 |
| Evo 1 | BRCA2 | coding | SNV | 470 | 2976 | 3446 | 0.525 | 0.155 |
| NT 2.5B MS | BRCA2 | coding | SNV | 470 | 2976 | 3446 | 0.509 | 0.137 |
| RNA-FM | BRCA2 | coding | SNV | 470 | 2976 | 3446 | 0.506 | 0.149 |
| CADD | BRCA2 | noncoding | SNV | 145 | 361 | 506 | 0.915 | 0.854 |
| GPN-MSA | BRCA2 | noncoding | SNV | 145 | 361 | 506 | 0.908 | 0.845 |
| Evo 2 40B | BRCA2 | noncoding | SNV | 145 | 361 | 506 | 0.899 | 0.787 |
| Evo 2 40B base | BRCA2 | noncoding | SNV | 145 | 361 | 506 | 0.897 | 0.787 |
| PhyloP | BRCA2 | noncoding | SNV | 145 | 361 | 506 | 0.870 | 0.806 |
| Evo 2 7B base | BRCA2 | noncoding | SNV | 145 | 361 | 506 | 0.864 | 0.692 |
| Evo 2 7B | BRCA2 | noncoding | SNV | 145 | 361 | 506 | 0.864 | 0.701 |
| Evo 1 | BRCA2 | noncoding | SNV | 145 | 361 | 506 | 0.563 | 0.254 |
| RNA-FM | BRCA2 | noncoding | SNV | 145 | 361 | 506 | 0.562 | 0.321 |
| NT 2.5B MS | BRCA2 | noncoding | SNV | 145 | 361 | 506 | 0.541 | 0.342 |
| CADD | BRCA2 | both | SNV | 883 | 4336 | 5219 | 0.877 | 0.731 |
| Evo 2 40B base | BRCA2 | both | SNV | 883 | 4336 | 5219 | 0.843 | 0.554 |
| Evo 2 40B | BRCA2 | both | SNV | 883 | 4336 | 5219 | 0.842 | 0.533 |
| GPN-MSA | BRCA2 | both | SNV | 883 | 4336 | 5219 | 0.833 | 0.569 |
| Evo 2 7B | BRCA2 | both | SNV | 883 | 4336 | 5219 | 0.828 | 0.563 |
| Evo 2 7B base | BRCA2 | both | SNV | 883 | 4336 | 5219 | 0.809 | 0.536 |
| PhyloP | BRCA2 | both | SNV | 883 | 4336 | 5219 | 0.757 | 0.404 |
| Evo 1 | BRCA2 | both | SNV | 883 | 4336 | 5219 | 0.536 | 0.225 |
| RNA-FM | BRCA2 | both | SNV | 883 | 4336 | 5219 | 0.511 | 0.171 |
| NT 2.5B MS | BRCA2 | both | SNV | 883 | 4336 | 5219 | 0.511 | 0.177 |
| AlphaMissense | BRCA1 | coding | SNV | 1645 | 432 | 2077 | 0.889 | 0.690 |
| CADD | BRCA1 | coding | SNV | 1645 | 432 | 2077 | 0.814 | 0.521 |
| Evo 1 | BRCA1 | coding | SNV | 1645 | 432 | 2077 | 0.519 | 0.231 |
| Evo 2 40B | BRCA1 | coding | SNV | 1645 | 432 | 2077 | 0.843 | 0.572 |
| Evo 2 40B base | BRCA1 | coding | SNV | 1645 | 432 | 2077 | 0.830 | 0.559 |
| Evo 2 7B | BRCA1 | coding | SNV | 1645 | 432 | 2077 | 0.823 | 0.571 |
| Evo 2 7B base | BRCA1 | coding | SNV | 1645 | 432 | 2077 | 0.797 | 0.534 |
| GPN-MSA | BRCA1 | coding | SNV | 1645 | 432 | 2077 | 0.833 | 0.579 |
| NT 2.5B MS | BRCA1 | coding | SNV | 1645 | 432 | 2077 | 0.509 | 0.212 |
| PhyloP | BRCA1 | coding | SNV | 1645 | 432 | 2077 | 0.781 | 0.454 |
| RNA-FM | BRCA1 | coding | SNV | 1645 | 432 | 2077 | 0.497 | 0.210 |
| CADD | BRCA1 | noncoding | SNV | 886 | 239 | 1125 | 0.909 | 0.800 |
| Evo 1 | BRCA1 | noncoding | SNV | 886 | 239 | 1125 | 0.467 | 0.212 |
| Evo 2 40B | BRCA1 | noncoding | SNV | 886 | 239 | 1125 | 0.974 | 0.903 |
| Evo 2 40B base | BRCA1 | noncoding | SNV | 886 | 239 | 1125 | 0.970 | 0.899 |
| Evo 2 7B | BRCA1 | noncoding | SNV | 886 | 239 | 1125 | 0.959 | 0.870 |
| Evo 2 7B base | BRCA1 | noncoding | SNV | 886 | 239 | 1125 | 0.962 | 0.869 |
| GPN-MSA | BRCA1 | noncoding | SNV | 886 | 239 | 1125 | 0.918 | 0.775 |
| NT 2.5B MS | BRCA1 | noncoding | SNV | 886 | 239 | 1125 | 0.649 | 0.451 |
| PhyloP | *BRCA1* | noncoding | SNV | 886 | 239 | 1125 | 0.898 | 0.709 |
| RNA-FM | *BRCA1* | noncoding | SNV | 886 | 239 | 1125 | 0.468 | 0.208 |
| CADD | *BRCA1* | both | SNV | 3070 | 823 | 3893 | 0.876 | 0.730 |
| Evo 1 | *BRCA1* | both | SNV | 3070 | 823 | 3893 | 0.497 | 0.216 |
| Evo 2 40B | *BRCA1* | both | SNV | 3070 | 823 | 3893 | 0.901 | 0.677 |
| Evo 2 40B base | *BRCA1* | both | SNV | 3070 | 823 | 3893 | 0.897 | 0.673 |
| Evo 2 7B | *BRCA1* | both | SNV | 3070 | 823 | 3893 | 0.904 | 0.733 |
| Evo 2 7B base | *BRCA1* | both | SNV | 3070 | 823 | 3893 | 0.891 | 0.713 |
| GPN-MSA | *BRCA1* | both | SNV | 3070 | 823 | 3893 | 0.863 | 0.650 |
| NT 2.5B MS | *BRCA1* | both | SNV | 3070 | 823 | 3893 | 0.550 | 0.285 |
| PhyloP | *BRCA1* | both | SNV | 3070 | 823 | 3893 | 0.811 | 0.495 |
| RNA-FM | *BRCA1* | both | SNV | 3070 | 823 | 3893 | 0.493 | 0.206 |

